# Genomics, Molecular and Evolutionary Perspective of NAC Transcription Factors

**DOI:** 10.1101/608885

**Authors:** Tapan Kumar Mohanta, Dhananjay Yadav, Adil Khan, Abeer Hashem, Baby Tabassum, Abdul Latif Khan, Eslayed Fathi Abda_Allah, Ahmed Al-Harrasi

## Abstract

NAC (NAM, ATAF1,2, and CUC2) transcription factors are one of the largest transcription factor families found in plants and are involved in diverse developmental and signalling events. Despite the availability of comprehensive genomic information from diverse plant species, the basic genomic, biochemical, and evolutionary details of NAC TFs have not been established. Therefore, NAC TFs family proteins from 160 plant species were analyzed in the current study. The analysis, among other things, identified the first algal NAC TF in the Charophyte, *Klebsormidium flaccidum.* Furthermore, our analysis revealed that NAC TFs are membrane bound and contain monopartite, bipartite, and multipartite nuclear localization signals. NAC TFs were also found to encode a novel chimeric protein domain and are part of a complex interactome network. Synonymous codon usage is absent in NAC TFs and it appears that they have evolved from orthologous ancestors and undergone significant duplication events to give rise to paralogous NAC TFs.

## Introduction

Next-generation sequencing (NGS) has fostered the sequencing of many plant genomes. The availability of so many genomes has allowed researchers to readily identify genes, examine genetic diversity within a species, and gain insight into the evolution of genes and gene families. Gene expression is regulated in part by different families of proteins known as transcription factors (TFs). TFs are involved in inducing the transcription of DNA into RNA. They include numerous and diverse proteins, all of which contain one or more DNA-binding motifs. The DNA-binding domain enables them to bind to the promoter or repressor sequence of DNA that is present either at the upstream, downstream, or within an intron region of a coding gene. Some TFs bind to a DNA promoter region located near the transcription start site of a gene and help to form the transcription initiation complex. Other TFs bind to regulatory enhancer sequences and stimulate or repress transcription of the related genes. Regulating transcription is of paramount importance to controlling gene expression and TFs enable the expression of an individual gene in a unique manner, such as during different stages of development or in response to biotic or abiotic stress. TFs act as a molecular switch for temporal and spatial gene regulation. A considerable portion of a genome consists of genes encoding transcription factors. For example, there are at least 52 TF families in *Arabidopsis thaliana*, and the NAC (no apical meristem (NAM) TF family is one of them.

NAC TFs are characterised by the presence of a conserved N-terminal NAC domain comprising approximately 150 amino acids and a diversified C-terminal end. The DNA binding NAC domain is divided into five sub-domains designated A-E. Sub-domain A is apparently involved in the formation of functional dimers, while sub-domains B and E appear to be responsible for the functional divergence of NAC genes ^1–4^. The dimeric architecture of NAC proteins can remain stable even at a concentration of 5M NaCl ^4^. The dimerization is established by Leu14-Thr23, and Glu26-Tyr31 amino acid residues. The dimeric form is responsible for the functional unit of stress-responsive SNAC1 and can modulate DNA-binding specificity. Sub-domains C and D contain positively charged amino acids that bind to DNA. The crystal structure of the SNAC1 TF revealed the presence of a central semi-β-barrel formed from seven twisted anti-parallel β-strands with three α-helices ^4^. The NAC domain is most responsible for DNA binding activity that lies between amino acids Val119-Ser183, Lys123-Lys126, with Lys79, Arg85, and Arg88 reside within different strands of β-sheets ^2,5,6^. The remaining portion of the NAC domain contains a loop region composed of the amino acids, Gly144-Gly149 and Lys180-Asn183, which are very flexible in nature ^4^. The loop region of SNAC1 is quite long and different from the loop region of ANAC, an abscisic-acid-responsive NAC, and could underlie the basis for different biological functions. NAC TFs possesses mono or bipartite nuclear localization signals which contain a Lys residue in sub-domain D ^1, 6–8^. In addition, NAC proteins, as part of a mechanism of self-regulation, also modulate the expression of several other proteins ^6,9^. The D subunit of a few NAC TFs contain a hydrophobic negative regulatory domain (NRD), comprised of L-V-F-Y amino acids, which is involved in suppressing transcriptional activity ^10^. For example, the NRD domain can suppress the transcriptional activity of Dof, WRKY, and APETALA 2/dehydration responsive elements (AP2/DRE) TFs ^10^.

Studies indicate that the diverse C-terminal domain contains a transcription regulatory region (TRR) which has several group-specific motifs that can activate or repress transcription activity ^11–14^. The diverse C-terminal region imparts differences in the function of individual NAC proteins by regulating the interaction of NAC TFs with diverse target proteins. Although the C-terminal region of NAC TFs is diverse, it also contains group-specific conserved motifs ^15^. Although the diverse aspects of NAC TFs have been studied, most were conducted within individual plant species. A detailed comparative study of the genomic, molecular biology, and evolution of NAC TFs has not been conducted. Therefore, a comprehensive analysis of NAC TFs is presented in the current study.

## Results and discussion

### NAC transcription factors exhibit diverse genomic and biochemical features

Advancements in genome sequencing technology have enabled the discovery of the genomic details of hundreds of plant species. The availability this genome sequence data allowed us to characterize the genomic details of NAC TFs in diverse plant species. The presence of NAC TFs in 160 species (18774 NAC sequences) was identified and served as the basis of the conducted analyses. Comparisons of NAC sequences revealed that *Brassica napus* has the highest number (410) of NAC TFs, while the pteridophyte plant, *Marchantia polymorpha*, was found to contain the lowest number (9) (Table 1). On average, monocot plants contain a higher (141.20) number of NAC TFs relative to dicot plants (125.56). Except for *Hordeum vulgare* (76), *Saccharum officinarum* (44), and *Zostera marina* (62) all other monocot species possess more than one hundred NAC TFs each (Table 1). Lower eukaryotic plants, bryophytes and pteridophytes also possess NAC TFs. In addition, the algal species, *Klebsormidium flaccidum*, also contains NAC TFs and this finding represents the first report of NAC TFs in algae (Table 1). A NAC TF in *Trifolium pratense* (Tp57577_TGAC_v2_mRNA14116) was found to be the largest NAC TF, comprising 3101 amino acids, while a NAC TF in *Fragaria x ananassa* (FANhyb_icon00034378_a.1.g00001.1) was found to be the smallest NAC TF, comprising only 25 amino acids. Although it only contains a 25 amino acid sequence, it still encodes a NAC domain. Typically, NAC TFs contain a single NAC domain located near the N-terminal region of the protein. The current analysis, however, also identified NAC TFs with two NAC domains. At least 77 of the 160 studied species were found to contain two NAC domains (Table 1).

**Table 1.**
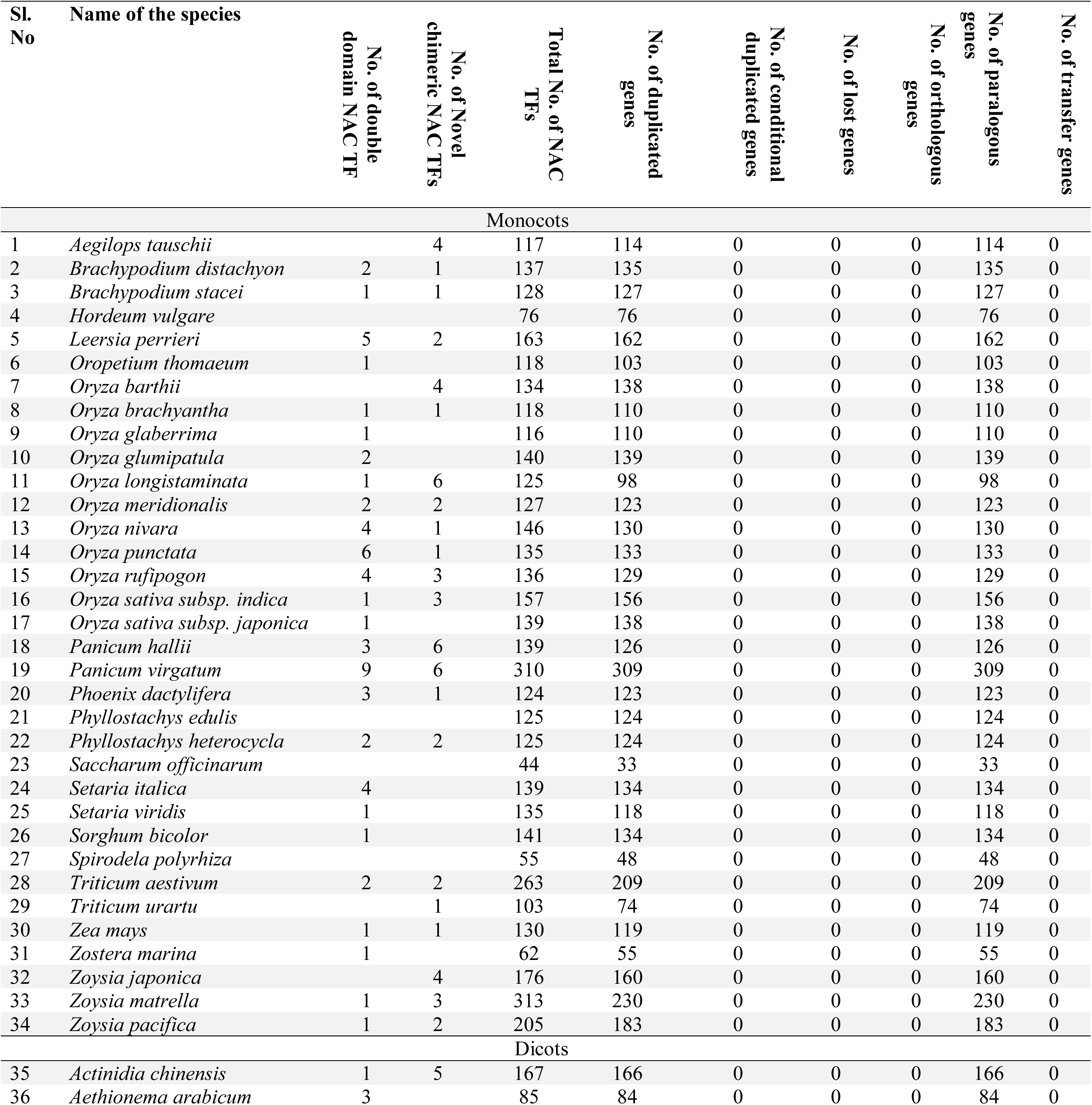
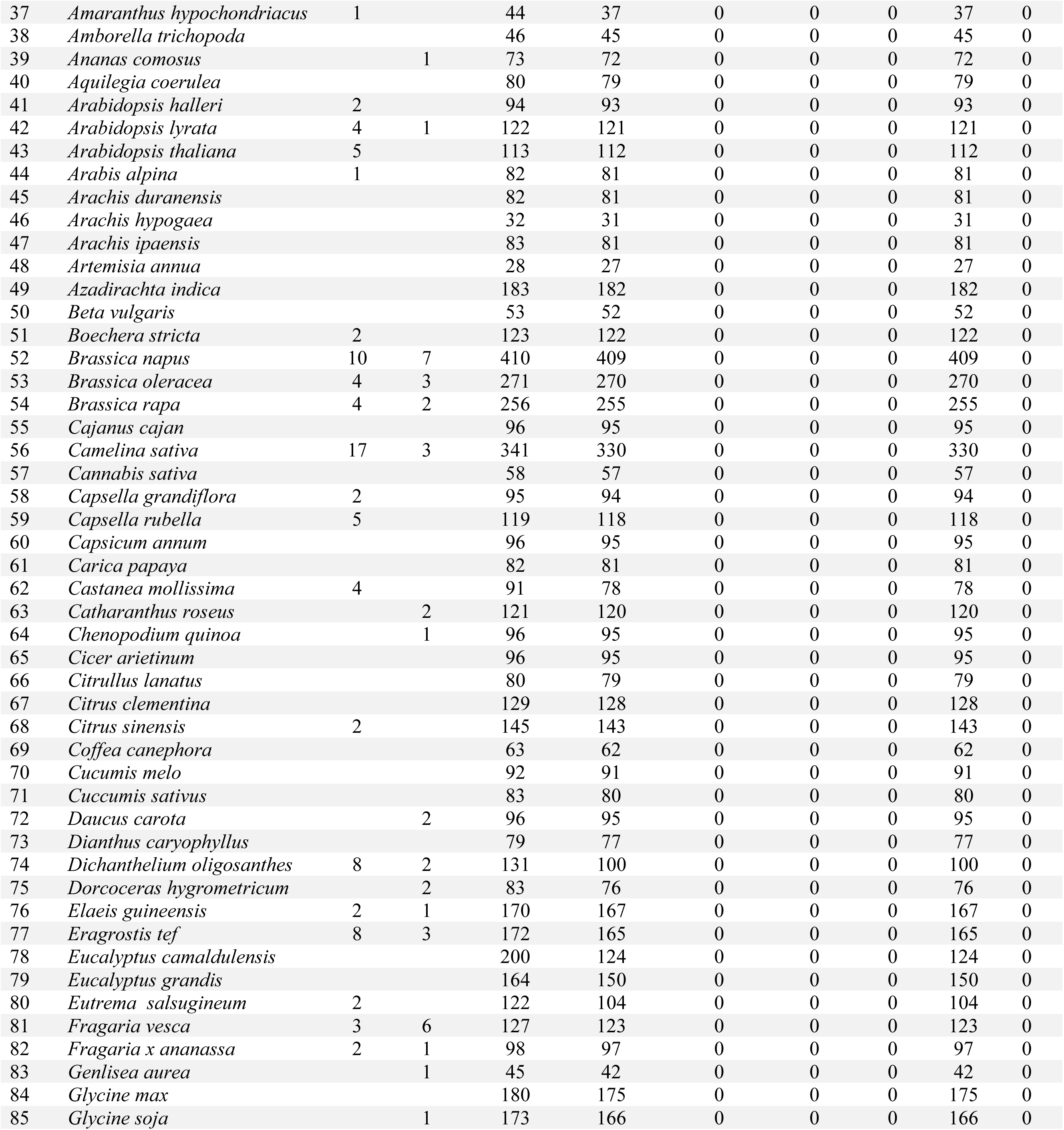
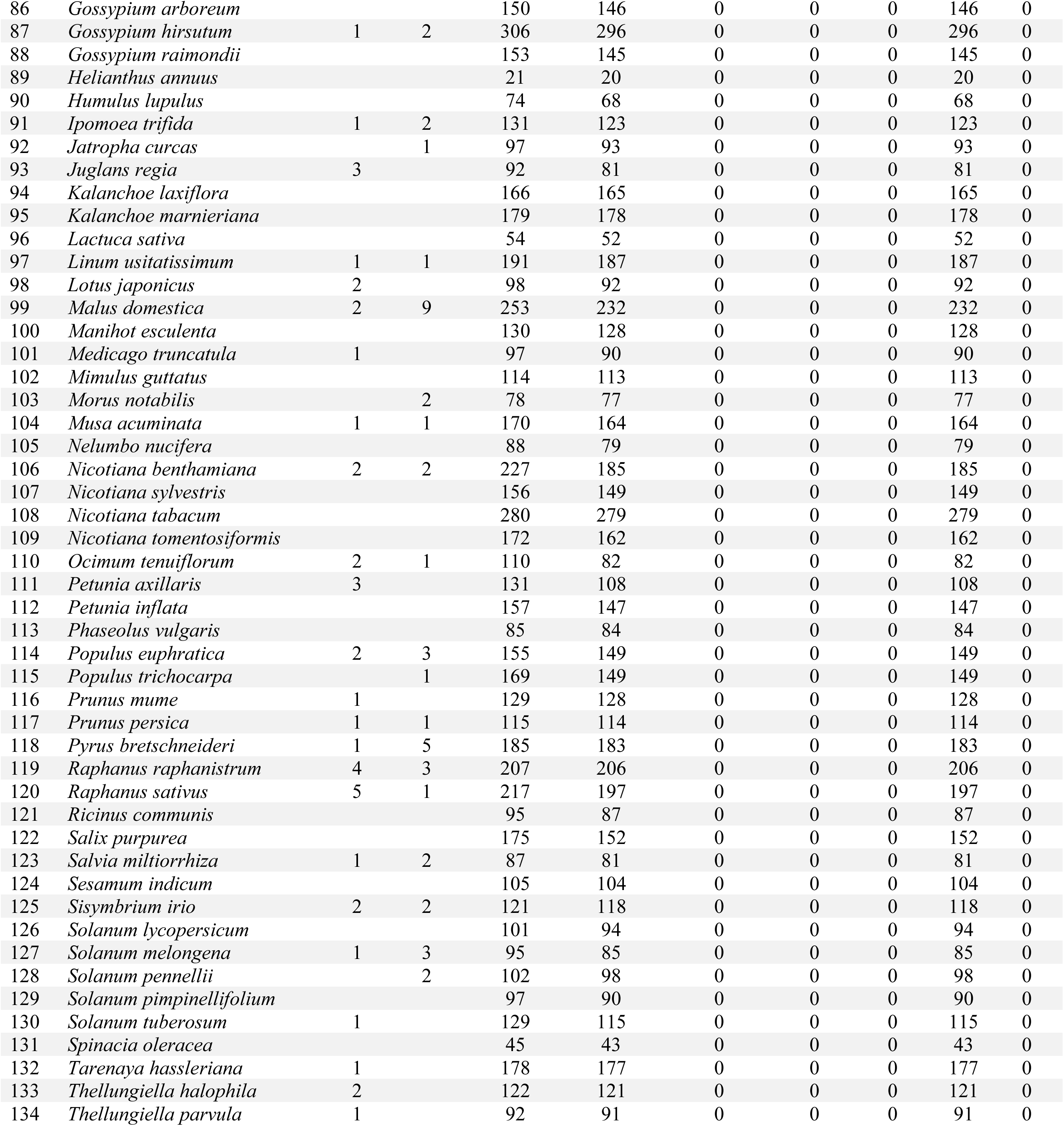
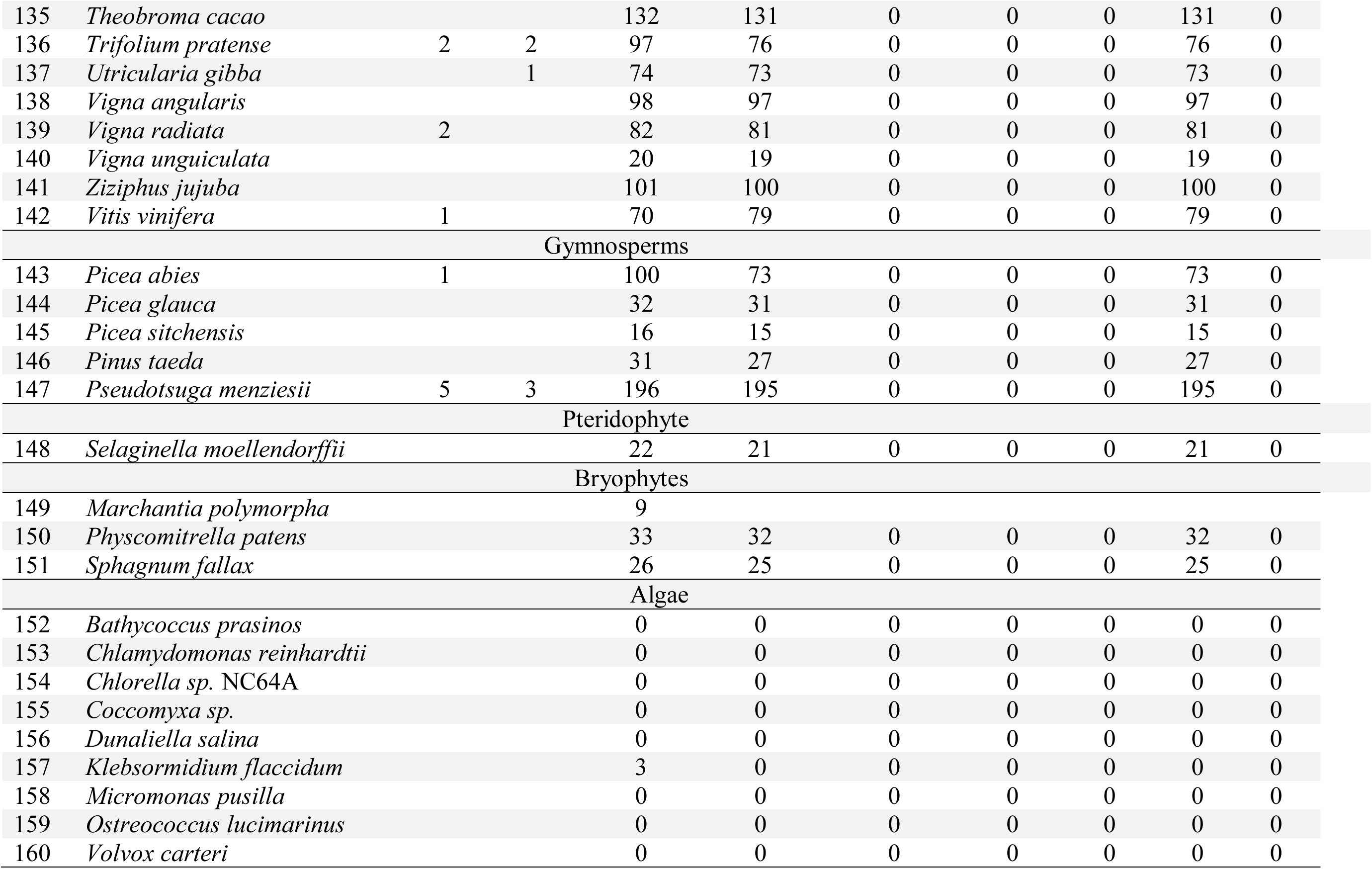
Genomic details of NAC TFs of plants

Multiple sequence alignment revealed the presence of a conserved consensus sequence at the N-terminus. The major conserved consensus sequences are P-G-F-R-F-H-P-T-D-D/E-L-I/V, Y-L-x_2_-K, D-L-x-K-x_2_-P-W-x-L-P, E-W-Y-F-F, G-Y-W-K-A/T-T-G-x-D-x _1-2_-I/V, G-x-K-K-x-L-V-F-Y, and T-x-W-x-M-H-E-Y. Among these consensus sequences, D-D/E-L-I/V, E-W-Y-F-F, G-Y-W-K, and M-H-E-Y are the conserved motifs most observed. The D-D/E-L motif is a characteristic feature of the calcium-binding motifs present in the EF-hand of calcium-dependent protein kinases and the presence of this motif in NAC TFs indicates that they have the potential to regulate Ca^2+^ signalling events in cells ^16^. The D-D-E/E motif is located in the β’ sheet whereas the Y-L-x_2_-K motif is in the α1a/b chain. Except for G-F-R-F-H-P-T-D-D/E-L-I/V, the conserved consensus sequences contain the positively charged amino acids Lys (L) and Arg (K) that can bind to negatively charged DNA. Welner et al. (2012) published the crystal structure of ANAC019 and reported that Y^94^-W-K-A-T-G-T-D in β3, I^11^-K-K-A-L-V-F-Y of β4, K^123^-A-P-K-G-T-K-T-N-W in the loop between β4 and β5, and I^133^-M-H-E-Y-R of β5 and Y^160^-K-K-Q at the C-terminal end are located close to the bound DNA and are associated with DNA binding activity^17^. They reported that Y^94^-W-K-A-T-G-T-D is responsible for the specific recognition of DNA and binds at the major groove within DNA, whereas I^11^-K-K-A-L-V-F-Y, K^123^-A-P-K-G-T-K-T-N-W, I^133^-M-H-E-Y-R, and Y^160^-K-K-Q bind to the backbone of the DNA molecule and provide affinity for DNA binding activity ^17^. In the present analysis of 160 plant species, the identification of the conserved consensus sequences G-Y-W-K-A/T-T-G-x-D-x_1-2_-I/V, G-x-K-K-x-L-V-F-Y, and T-x-W-x-M-H-E-Y is in agreement with Welner et al (2012); suggesting that NAC TFs contain conserved consensus sequences for specific DNA recognition and increasing the affinity for DNA binding.

Hao et al., (2010) reported that the D subunit of NAC TFs contain a hydrophobic L-V-F-Y amino acid motif that suppresses WRKY, Dof, and APETALA2 transcriptional regulators. This suggests that NAC TFs may also function as a negative regulator of transcription ^10^. Our sequence alignment, however, revealed that NAC TF family proteins in many different and diverse plant species possess a conserved hydrophobic L-V-F-Y motif. As reported by Hao et al., (2010), all of the NAC TFs have the potential to suppress the transcriptional activity of WRKY, Dof, and APETALA 2/dehydration responsive element TFs; however, it is highly unlikely that an organism could regulate transcriptional events in a specific and sustained manner if NAC will conduct transcriptional repression.

The molecular weight of NAC TFs ranged from 346.46 kilodaltons (kDa) (*Trifolium pratense*_Tp57577_TGAC_v2) to 2.94 kDa (*Fragaria* x *ananassa*_FANhyb_icon00034378_a.1.g00001.1) (Figure 1). Among the studied NAC TFs, only 10 NAC proteins have a molecular weight (MW) more than 200 kDa and approximately 99 are between 100 to 200 kDa. The MW of the majority of the NAC proteins range between 40 to 55 kDa (Figure 1). The Isoelectric point (pI) of the NAC proteins ranged from 11.47 (Brast01G304500.1.p, (*Brachypodium stacei*) to 3.60 (ObartAA03S_FGP19036, *Oryza barthii*). The majority of the NAC TFs fell within a pI rage of 5-8 (Figure 2). Among the 18774 analysed NAC TFs, the pI of 99 of them were ≥ 10. Approximately 69.28% of the NAC TFs had a pI that was in an acidic range, whereas the remaining 30.72% had a pI within in a basic range. A protein with a pH below the pI carries a net positive charge, whereas a protein with a pH above the pI carries a net negative charge. The pI of a protein determines its transport, solubility, and sub-cellular localization ^18–20^. Biomembranes, such as those surrounding the nucleus, are negatively charged; as a result, positively charged (acidic pI) NAC TFs are readily attracted to the nuclear membrane and subsequently transported into the nucleus to function in transcriptional regulation. There are, however, approximately 30.72% NAC TFs that possess a basic Pi; suggesting that they are localized in the cytosol or plasma membrane of the cell. The major role of TFs is to bind to specific DNA sequences to regulate transcription. The majority of proteins have either an acidic or basic pI and those with a neutral pI close to 7.4 are few because proteins tend to be insoluble, unreactive, and unstable at a pH close to its pI. This is the main reason why among the 18774 NAC TFs analysed, only two (XP_010925972.1, *Elaeis guineensis*; Lus10008200, *Linum usitatissimum*) had a pI 7.4. The existence of NAC proteins with a pI above 10 led us to speculate whether these TFs function while attached to a transmembrane domain. Therefore, additional analyses were conducted to determine if NAC TFs also have the potential to bind to the transmembrane domain or if the NAC TFs with a basic pI remain within the cytosol.

**Figure 1.**
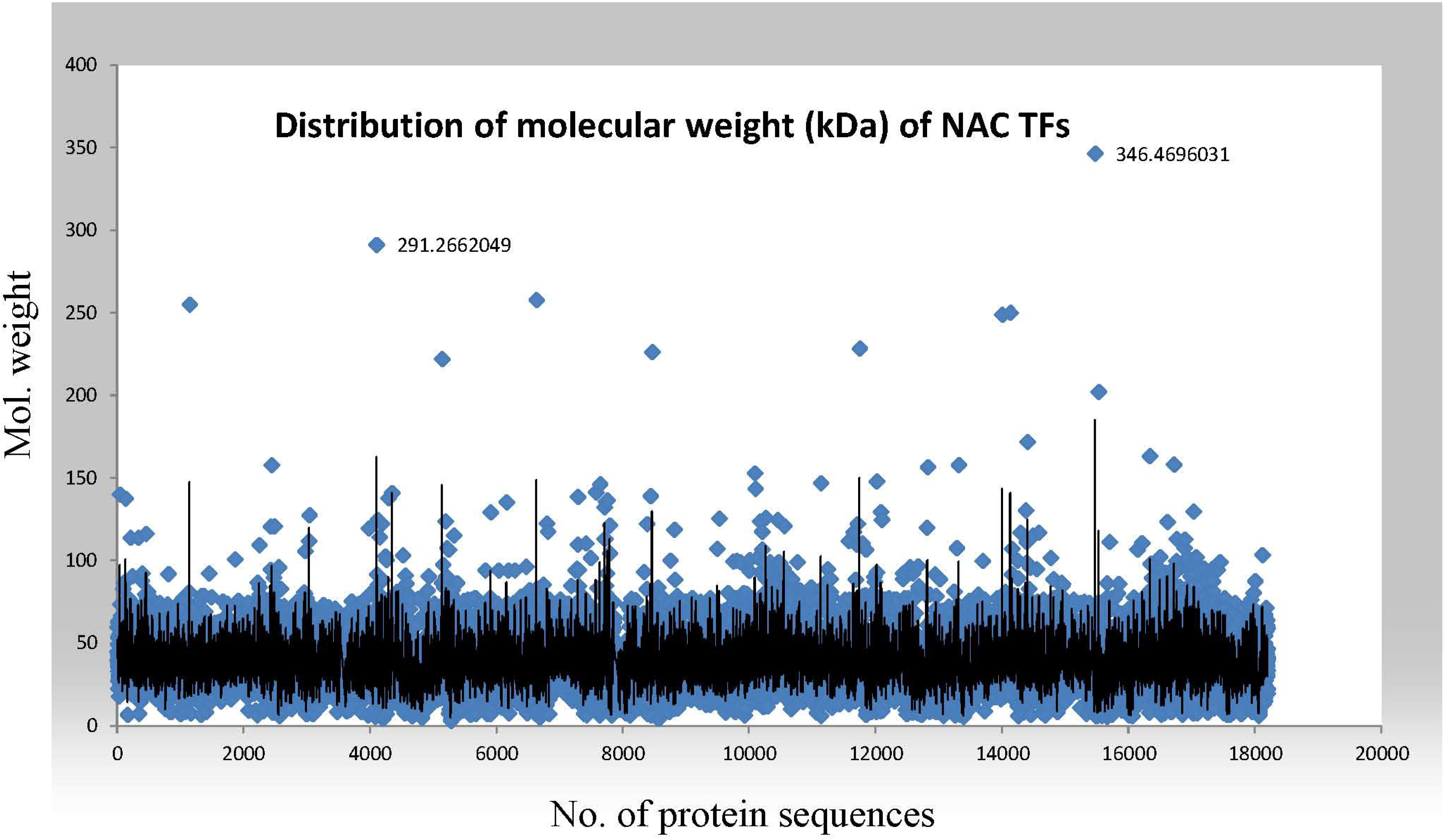
The distribution of the molecular weight of NAC TFs. The molecular weight of NAC TFs ranged from 2.94 kDa (*Fragaria* x *ananassa*, FANhyb_icon00034378_a.1.g00001.1) to 346.46 kDa (*Trifolium pratense*, Tp57577_TGAC_v2_mRNA14116). The average molecular weight of NAC TFs was 38.72 kDa. In total, 17158 NAC TFs were utilized in the analysis of molecular weight. The analysis was conducted using a protein isoelectric point calculator (http://isoelectric.org/).

**Figure 2.**
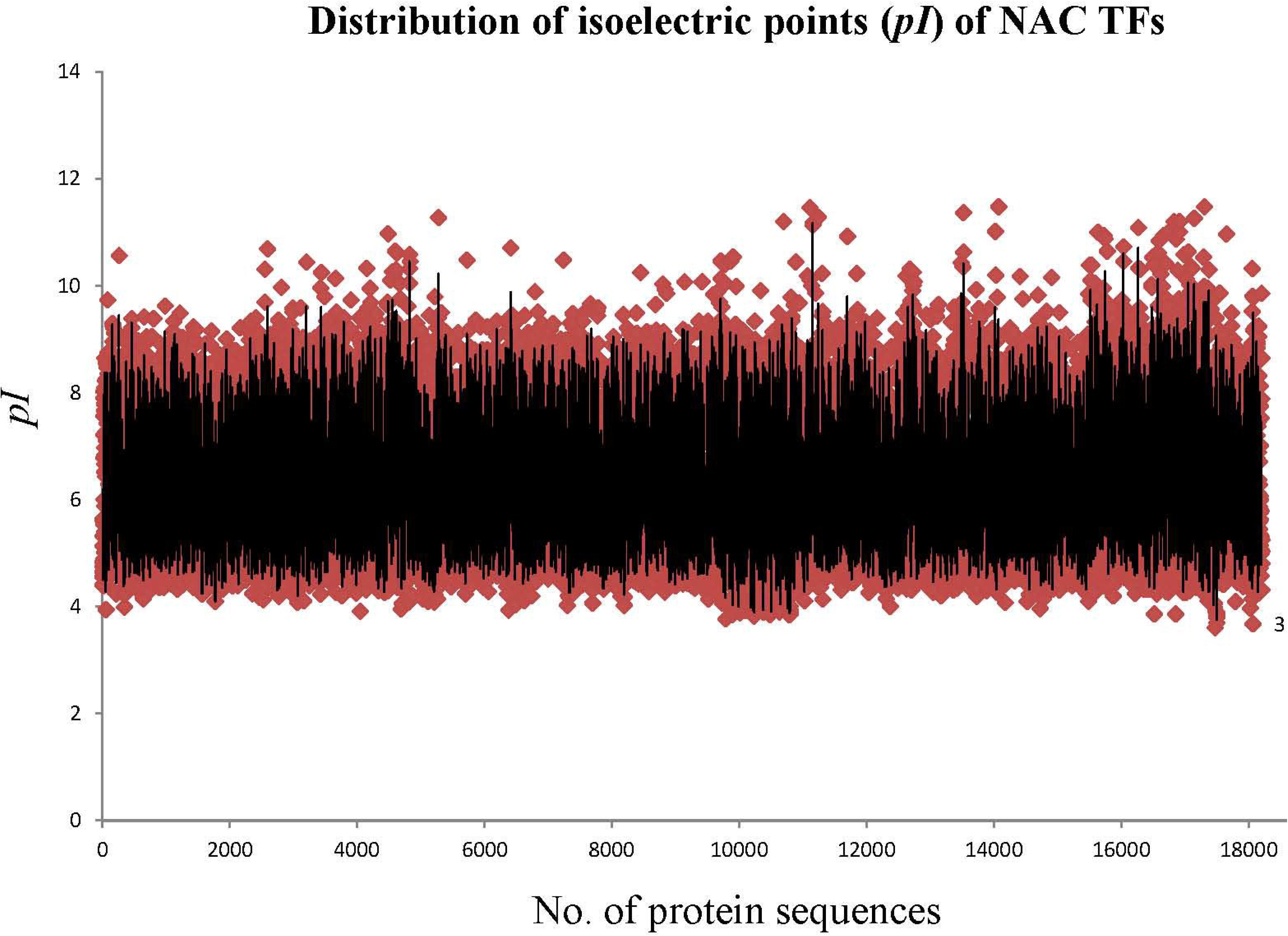
The distribution of the isoelectric point of NAC TFs. The isoelectric point of NAC TFs ranged from pI 3.78 (OB07G17140.1, *Oryza brachyantha*) to pI 11.47 (Sevir.3G242500, *Setaria viridis*). The average isoelectric point of NAC TFs was 6.38. A total of 17158 NAC TFs were utilized in the analysis of the pI of NAC TFs. The analysis of pI was conducted using a protein isoelectric point calculator (http://isoelectric.org/).

### NAC TFs are membrane bound

Transcription factors regulate diverse cellular events at transcriptional, translational, and posttranslational levels. They are also involved in nuclear transport and posttranslational modifications. In several cases, TFs are synthesized but remain inactive in the cytoplasm and are only induced into activity through non-covalent interactions ^21,22^. TFs are able to remain inactive through their physical association with intracellular membranes and are released by proteolytic cleavage. NAC TFs are a family of proteins whose numbers are in the hundreds in the majority of plant species. The fact that NAC TFs are such a large protein family, it is not surprising that NAC TFs have evolved diverse functional roles. Therefore, it is plausible that NAC TFs may be associated with sub-cellular organelle other than the nucleus to fulfil their diverse functional roles. It is essential, however, to confirm if NAC TFs contain signalling sequences for transmembrane localization. Therefore, we analysed the NAC gene sequences to determine if the signalling sequences present in NAC TFs possess a transmembrane domain.

Results indicated that at least 2190 (8.57%) NAC TFs possess a transmembrane domain. Transmembrane domains were found at both the N- and C-terminal ends of NAC proteins. In the majority of the cases, however, the transmembrane domain was located towards the C-terminal end. Seo et al., (2008) indicated the presence of a transmembrane domain in TFs and suggested that transmembrane domain functions through two proteolytic mechanisms, commonly known as regulated ubiquitin/proteasome-dependent (RUP) and regulated intramenbrane proteolysis (RIP) ^23,24^. The bZIP plant TF is present as an integral membrane protein associated with stress response in the endoplasmic reticulum (ER) ^25–28^. Studies suggest that the majority of membrane bound TFs are associated with the ER and a membrane bound TF was also found to be involved in cell division ^29,30^. At least 10% of the TFs in *Arabidopsis thaliana* have been reported to be transmembrane bound ^30^. The collective evidence clearly indicates that membrane-mediated transcriptional regulation is a common stress response and that NAC TFs play a vital role in stress resistance in the ER. Therefore, these membrane-bound NAC TFs can be of great importance for the manipulation of stress resistance using biotechnology.

### NAC TF contain monopartite, bipartite, non-canonical, and nuclear export signal sequences

The import of NAC TFs into the nucleus is mediated by nuclear membrane-bound importins and exportins that form a ternary complex consisting of importin α, importin β1, and a cargo molecule. Importin α serve as an adaptor molecule of importin β1 and recognises the nuclear localization signal (NLS) of the cargo protein needing to be imported. Importin β1 and β2, however, also recognize the NLS directly and bind to the cargo protein. Although the NLS of TFs have been widely studied in the animal kingdom, their study in plants has been more restricted. Therefore, the NLS of NAC TFs was examined in the current study. Results indicate that NAC TFs contain diverse NLS. The NLS were found in the N- and C-terminal regions of NAC TF proteins. Some NAC TFs were found to contain only one NLS whereas other contain multiple NLS. At least 3579 of the total NAC TFs analysed were found to contain either one or multiple NLS. More specifically, 2604 NAC TFs were found to possess only one NLS at the N-terminal end of the NAC protein, whereas 975 were found to possess two NLS, 254 possess three NLS, and 48 were possess four NLS. The NLS were located towards the N-terminal end in the majority of NAC proteins.

NLS motifs are rich in positively charged amino acids and bind to importin α to be imported into the nucleus. The NLS motifs are classified as monopartite or bipartite. A monopartite NLS contains a single cluster of positively charged amino acids and are grouped into two subclasses, class-I and class-II. Class-I possesses four consecutives positively charged amino acids and class-II contains three positively charged amino acids, represented by K(K/R)-x-K/R; where x represents any amino acid that is present after two basic amino acids. Bipartite NLS motifs contain two clusters of positively charged amino acids separated by a 10-12 amino acid linker sequence. Bipartite NLS motifs are characterised by the consensus sequence K-R-P-A-A-T-K-K-A-G-Q-A-K-K-K-K. In addition to monopartite and bipartite NLS motifs, importin α also recognises non-canonical NLS motifs. Non-canonical NLS motifs are longer and considerably variable relative to monopartite and bipartite NLS motifs and are classified as class-III and class-IV NLS. Non-canonical NLS motifs are usually present in the C-terminal end and bind with importin β2. Class-III and class-IV NLS motifs contain K-R-x(W/F/Y)-x_2_-A-F and (P/R)-x_2_-K-R-(K/R) consensus sequences, respectively. We identified at least 1702 unique NLS consensus sequences in the N-terminal region of NAC TFs. The monopartite class I NLS motifs were found to contain more than four consecutive basic amino acids with the number of their consecutive basic amino acids ranging from four to fourteen (K-K-K-K-K-K-K-K-K-K-K-K-K-K-K). The bipartite NLS motifs contain two clusters of consecutive basic amino acids separated by up to twenty-four linker amino acids (K-K-K-x_3_-R- x_2_-R- x_4_-K- x_3_-K- x_3_-K-x-K- x_2_-R-K-K).

The non-canonical NLS motifs contain at least six centrally-located, positively charged amino acids (K-x-R-R-R-P-R-R-x_2_-R-K) flanked by positively charged amino acids on both sides. Our analysis of the N-terminal NLS of NAC TFs, however, did not identify any NAC TFs containing this consensus sequence. Instead, several new variants of this consensus sequence were identified with multiple clusters of positively charged amino acids. These NLS were designated as multipartite NLS motifs. A few examples of the multipartite NLS include, <u>K-K-K-K</u>-x_7_-<u>K-K-K-</u> <u>K</u>-x_7_-<u>K-K-K-K</u>, <u>K-K-K-K</u>-x-K-x_5_-K-x-K-K-x_7_-<u>K-K-K-K</u>-x_2_-<u>K-K-K</u>, <u>K-K-K</u>-x_2_-K-K-x-K-x_5_-K-x_4_-<u>K-K-K-R</u>-x-<u>K-R-K</u>-x-K-x_4_-<u>K-K-K-R-K-K</u>, <u>K-K-R-</u>x-R-K-x_2_-K-x-K-x_2_-<u>K-K-K</u>-x-RK-x_2_-<u>K-R-R</u>-x_2_-<u>K-K-K</u>-x-R, <u>K-K-R</u>-x-R-K-x_2_-K-x-K- x_2_-<u>K-K-R</u>-x-R-K-x_2_-K-x-K-x_2_-<u>K-K-R</u>-x-R-K-x_2_-K-x_2_-K-x-K-x-R, K-x_2_-<u>K-K-K</u>-x_3_-<u>K-K-K-K-K</u>-x-K-x_8_-K-x_9_-K-x_2_-<u>K-K-R</u>-x_2_-<u>K-K-K-K</u>-x-K, K-x_2_-<u>K-K-K</u>-x_3_-<u>K-x-K-K-K</u>-x-<u>K-K-K</u>-x_2_-<u>K-K-K</u>-x-K, <u>R-K-R</u>-x-R-x-<u>R-K-K</u>-x_2_-K-x-<u>K-K-K-R</u>-x_2_-K-x_2_-KK-x_2_-<u>R-R-K</u>-x_2_-K, and <u>R-K-R</u>-x-R-x-R-x_2_-K-x-<u>K-K-K-R</u>-x_2_-K-x4-K-R-x_2_-<u>R-R-K</u>-x-K-x_2_-R. Much of the diversity of NLS motifs is associated with the sequence of the variable linker amino acids. In our analysis, we removed the linker amino acid sequences, represented as x, to obtain a more concise picture of NLS diversity. Removing the linker amino acids present in monopartite, bipartite, and multipartite NLS motifs resulted in the identification of 97 different NLS consensus sequences in the N-terminal region of NAC TFs. The unique NLS signal sequences were R-K-R-R-K, K-K-K, K-R-K, K-K-R, K-R-R, R-R-R, R-K-K, R-K-R, K-K-K-K, R-K-R-K, R-R-K, R-R-R-R, K-K-R-K, K-K-R-K-R, K-R-K-R, R-K-R-R-R, R-K-R-R, K-K-K-K-K, R-R-K-R, K-R-K-R-R-K, R-R-K-K, R-R-R-K, K-R-R-K, K-K-R-R, R-K-R-K-R, R-R-R-R-R, K-R-K-K, K-R-R-R, K-R-K-R-R-R, K-K-K-R, R-K-K-K, K-R-K-K-K, K-R-K-R-K, R-K-K-R, K-R-R-R-R, R-R-K-R-R-K, K-K-K-K-R, K-K-K-R-K, K-K-R-K-K, K-K-K-K-K-K, R-K-K-R-K, R-K-R-K-K, R-K-K-K-K, R-K-R-K-R-K, R-R-K-K-K, R-R-K-R-R-R, R-R-R-R-K, K-K-K-K-K-K-K, K-K-K-K-K-K-K-K, K-K-K-K-R-R, K-K-R-R-R, K-K-R-R-R, R-R-K-R-K-R, R-R-R-K-K-K, R-R-R-R-R-R, R-R-R-R-R-R-K, K-K-K-K-K-R, K-K-K-K-K-R-K, K-K-K-R-K-K, K-K-K-R-R-R-R-R, K-R-K-K-K-K, K-R-K-R-R, K-R-R-K-R, R-K-K-K-K-R, R-K-R-K-K-K, R-K-R-R-K-R-K, R-R-K-K-K-K, R-R-R-K-K, R-R-R-K-R, R-R-R-R-K-K, R-R-R-R-R-K, K-K-K-K-K-K-K-K, K-K-K-K-K-K-K-K-K-K-K-K-K-K, K-K-K-K-K-K-K-K-K-K-K-K-K-K-K, K-K-K-K-K-R-R, K-K-K-K-R-K-R, K-K-K-R-K-R, K-K-K-R-R-R-R, K-K-K-R-R-R-R-R-R, K-K-R-R-R-R-R-R-R-R, K-R-K-K-R, K-R-K-R-K-R-K-K, K-R-R-K-K, R-K-K-K-K-K-R, R-K-K-R-K-K-R, R-K-K-R-K-R, R-K-R-K-R-R, R-K-R-K-R-R-K, R-K-R-R-K, R-K-R-R-K-R, R-R-K-K-R, R-R-K-K-R-K, R-R-K-R-K, R-R-R-K-R-R, R-R-R-R-K-K-K, R-R-R-R-K-R, R-R-R-R-R-R-R, and R-R-R-R-R-R-R-R. The R-K-R-R-K consensus sequence was found to be present 347 times, K-K-K 297 times, K-R-K 185 times, K-K-R 165 times, K-R-R 153 times, R-R-R 96 times, R-K-K 95 times, R-K-R 83 times, K-K-K-K 75 times, R-R-K 74 times, R-R-R-R 58 times, K-K-R-K 49 times, K-K-R-K-R 49 times, and K-R-K-R 40 times. At least 27 NLS amino acid consensus sequences were only found once among the 160 studied species.

The C-terminal end of NAC TF proteins also contain monopartite, bipartite, and multipartite NLS motifs. The multipartite NLS motifs found in the C-terminal end of NAC proteins were <u>R-K-R</u>-x-R-x-<u>R-K-K</u>-x_4_-K-x-<u>K-K-K</u>-R-x_3_-K-x_3_-K-K-x_3_-<u>R-R-K</u>-x_2_-K, <u>R-R-R</u>-x_4_-K-K-x_6_-R-x_2_-R-x_2_-R-R-x_4_-<u>R-R-R</u>-x_6_-R-x_2_-R-R-x_9_-<u>R-R-R-R-R-R-R</u>-x_2_-R-R, <u>K-K-K</u>-x_4_-K-K-x-K-x_5_-K-x_4_-K-<u>K-K-R</u>-x-<u>K-R-K</u>-x-K-x_4_-<u>K-K-K-R-K-K</u>, <u>K-K-R</u>-x_4_-K-x_2_-K-x-K-x_2_-<u>K-K-R</u>-x-R-K-x_4_-K-x_2_-K-x-<u>K-K-R</u>-x-R-K-x_4_-K-x_2_-K-x-K-x-R, <u>K-K-R</u>-x-R-K-x_2_-K-x-K-x_2_-<u>K-K-K</u>-x-R-K-x_2_-<u>K-R-R</u>-x_2_-<u>K-K-K</u>-x-R, <u>K-K-R</u>-x-R-K-x_2_-K-x-K-x_2_-<u>K-K-R</u>-x-R-K-x_2_-K-x-K-x_2_-<u>K-K-R</u>, <u>R-K-R</u>-x-R-x_3_-<u>K-K-R-R</u>-x_2_-K-x_9_-K-x_4_-R-x-K-x_2_-R-x-R-R-x_5_-<u>K-K-R</u>, <u>R-K-R</u>-x-R-x-R-x_5_-K-x-<u>K-K-K-R</u>-x_3_-K-x_4_-K-R-x_2_-<u>R-R-K</u>, R-R-x-<u>R-R-R</u>-x-R-R-x_8_-R-x_6_-R-R-x_5_-<u>R-R-R</u>-x-R-x_5_-R-x_8_-<u>R-R-R-R</u>, R-R-x-R-R-x-R-x-<u>R-R-R</u>-x_9_-R-x_2_-<u>R-R-K-R-K</u>-x-R-x_4_-<u>R-R-R-R-R-R</u>-x_4_-R-K, R-x-<u>R-R-R-R</u>-x_6_-R-x_11_-R-x_8_-R-R-x_3_-<u>R-R-R</u>-x_2_-R-R-x-R-x-R-x_6_-<u>R-R-R-R-R</u>-x_4_-R-R-x_2_-R, R-x-R-R-x_3_-<u>K-R-R-R</u>-x_2_-R-x-R-R-x-R-x-R-x_7_-R-x_3_-<u>R-R-R</u>-x_7_-R-x_2_-<u>R-R-R-R</u>, R-x-R-x<u>-R-R-R</u>-x_3_-<u>R-R-R</u>-x_3_-R-x-R-x_2_-R-x_4_-<u>R-R-R</u>-x_5_-R-K-x-R-x_3_-R-R- x_13_-R-R-x-K-x_5_-R-R-x_6_-<u>K-R-R</u>, and others. Removal of the linker amino acids present in between the consecutive basic amino acids, resulted in the identification of 94 unique consensus sequences. These include K-K-K, K-K-R, R-R-R, K-R-K, K-K-R-K-R, R-K-K, K-K-R-K, R-K-R-K, K-K-K-K, K-R-K-R, K-R-R, R-K-R, R-R-K, R-R-R-R, K-R-K-K, R-K-R-R-K, andothers. The NLS consensus sequence K-K-K was identified 144 times, K-K-R 83 times, R-R-R 65 times, K-R-K 60 times, K-K-R-K-R 58 times, R-K-K 47 times, K-K-R-K 45 times, R-K-R-K 40 times, K-K-K-K 39 times, K-R-K-R 37 times, K-R-R 36 times, R-K-R 35 times, R-R-K 31 times, R-R-R-R 24 times, K-R-K-K 17 times, and R-K-R-R-K 17 times. A comparison of the 97 NLS consensus sequence present in N-terminal region with the 94 NLS sequences present in the C-terminal region indicated that 84 NLS consensus sequences were shared between the N-terminal and C-terminal regions. This indicates that there is a close relationship between the NLS sequences in these two regions. An analysis of the unique NLS consensus sequence in the N-and C-terminal regions indicated that 13 NLS consensus sequences were unique to the N-terminal region, namely R-K-R-R-K, R-R-R-R-K, K-K-K-K-R-R, K-R-K-K-K-K, R-R-K-K-K-K, K-K-K-K-K-R-R, K-K-K-R-K-R, R-K-K-R-K-K-R, R-K-R-K-R-R, R-K-R-R-K-R, R-R-K-K-R-K, R-R-R-K-R-R, and R-R-R-R-K-R. Similarly, nine NLS consensus sequences were unique to the C-terminal region, namely K-K-R-R-K, K-K-K-K-R-K, R-R-K-K-K-R-R-R-R-R-R-R, K-K-R-K-R-K, K-R-R-R-K, R-K-K-R-K-K, R-K-K-R-R, R-R-K-R-R-R-K, and R-R-R-R. Up to six classes of NLS have been reported to be associated with importin α subunit ^31^. To the best of our knowledge, this is the first report describing such a high level of diversity and dynamism in the NLS consensus sequences of NAC TFs and plant transcription factors in general. This is also the first report of the presence of unique NLSs in the N-and C-terminal regions of NAC TFs.

Several nuclear-associated proteins contain NLS, as well as nuclear export signals (NESs). Proteins that perform their function within the nucleus need to be exported out of the nucleus and into the cytoplasm to undergo proteosomal degradation. Therefore, a NES is required in addition to an NLS. A Ran-GTP complex binds directly to an NES and mediates the nuclear export process of cargo molecules ^32^. NES sequences contain a hydrophobic, conserved L-V-F-Y (substitute L-V/I-F-M) motif separated by variable linker amino acids at both ends ^33^. The presence of an L-V-F-Y motif in all NAC proteins, suggests that all NAC proteins have the potential to be exported out of the nucleus. Hao et al. (2010), however, reported that the hydrophobic L-V-F-Y motif functions as a transcriptional repressor of WRKY, Dof, and APETALA TFs. If the L-V-F-Y motif acts as a transcriptional repressor, then the transcriptional activity of these TFs would be lost; resulting in an unstable genome. Therefore, we suggest that the L-V-F-Y motifs do not function as a transcriptional repressor but rather as a NES.

### NAC TFs possess a complex interactome network

The interacting partner of a protein can provide significant information about its potential function and an entire protein-protein interactome network can greatly assist in unravelling the signalling cascade of the proteins. Different cascades are interlinked in signalling systems and form intricate constellations that provide information about cell response and function. Thus, the interactome network of NAC TFs in *A. thaliana* were explored. The presence of a dynamic network was revealed and a diverse set of interacting protein partners of NAC TFs were identified (Figure 3, Table 2). Results indicated that NAC TFs interact with RNS1 (ribonuclease 1), ERD14 (early responsive to dehydration 14), VND1 (vascular related NAC domain 1), VND7, GAI (gibberellins inducible), ZF-HD1 (zinc finger homeodomain 1), TCP8 (<u>T</u>eosinte branched, <u>C</u>ycloidea, and <u>P</u>roliferating cell nuclear antigen factor 8), TCP20, CPL1 (C-terminal domain phosphatise-like1), RHA1A (ring H2 finger H1A), RHA2A, SHR (short root), PHB (phabulosa), PLT2 (plethora 2), MYB59, HB23 (homeobox 23), HB30, NAC1, NAC6, NAC19, NAC32, NAC41, NAC45, NAC50, NAC52, NAC76, NAC83, NAC97, NAC101, NAC105, IAA14 (auxin responsive protein indole3-acetic acid), HAI1 (protein phosphatase), ABI1 (ABA insensitive 1), RVE2 (reveille), PYL4 (PYR-like 4), BRM (brahma), HB52, RCD1 (radical-induced cell death 1), JMJ14 (jumonji 14), TPL (TOPLESS), F2P16 (TOPLESS related), TOPLESS, RING/U-box, ZF-domain (zinc finger domain), SRO1 (similar to RCD 1), CUC2 (cup shaped cotyledon 2), PAS1 (pasticcino 1), TI1 (defensin like 1), TSPO (tryptophan rich sensory protein), TIP2.2 (TCV-interacting protein 2.2), TIP3.1, T21F11.18 (TOPLESS related), LRR (leucine rich repeat), RPA2 (replicon protein A2), and VR-NAC (Table 2). The interaction of NAC TF proteins with the diverse number of listed proteins has been experimentally validated in *A. thaliana.* The interactome network includes stress responsive proteins, other transcription factors, hormonal signalling proteins, protein phosphatases, and defense related proteins. In addition to experimentally-validated interacting proteins, bioinformatic mining indicated that NAC TFs also interact with several other proteins (Table 2). Some of the identified interacting proteins were MYB, NAC, NTL (NTL2-like), UBC30 (ubiquitin conjugating enzyme 30), ATM (Ataxia-Telangiectasia mutated), ATR (serine/threonine kinase ATR), KNAT (knox tail), AOX1A (alternative oxidase 1A), ASG2 (altered seed germination 2), NYE (nonyellowing), CPL, TMO6 (target of monopteros 6), RHA2A (ring H2-finger A2A), XCP (xylem cysteine peptidase), DBP (downstream auxin binding), WAK5 (wall associated kinase 5), RCD, CYP71A25 (cytochrome P71A25), Stay green, ERF (ethylene responsive transcription factor), PPR (pentatricopeptide), WOX (Wuschel related homeobox), PPD6 (psbP-domain protein 6), MFDX (mitochondrial ferredoxin), LEA (late embryogenesis abundant), IRX (irregular xylem), CESA4 (cellulose synthase A4), SCRL20 (SCR-like 20), PIP1-5 (aquaporin), PUP4 (purine permease 4), XERO1 (dehydrin xero 1), SWAP (suppression of white apricot), TIR-NBS, NBS-LRR, PAS, CHI (chitinase), MC5 (metacaspase 5), XCP (xylem cysteine peptidase), RNS, LAC (laccase), TIR-NBS (toll/interleukin receptor-nucleotide binding site), NBS-LRR, DTA4 (downstream target 4), BAG6 (BCL-2 associated anthogene 6), and others as well (Table 2). Some NAC TFs are co-expressed with other proteins. These include DREB2A, XCP1, XCP2, ATM, ATR, MYB63, MYB69, MYB83, IRX1, AOX1A, RCD1, ASG2, ERD1, TMO6, DOF6, SHR, PLT2GRP20, CYP86C4, VND7, NAC6, NAC32, NAC97, NAC19, NAC102, HAI1, WRKY33, WRKY46, WRKY53 and others (Table 2). Some of the NAC TFs directly interact with the interacting partner while others form complexes and appear to play an indirect role. NAC TFs act as a negative regulator of ABA signalling, while they induce JA/ET-associated marker genes ^34^.

**Figure 3.**
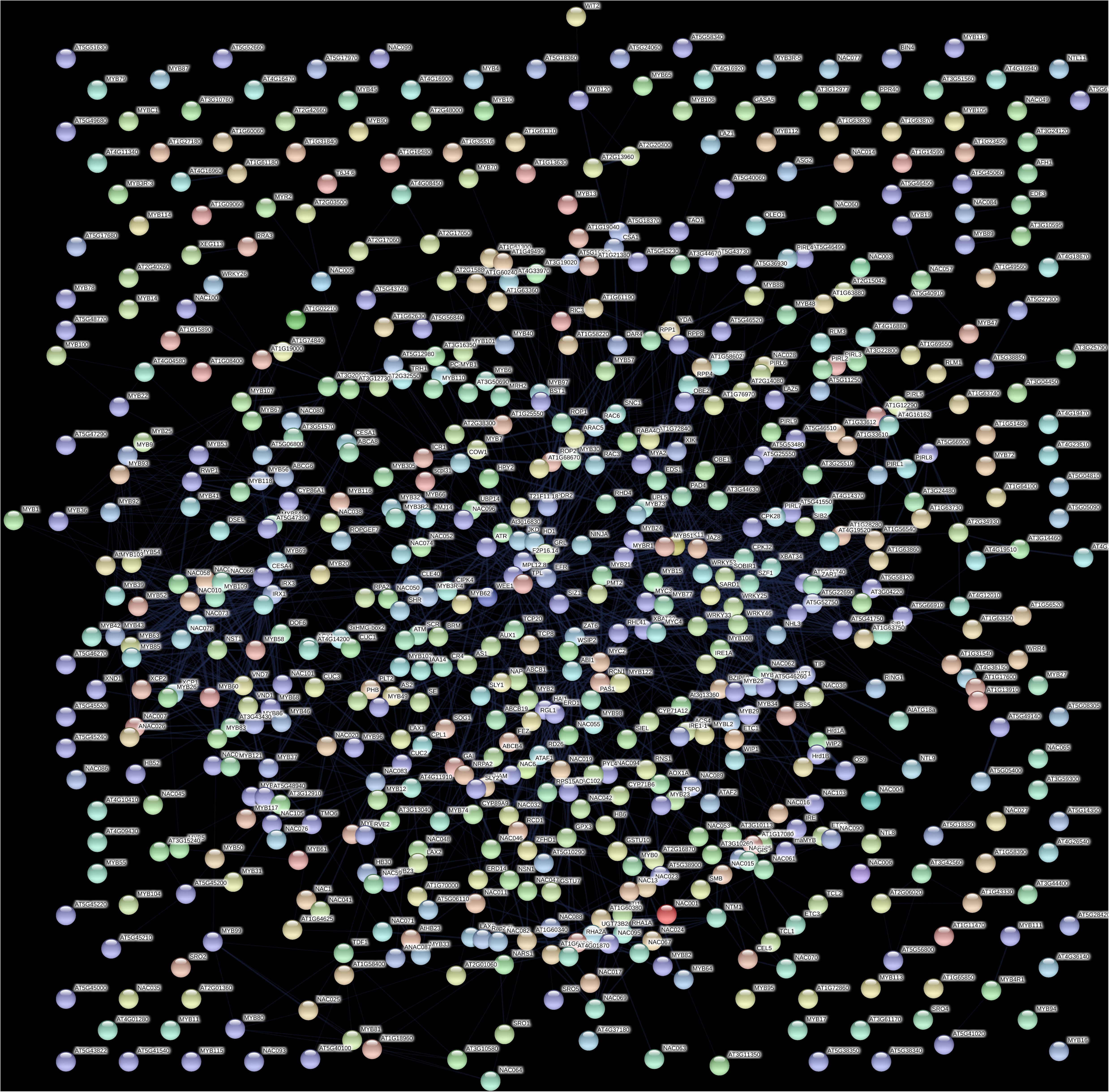
Interactome network of NAC TFs. The interactome network of NAC TF reflects a diverse complex of interacting proteins. The NAC TFs of *A. thaliana* were utilized in the interactome network analysis. The interactome map of *A. thaliana* was determined using the string database (https://string-db.org).

**Table 2.**
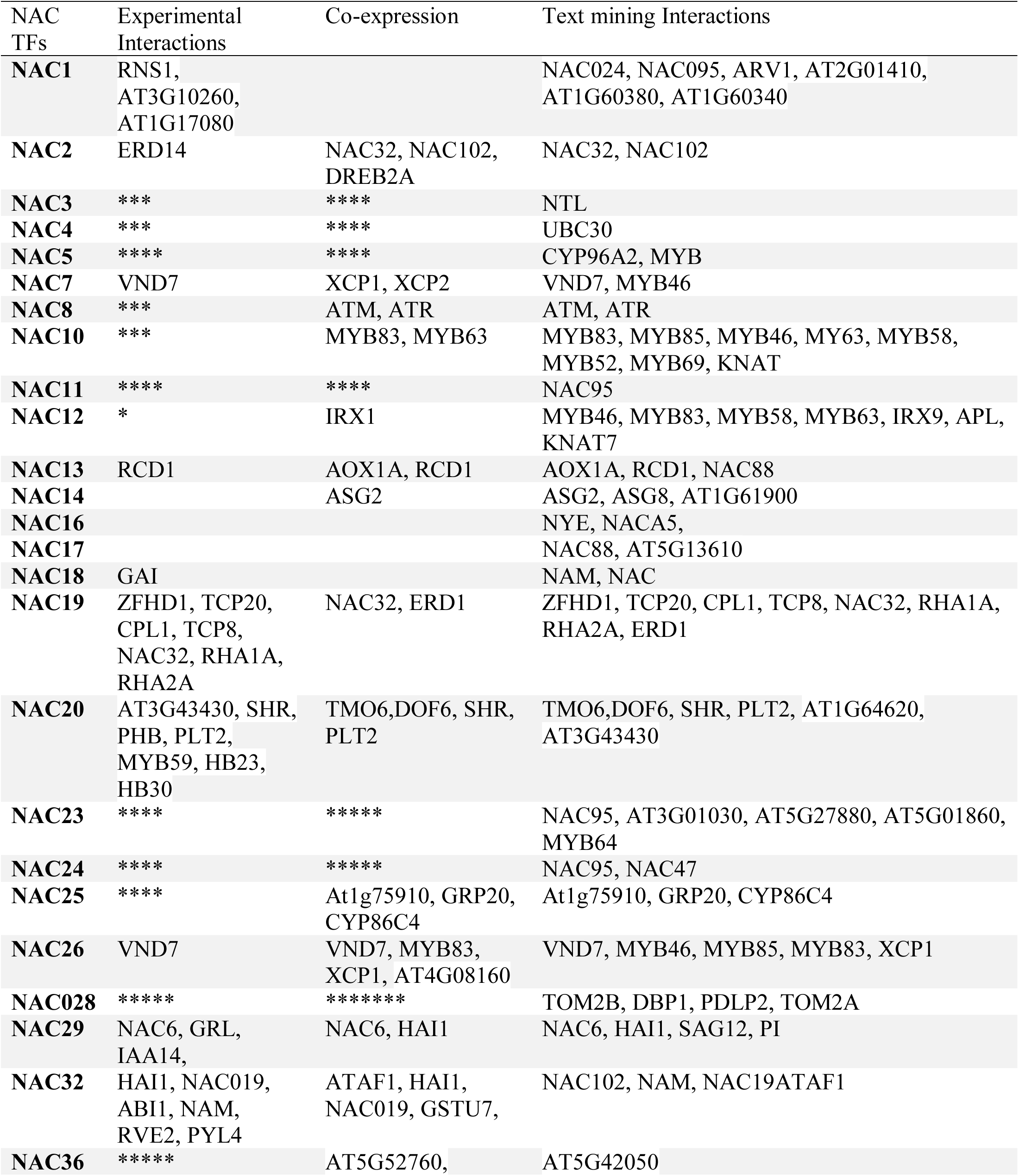
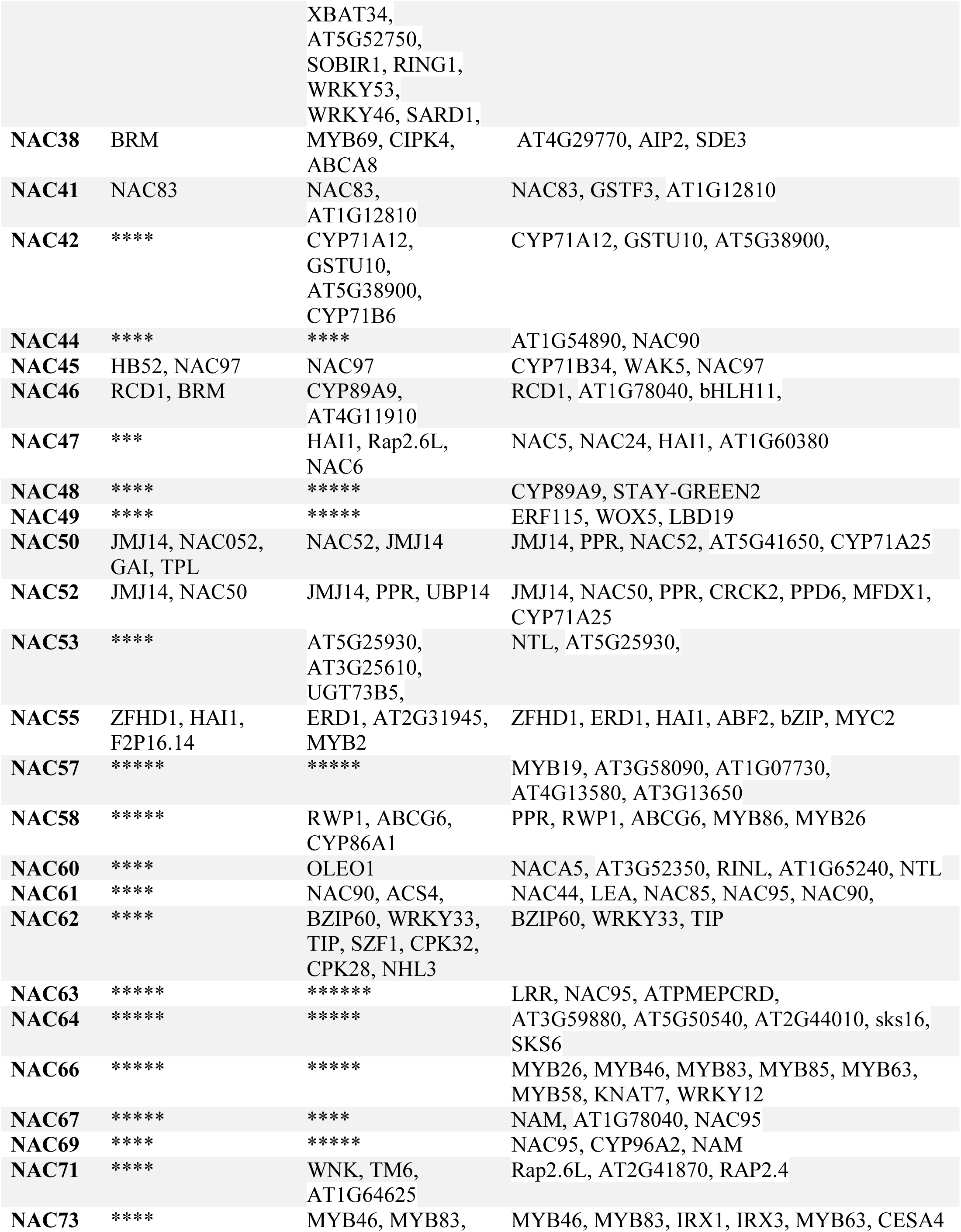
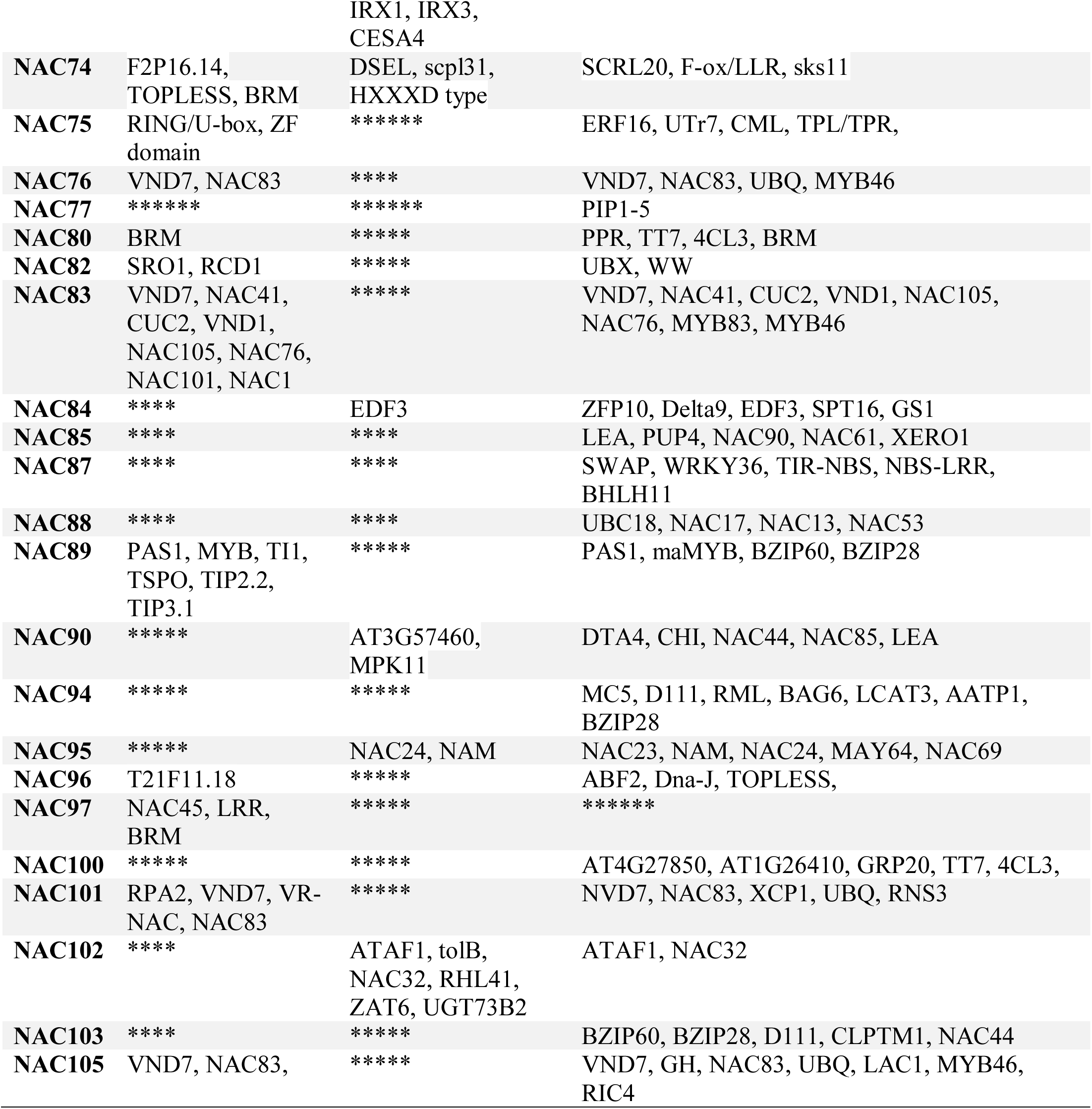
Interactome partners of NAC TFs in plants.

The expression of several of *NAC* genes are either up- or down-regulated by auxin, ethylene, or ABA, suggesting that NAC TFs play a role in plant hormonal signalling ^35–37^. One of the most challenging aspects of a protein-protein interactome network is that the interaction can vary depending upon the cell and its environment ^38^. Therefore, it is necessary to investigate the dynamic interactions of proteins in different cells and environmental conditions to completely understand their interacting partner and the cellular function of the TF. NAC TFs regulate *ERD* and *NCED* (ABA biosynthesis) genes through a direct interaction with their promoters ^39,40^. NAC TFs (ANAC019, ANAC055, and ANAC072) interact with ERD1 which encodes a Clp protease regulatory subunit ^41^. The over expression of one of these three NAC TFs, however, did not induce the up-regulation of ERD1 because the induction of ERD1 depends on the co-expression of a zinc finger homeodomain TF, ZFHD1 ^41^. ANAC019 and ANAC055 interact with ABI (abscisic acid insensitive), and at least five MYB TFs can bind to the NAC TF promoter region ^42,43^. In this case, the NAC DNA binding domain mediates the interaction with RHA2A and ZFHD1 ^43^.

### NAC TFs encodes chimeric proteins and contain multiple binding sites

NAC TFs are characterised by the presence of a DNA binding domain. Several NAC TFs, however, contain more than one NAC domain. Chimeric NAC TFs have also been identified. At least 45 variants of chimeric NAC TFs were identified in our analysis (Figure 4). Several of the NAC TFs were also found to possess as many as three or four NAC DNA binding domains. Furthermore, the NAC domains were found to be associated with PPR (pentatricopeptide), protein kinase, PI3_4_kinase_3, EF-hands (elongation factor), CRM, peptidase A1, WRKY, cytochrome B561, OFOF, FFO, Dna_J2, ZF_B, TIR, LRR, CS, F-box, IQ, PPC, ENT, ABC_TM1F, RWP_RK, PB1, PABC, ACT, INTEGRA, RESPO, JMJC, SAM, BRX, G_TR_2, RORP, CHCH, TPR, YJEF_N, HTH, HOMEO, GH16, ANK_REP_REGION, Peroxidase, LONGIN, V_SNA, RECA_2, KH_TY, APAG, RRM, carrier, and a DCO domain. At least four NAC TFs from *A. thaliana*, ten from *B. napus*, four from *B. rapa*, two from *M. domestica*, four from *P. virgatum*, 17 from *C. sativa*, eight from *D. oligosanthes*, eight from *E. tef*, and five from *L. perrieri* were found to possess 2 NAC domains (Supplementary Table 1). NAC TFs in several other species were also found to contain two NAC domains (Supplementary Table1). When two NAC domains were present, both domains were located towards the N-terminal end. NAC TFs of at least three species, *O. rufipogon*, *B. stacei*, and *Camelina sativa* were found to possess three NAC domains whereas the NAC TFs in *A. lyrata* (gene id: 338342), *C. sativa* (Csa16g052260.1), and *E. tef* (462951506) were found to possess four NAC domains (Figure 4).

**Figure 4.**
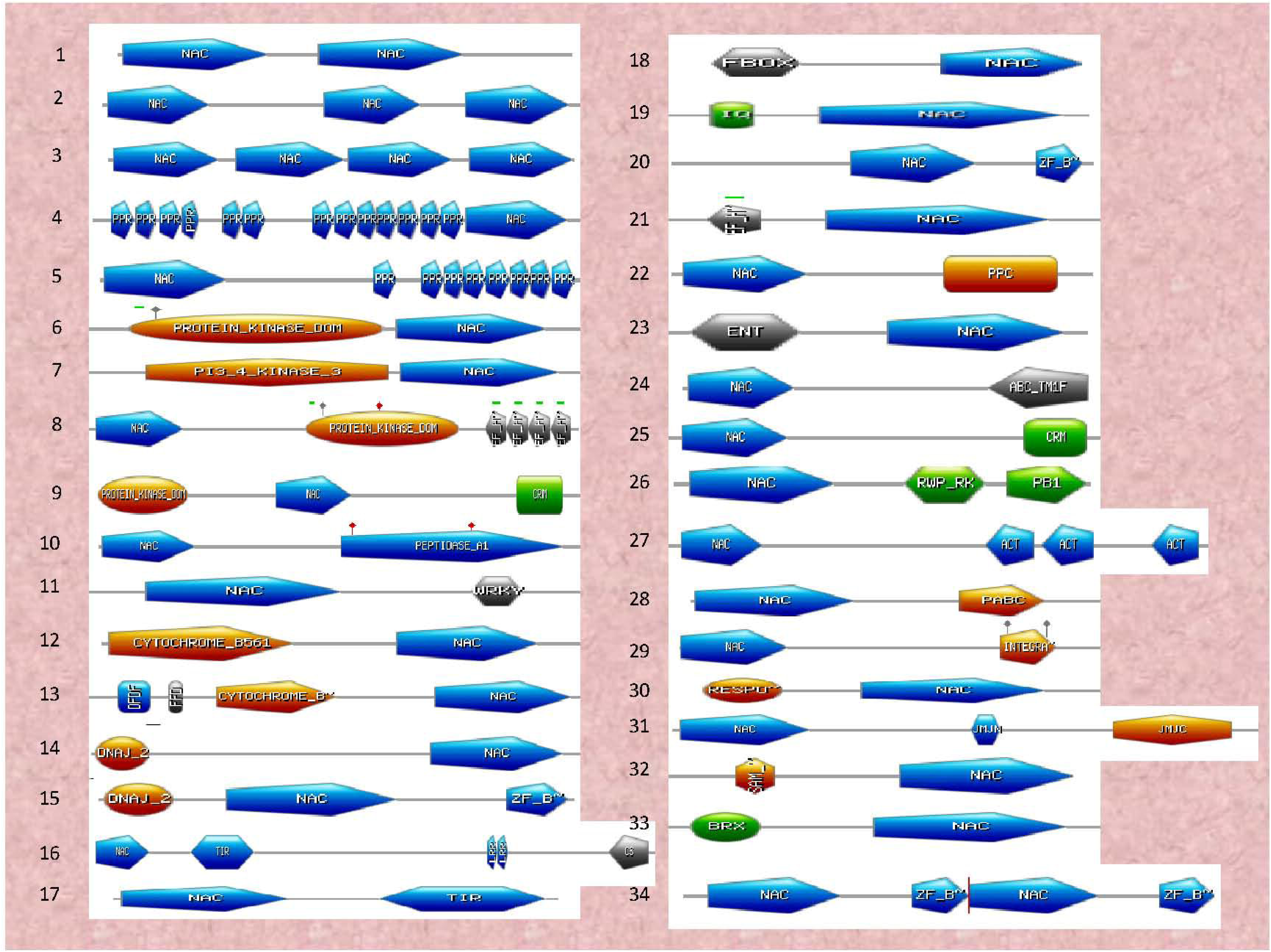
Chimeric NAC domains. NAC TFs possess chimeric NAC domains with at least 34 diverse chimeric NAC domains identified in the studied species. The identification of chimeric NAC domain sequences was determined using the ScanProsite and InterProScan server.

Other chimeric domains were also identified in different regions of the NAC protein. PPR domains were found in both the N-terminal or C-terminal region, a protein kinase domain was found upstream to the NAC domain, and a NAC domain was found to be adjacent to a protein kinase and EF-hand domain. Additionally, a protein kinase domain was found to be followed by either a NAC and a CRM domain, a NAC domain was followed by a peptidase_A1 domain, a NAC domain was followed by the presence of a WRKY domain, a cytochrome_B561 domain was followed by either a NAC domain and a CRM domain, a DFDF and a FFD domain were followed by a cytochrome B and NAC domain, a DNAJ_2 domain was followed by a NAC domain, a DNAJ_2 domain was followed by a NAC and ZF_B domain, a NAC domain was followed by TIR, LRR and CS domains, a NAC domain was followed by a TIR domain, an F-box domain was followed by a NAC domain, an IQ domain was followed by a NAC domain, a NAC domain was followed by a ZF_B domain, an EF-hand domain was followed by a NAC domain, a NAC domain was followed by a PPC domain, an ENT domain was followed by a NAC domain, a NAC domain was followed by an ABC_TM1F, a NAC domain was followed by CRM domain, a NAC domain was followed by a RWP_RK and a PB1 domain, a NAC domain was followed by three ACT domains, a NAC domain was followed by a PABC domain, a NAC domain was followed by an INTEGRA domain, a RESPO domain was followed by a NAC domain, a NAC domain was followed by a JMJN and a JMJC domain, a SAM domain was followed by a NAC domain, a BRX domain was followed by a NAC domain, a NAC domain was followed by ZF_B, NAC and ZF_B domains, an F-box and protein kinase domain was followed by a NAC domain, a NAC domain was followed by a G_TR_2 domain, an RDRP domain was followed by a NAC domain, a NAC domain was followed by a CHCH domain, a TPR domain followed by a NAC domain, an F-box domain followed by NAC and F-box domains, a NAC domain was followed by a YJEF_N domain, a NAC domain was followed by a HTH domain, a homeobox domain was followed by a NAC domain, a NAC domain was followed by three GH16_2 domains, an ANK repeat region was followed by a NAC domain, a NAC domain was followed by a peroxidase domain, a NAC domain was followed by a LONGIN and V_SNA domain, a NAC domain was followed by RECA_2 and RECA_3 domains, a KH domain was followed by a NAC domain, a NAC domain was followed by a RAB domain, a JMJN domain was followed by a NAC domain, a NAC domain was followed by an APAG domain, an RRM domain was followed by a NAC domain, a carrier domain was followed by a NAC domain, and a NAC domain was followed by DCO domain (Figure 4).

The presence of chimeric domains within NAC TFs is of particular interest, especially for understanding why they are there and how they impact the function of a specific NAC TF. The most common domains, such as PPR, TIR, WRKY, protein kinase, ZF_B, EF-hands, cytochrome B, DNAJ, F-box, peroxidase, and GH16 are involved in diverse cellular processes, including transcriptional regulation of plant development and stress response ^44–52^. The association of a TIR domain with an NBS-LRR domain is an example of the association of TF domains with other domains to form chimeric proteins ^53^. The presence of different domains with the NAC domain could potentially enable the NAC domain to assist in the function of the associated domains and vice versa. For example, NAC TFs could have the potential to regulate peroxidase by possessing a peroxidase domain within the NAC TF, instead of regulating it separately with another TF. The presence of multiple domains can enable the co-regulation of diverse functional sites within the NAC TFs. The presence of chimeric TFs has been recently reported in WRKY TFs as well ^54,55^. Therefore, the presence of chimeric domains in NAC TFs can impart a significant dynamic aspect to the ability of NAC TFs to regulate gene expression.

In addition to the presence of multiple chimeric domains, NAC TFs were also found to contain diverse active/binding motifs for several other proteins. It is possible that NAC TFs may play a dual role as a transcription factor and as an enzyme. At least 404 NAC TFs were found to possess other functional motifs comprising 101 unique functional sequences (Supplementary Table 2). In addition to NAC domains, the other function sequences included a Fe-2S ferredoxin-type iron-sulfur binding region signature, 2-oxo acid dehydrogenase acytltransferase component lipoyl binding site, 4Fe-4S ferredoxin-type iron-sulfur binding domain profile, 7,8-dihydro-6- hydroxymethylepterin-pyrophosphokinase signature (30), ABC transporter family signature (2), adenosine and AMP deaminase signature (2), adipokinetic hormone family signature, aldehyde dehydrogenase cysteine active site (4), aldehyde dehydrogenase glutamic acid active site (28), aldo/keto reductase family putative active site signature (4), alkaline phosphatise active site (5), aminoacyl-transfer RNA synthetase class-I signature, aminotransferase class-II pyridoxal-phosphate attachment site (15), antenna complexes beta subunit signature (2), ArgE/dapE/ACY1/CPG2/yscS family signature 1 (2), aspartate and glutamate racemases signature 1 (3), aspartokinase signature (2), ATP binding site and proton acceptor, ATP synthase alpha and beta subunit signature (19), ATP dependent DNA ligase AMP-binding site (2), bacterial regulatory proteins araC family signature, beta-ketoacyl synthases active site (2), C-5 cytosine-specific DNA methylases active site, cadherin domain signature, carbamoyl-phosphate synthase subdomain signature 2, cysteine protease inhibitor signature (19), cytochrome p450 cysteine heme-iron ligand signature (7), endopeptidase Clp serine active site (3), eukaryotic and viral aspartyl proteases active site (10), FGGY family of carbohydrate kinase signature 2, fumarate lyases signature, GHMP kinases putative ATP-binding domain, glucoamylase active site region signature, glyceraldehyde 3-phosphate dehydrogenase active site, glycoprotease family signature, glycohydrolase family 5 signature, glycosyl hydrolase family 9 active site signature 2, heavy metal associated domain, hemopexin domain signature (2), histone H4 signature (4), HMG-I and HMG-Y DNA binding domain (A+T hook) (9), immunoglobulins and major histocompatibility complex protein signature (4), inorganic pyrophosphate signature (13), iron-containing alcohol dehydrogenase signature 1, legume lectins beta-chain signature (15), lipocalin signature (23), mannitol dehydrogenase signature, N-6 adenine-specific DNA methylases signature (7), neutral zinc metallopeptidase, zinc binding region signature, Nt-DnaJ domain signature (2), peroxidase active site signature, pfkB family of carbohydrate kinases signature 1 and 2 (4), phospholipase A2 histidine active site (5), phosphopantetheine attachment site (17), polygalacturonase active site (2), polyprenyl synthases signature, PPM-type phosphatase domain signature, prokaryotic membrane lipoprotein lipid attachment site, putative AMP binding domain signature, regulator of chromosome condensation (RCC1) signature 2 (2), ribosomal protein L24e signature (7), ribosome binding factor A signature, rubredoxin signature (2), serine protease, subtilase family aspartic acid active site (21), serine protease, trypsin family, serine active site, serine/threonine protein kinase active-site signature (3), sigma-54 interaction domain ATP-binding site A signature, signal peptidase I serine active site (3), signal peptidase I signature 3 (4), soybean trypsin inhibitor (Kunitz) protease inhibitor family signature, SRP54-type proteins GTP-binding domain signature, sugar transport proteins signature 2, synaptobrevin signature, syntaxin/epimorphin family signature, TonB-dependent receptor proteins signature 1 (7), translationally controlled tumor protein (TCTP) domain signature 2, Trp-Asp (WD) repeats signature (12), tubulin subunit alpha, beta, and gamma signature (2), tubulin-beta mRNA autoregulation signal (2), zinc carboxypeptidases, zinc binding region 2 signature (11), zinc finger BED-type profile, zinc finger C2H2 type domain signature, and a zinc-containing alcohol dehydrogenase signature (2) (Supplementary Table 2). This is the first study to report the presence of such a diverse number of functional sites and signature motifs in NAC TFs. Although the majority of the functional domains are associated with a specific function in plants, the presence of a histocompatibility complex and a translationally controlled tumor protein (TCTP) sequence are of particular interest. These proteins are specifically found in animal systems and the histocompatibility complex is the major contributing factor regulating the binding of antigens. More specifically, TCTP is a highly conserved protein that is involved in microtubule stabilization, calcium binding, and apoptosis and is associated with the early growth phase of tumors ^56^. The presence of MHC and TCTP in association with NAC domains suggests that this combination may be playing a crucial role in the plant immune system and in uncontrolled cell growth. The presence of diverse functional sites in NAC TFs indicates that NAC TFs are involved in diverse cellular functions and metabolic pathways. This statement is supported by the large number of NAC TFs that are present in plant genomes.

### NAC TFs are involved in diverse cellular processes

NAC TFs are known to possess diverse chimeric domains, as a result, it is more than likely that NAC TFs are also involved in the regulation of diverse cellular pathways and cellular processes. To help substantiate this premise, the interactome associated with NAC TFs in *A. thaliana* was analysed. Results indicated that NAC TFs are potentially involved in a least 289 different cellular processes and pathways (Supplementary Table 3). The majority are related to cell, tissue, and organ (root, stem, meristem) development, as well as signalling processes. Several NAC TFs also appear to be associated with phytohormone signalling, including auxin, gibberellin, jasmonic acid, and salicylic acid signalling pathways. NAC TFs were also found to be associated with pathways involved in the response to bacterial, fungal, UV, heat and other biotic and abiotic stresses (Supplementary Table 3). At least 202 genes in the NAC TF interactome network were found to be associated with pathways related to the nucleus, 239 were associated with intracellular membranes, and 241 were associated with intracellular organelles, 20 with the endoplasmic reticulum, and 3 with the nuclear matrix. If the association is designated based on the description of a pathway, 127 genes were found to be associated with transcription factor activity and sequence-specific DNA binding, 143 with DNA binding, 146 with nucleic acid binding, 220 with organic cyclic compound binding, 220 with heterocyclic compound binding, 65 with ATP binding, 49 with macromolecular complex binding, 48 with chromatin binding, 35 with ADP binding, 25 with sequence-specific DNA binding, 18 with transcription regulatory region binding, 8 with structural constituents of the cell wall, 11 with auxin transport activity, 2 with LRR binding, and 2 with bHLH transcription factor binding. These data clearly indicate that NAC TFs are involved in diverse cellular processes. The identification of LRR protein in the pathway description of NAC TFs agrees with the presence of an LRR domain in a chimeric NAC domain of NAC TFs.

### NAC TFs are expressed in a spatiotemporal manner

Patterns of NAC TF gene expression were analysed in leaf and root tissues of *A. thaliana* treated with ammonia, nitrate, or urea (Figure 5 and Figure 6). Among a total of 120 *NAC* TFs, 95, 97, and 98 were differentially expressed in leaf tissue treated with ammonia, nitrate, or urea, respectively. Leaf tissues treated with ammonia, nitrate and urea exhibited 70.14, 117.11, and 58.35 FPKM expression values for *AtNAC1* (AT1G01010.1), *AtNAC4* (AT1G02230.1), and *AtNAC1* (AT1G01010.1), respectively. At least 46 genes in leaves exhibited expression of more than one FPKM in response to ammonia, 54 in response to nitrate, and 44 in response to urea. *AtNAC1* was highly expressed in ammonia and urea treated leaves. At least 24, 26, and 25 *NAC* TFs did not exhibit any expression in leaf tissues treated with ammonia, nitrate, or urea.

**Figure 5.**
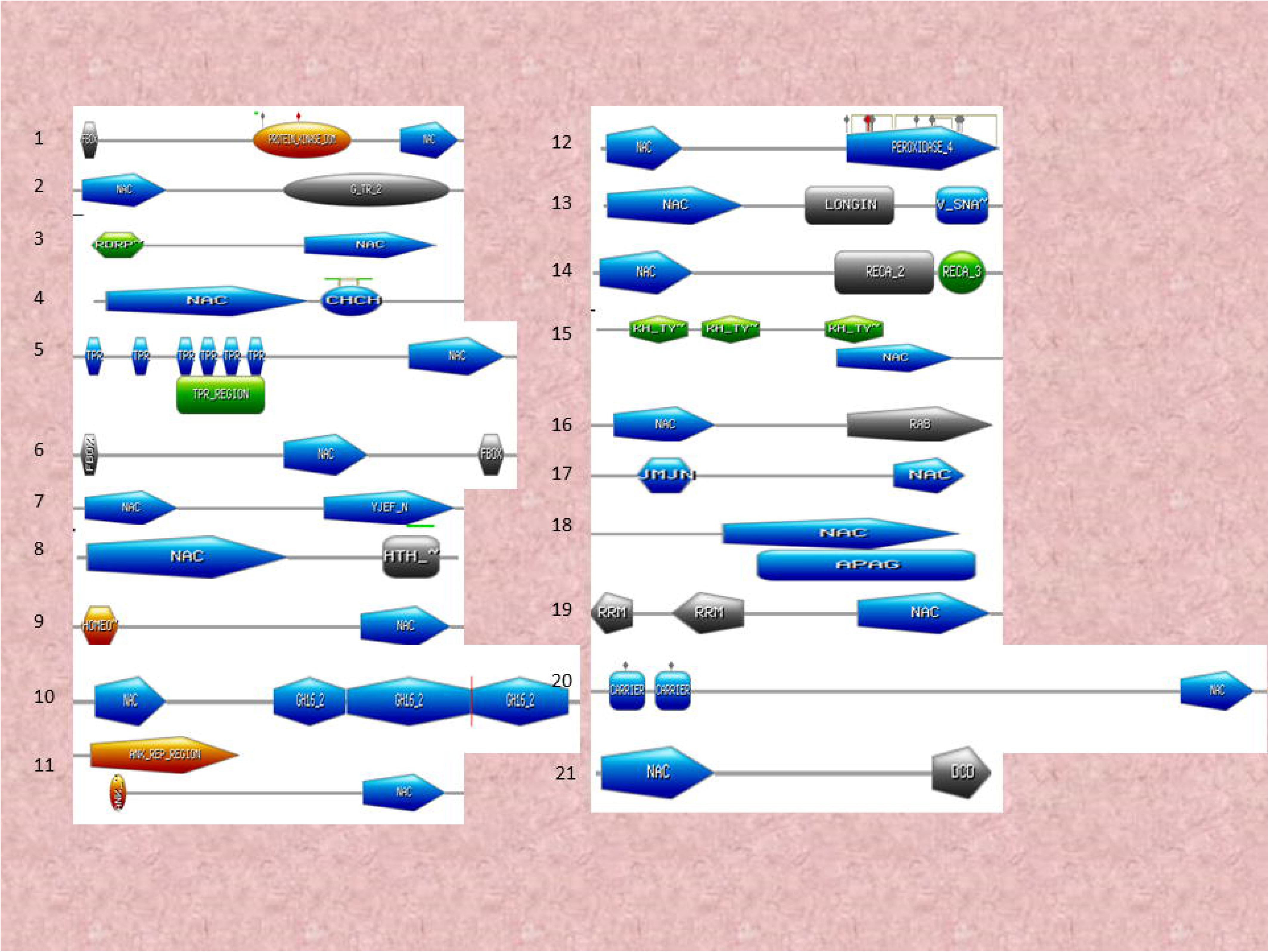
Chimeric NAC domains NAC TFs possess chimeric NAC domains with at least 21 diverse chimeric NAC domains identified in the studied species. The identification of chimeric NAC domain sequences was determined using the ScanProsite and InterProScan server.

**Figure 6.**
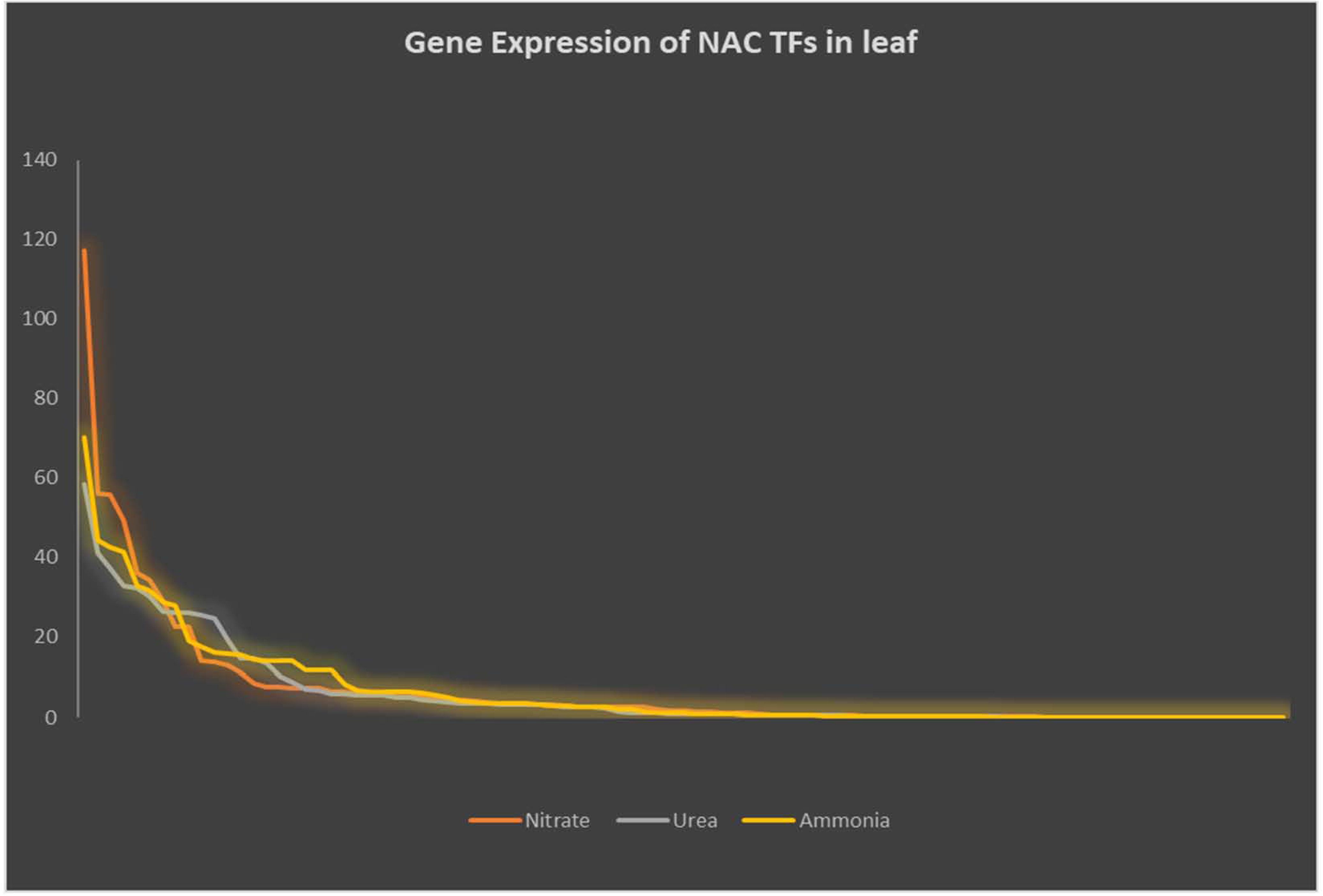
Differential expression of NAC TFs in leaves of *A. thaliana* plants treated with ammonia, nitrate, and urea. The expression of *A. thaliana* NAC TFs was analysed to determine their response to different sources of nitrogen. Expression data were obtained from the PhytoMine database in Phytozome and presented as FPKM (Fragments per Kilobase of transcripts per million mapped reads).

Relative to leaf tissues, the expression of *NAC* TFs in root tissues was more dynamic. Root tissue treated with urea exhibited the highest expression of NAC TFs relative to leaves treated with ammonia or nitrate (Figure 6). The number of *AtNAC* TFs whose expression was one or more FPKM in response to ammonia, nitrate, or urea were 75, 71, and 70, respectively. *AtNAC8* (AT5G08790.1) was highly expressed in ammonia-treated roots, whereas, *AtNAC91* (AT5G24590.2) was highly expressed in nitrate- and urea-treated roots. Urea, ammonia and nitrate (UAN) commonly serve as a source of nitrogen (N) for plants. Analysis of the levels of gene expression indicate that ammonia and nitrate modulate the expression of *NAC* TFs more than urea. A study utilizing *Pinus taeda* revealed that fertilization with ammonium, nitrate, or urea produces different effects on growth and drought tolerance ^57^. Results of the current analysis indicate that *AtNAC8* and *AtNAC91* are the major NAC TFs involved in nitrogen assimilation during plant growth.

### Codon usage in NAC TF is dynamic

Codon usage bias in NAC TFs of the examined species were studied. separately. Among 61 sense codons, only 14 were found in the all species. These included AAG (K), ACU (R), AGA (R), AGG (R), UCU (S), AUC (I), AUG (M), CAA (Q), CCU (P), GAA (E), GCU (A), GGA (G), UGG (0), and UUC (F) (Table 3). The most abundant codon was UCU (S), which was found 30 times in in *Humulus lupulus* NAC TFs (Table 3). The codons CGA (R), CGC (R), CGG (R), CGU (R) were absent in 127 of the 160 examined species. ACG (T), UCG (S), CAG (Q), CAC (H), CCA (P), CCC (P), CCG (P), and GCG (A) were absent in 126 of the examined species. The highest relative synonymous codon usage bias (RSCU) was found to be 1.35, 1.23, 1.29 for the codon AAA (K) in *Ocimum tenufolium, Picea sitchensis*, and *Ipomea trifida*. Synonymous codon-usage was not observed in NAC TFs. Relative codon usage is determined by dividing the ratio of observed frequency of codons by the expected frequency, provided that all of the synonymous codons for the same amino acids are used equally. Relative Synonymous Codon Usage (RSCU), however, is not related to the usage of amino acids. An RSCU > 1 indicates the occurrence of codons more frequently than expected, while an RSCU < 1 indicates that the codon occurs less frequently than expected ^58,59^. Non-synonymous substitution in organisms is subject to natural selection ^60,61^. Genes with lower non-synonymous selection leads to functional diversity of a gene. The presence of a low level of nonsynonymous codon usage in NAC TFs indicates that they are functional and have evolved from paralogous ancestors.

**Table 3.** Codon usage of NAC TFs in plants.

| Codons | Codon present in No. of species | Codon absent in No. of species | Average abundance of codons | Highest no. of codons | Name of the species with highest no. of codons |
| --- | --- | --- | --- | --- | --- |
| AAA (K) | 126 | 20 | 4.77 | 9.9 | *Glycine soja* |
| AAG (K) | 146 | 0 | 10.75 | 24.2 | *Sphagnum fallax* |
| AAC (N) | 144 | 2 | 3.66 | 14.2 | *Beta vulgaris* |
| AAU (N) | 127 | 19 | 9.25 | 20.5 | *Spinacia oleracea* |
| ACA (T) | 139 | 7 | 2.33 | 15.2 | *Citrus sinensis* |
| ACC (T) | 137 | 9 | 2.4 | 17 | *Amborella trichopoda* |
| ACG (T) | 20 | 126 | 5.91 | 13 | *Doroceras hygrometricum* |
| ACU (T) | 146 | 0 | 7.42 | 16.6 | *Sesamum indicum* |
| AGA (R) | 146 | 0 | 10.92 | 24.3 | *Klebsormidium flaccidum* |
| AGG (R) | 146 | 0 | 4.12 | 18.8 | *Amborella trichopoda* |
| CGA (R) | 19 | 127 | 5.22 | 13.9 | *Linum usitatissimum* |
| CGC (R) | 19 | 127 | 2.47 | 6 | *Linum usitatissimum* |
| CGG (R) | 19 | 127 | 3.93 | 8.6 | *Citrullus lanatus* |
| CGU (R) | 19 | 127 | 2.06 | 4.7 | *Linum usitatissimum* |
| AGC (S) | 143 | 3 | 3.54 | 24.2 | *Beta vulgaris* |
| AGU (S) | 144 | 2 | 1.83 | 5.2 | *Doroceras hygrometricum* |
| UCC (S) | 141 | 5 | 4.51 | 12.3 | *Aegilops tauschii* |
| UCG (S) | 20 | 126 | 2.64 | 6.4 | *Doroceras hygrometricum* |
| UCU (S) | 146 | 0 | 4.65 | 30.5 | *Humulus lupulus* |
| UCA (S) | 139 | 7 | 5.09 | 15.1 | *Morus notabilis* |
| AUA (I) | 124 | 22 | 4.80 | 15.3 | *Sphagnum fallax* |
| AUC (I) | 146 | 0 | 5.10 | 16.7 | *Sphagnum fallax* |
| AUU (I) | 126 | 20 | 8.71 | 15.9 | *Spinacia oleracea* |
| AUG (M) | 146 | 0 | 7.81 | 22.8 | *Sphagnum fallax* |
| CAA (Q) | 146 | 0 | 5.31 | 15.4 | *Fragaria vesca* |
| CAG (Q) | 20 | 126 | 13.3 | 22.6 | *Linum usitatissimum* |
| CAC (H) | 20 | 126 | 6.64 | 10.9 | *Beta vulgaris* |
| CAU (H) | 144 | 2 | 4.45 | 9.7 | *Setaria viridis* |
| CCA (P) | 20 | 126 | 11.09 | 16.3 | *Doroceras hygrometricum* |
| CCC (P) | 20 | 126 | 14.18 | 19.2 | *Amborella trichopoda* |
| CCG (P) | 20 | 126 | 5.10 | 11.1 | *Doroceras hygrometricum* |
| CCU (P) | 146 | 0 | 8.00 | 24.7 | *Klebsormidium flaccidum* |
| CUA (L) | 143 | 3 | 5.83 | 28.3 | *Sphagnum fallax* |
| CUC (L) | 123 | 23 | 5.74 | 23.6 | *Sphagnum fallax* |
| CUG (L) | 142 | 4 | 5.87 | 43.9 | *Sphagnum fallax* |
| CUU (L) | 145 | 1 | 5.94 | 32.6 | *Sphagnum fallax* |
| UUG (L) | 125 | 21 | 5.94 | 24.4 | *Sphagnum fallax* |
| UAA (L) | 124 | 22 | 5.37 | 17.2 | *Sphagnum fallax* |
| GAA (E) | 146 | 0 | 4.62 | 27 | *Klebsormidium flaccidum* |
| GAG (E) | 145 | 1 | 5.54 | 18.1 | *Sphagnum fallax* |
| GAC (D) | 145 | 1 | 5.05 | 14.9 | *Beta vulgaris* |
| GAU (D) | 144 | 2 | 5.86 | 21.7 | *Spinacia oleracea* |
| GCA (A) | 135 | 11 | 5.49 | 18.5 | *Citrus sinensis* |
| GCC (A) | 130 | 16 | 5.05 | 15 | *Amborella trichopoda* |
| GCG (A) | 20 | 126 | 4.64 | 11.2 | *Doroceras hygrometricum* |
| GCU (A) | 146 | 0 | 4.65 | 31.1 | *Setaria viridis* |
| GGA (G) | 146 | 0 | 4.63 | 27.5 | *Setaria viridis* |
| GGC (G) | 141 | 5 | 5.41 | 17.1 | *Amborella trichopoda* |
| GGG (G) | 145 | 1 | 2.7 | 6.7 | *Elaeis guineensis* |
| GGU (G) | 145 | 1 | 2.40 | 5.9 | *Elaeis guineensis* |
| GUA (V) | 140 | 6 | 1.46 | 3.5 | *Sphagnum fallax* |
| GUC (V) | 123 | 23 | 0.93 | 2 | *Morus notabilis* |
| GUG (V) | 142 | 4 | 4.35 | 11.8 | *Beta vulgaris* |
| GUU (V) | 143 | 3 | 5.38 | 16.6 | *Klebsormidium flaccidum* |
| UAC (Y) | 138 | 8 | 3.84 | 10.1 | *Morus notabilis* |
| UAU (Y) | 126 | 20 | 6.23 | 14.8 | *Solanum melongena* |
| UGG (W) | 147 | 0 | 3.89 | 14.5 | *Vitis vinifera* |
| UGC (C) | 143 | 3 | 5.14 | 15.6 | *Oropetium thomaeum* |
| UGU (C) | 145 | 1 | 3.9 | 9.6 | *Zoysia matrella* |
| UUC (F) | 146 | 0 | 4.60 | 25.4 | *Picea glauca* |
| UUU (F) | 126 | 20 | 10.67 | 19.2 | *Sphagnum fallax* |

### Rate of transition of NAC TFs is higher than the rate of transversion

Nucleotide mutation is an integral part of the evolution of a genome and leads to the acquisition of required traits and the elimination of detrimental traits from the genome. It is a regular process and hundreds of thousands of nucleotides have undergone addition or deletion events in the evolution of a genome. The alteration or conversion of a nucleotide occurs either through a transition or a transversion. A transition event involves the interchange of two-ring purines (A and G) or of one-ring pyrimidines (C and T). Transversion events the exchange of a purine for a pyrimidine or vice versa. The rate at which these two events occur is important to understanding of the evolution of a gene. Therefore, the rate of nucleotide substitution in NAC TFs was analysed. Results indicated that the rate of transition in NAC TFs is higher than the rate of transversion. The substitution of adenine with guanine was found to be highest in *Linum usitatissimum* (15.82), while the substitution of guanine to adenine was found to be the highest in *Lotus japonicas* (19.07). The lowest rate of substitution from adenine to guanine and vice versa was found in *Trifolium pratense* (9.73) and *Amborella trichopoda* (10.8), respectively (Table 4). The highest rate of substitution from thiamine to cytosine and vice versa was found in *Klebsormidium flaccidum* (7.19) and *Pseudotsuga menziesii* (11.59), respectively. The lowest rate of substitutions from thiamine to cytosine and vice versa was found in *Capsella grandiflora* (2.41) and *Cicer arietinum* (1.62), respectively (Table 4). These data make it evident that the rates of transition of purine (adenine and guanine) nucleotides are higher than the rates of pyrimidines. The highest rate of transversion from adenine to thiamine and vice versa was found in *Capsella grandiflora* (12.34 for adenine to thiamine and 9.91 for thiamine to adenine) (Table 4). The rate of substitution by transversion is slower relative to the rate of substitution by transition.

**Table 4.**
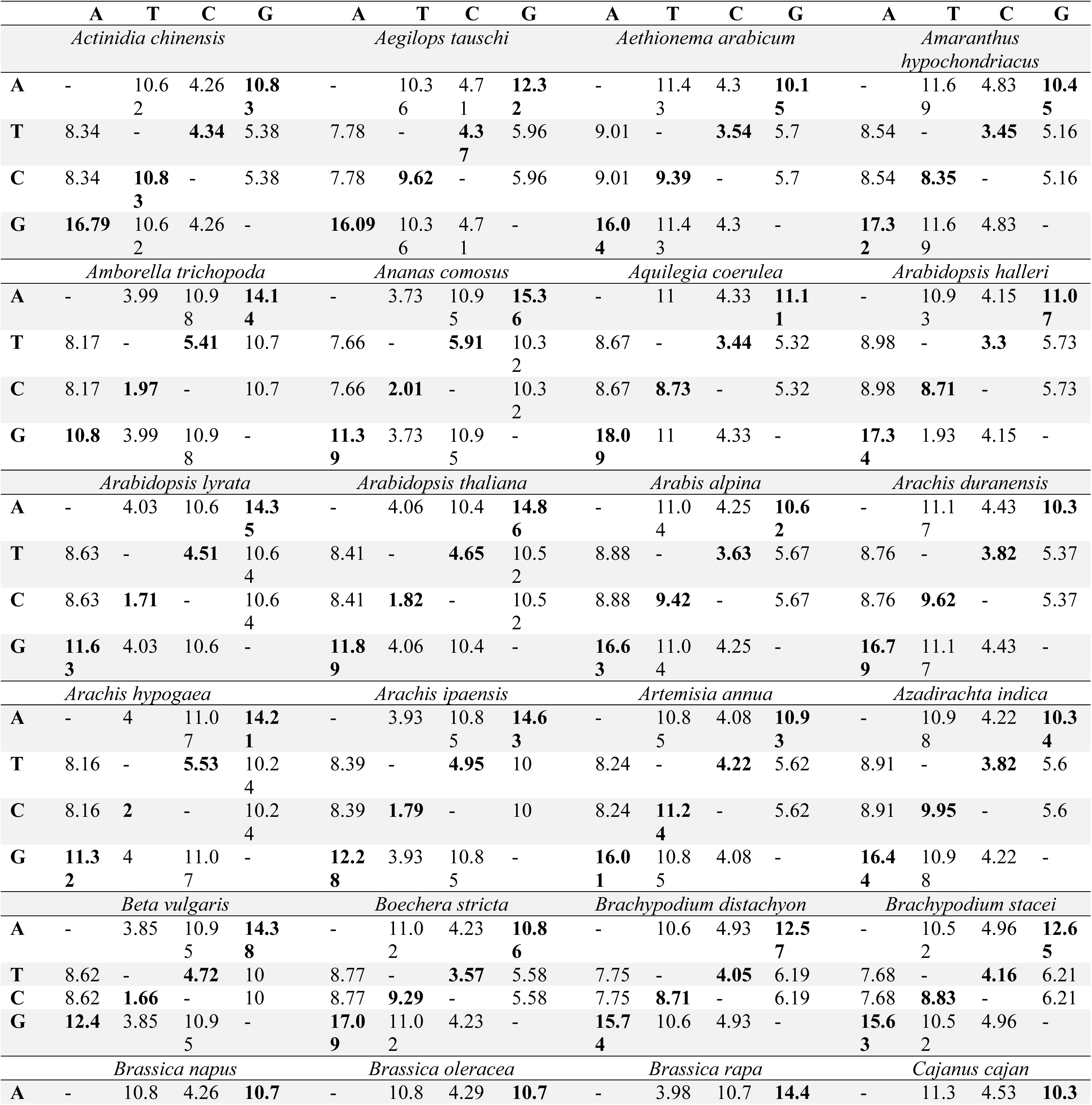
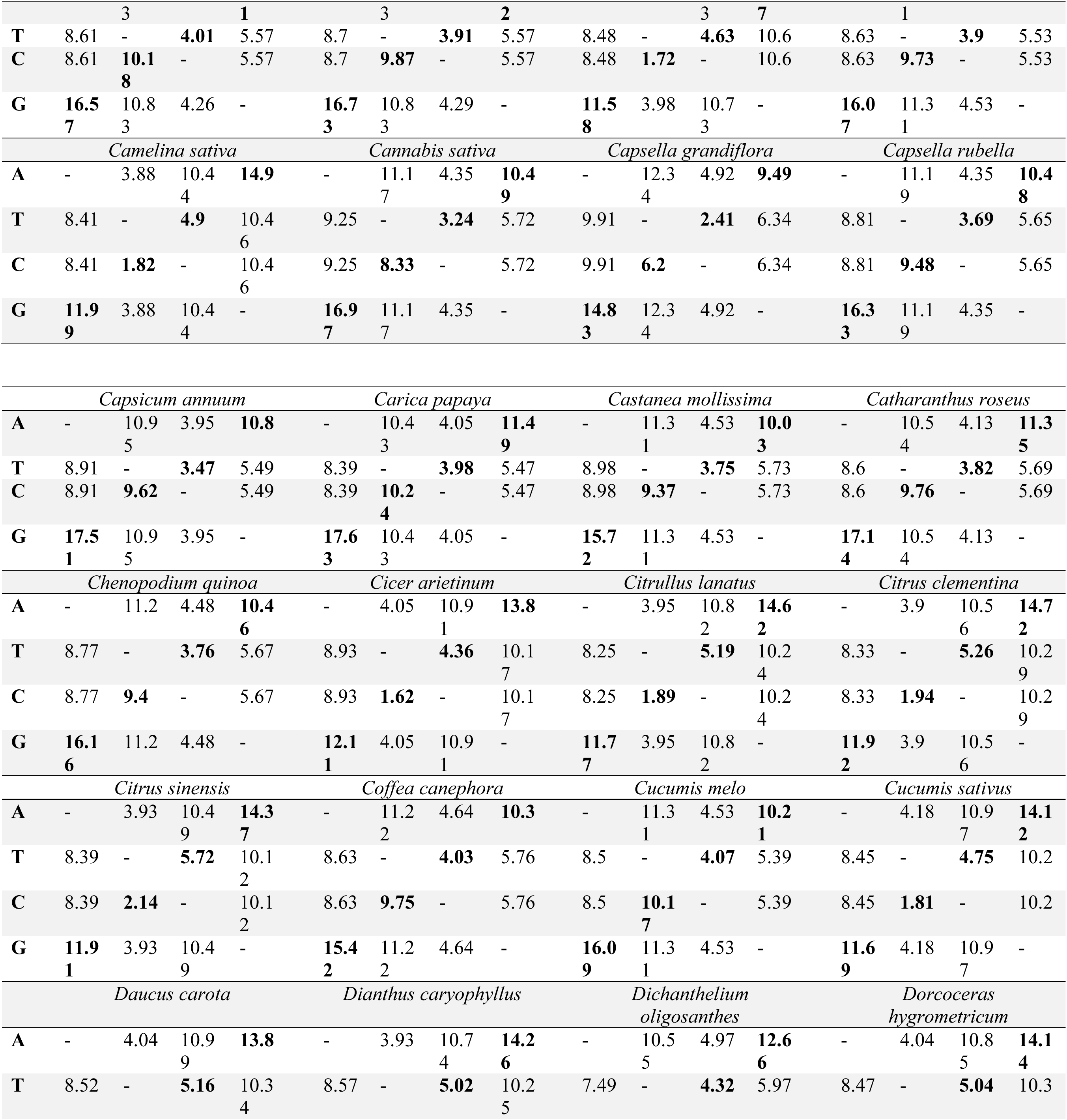
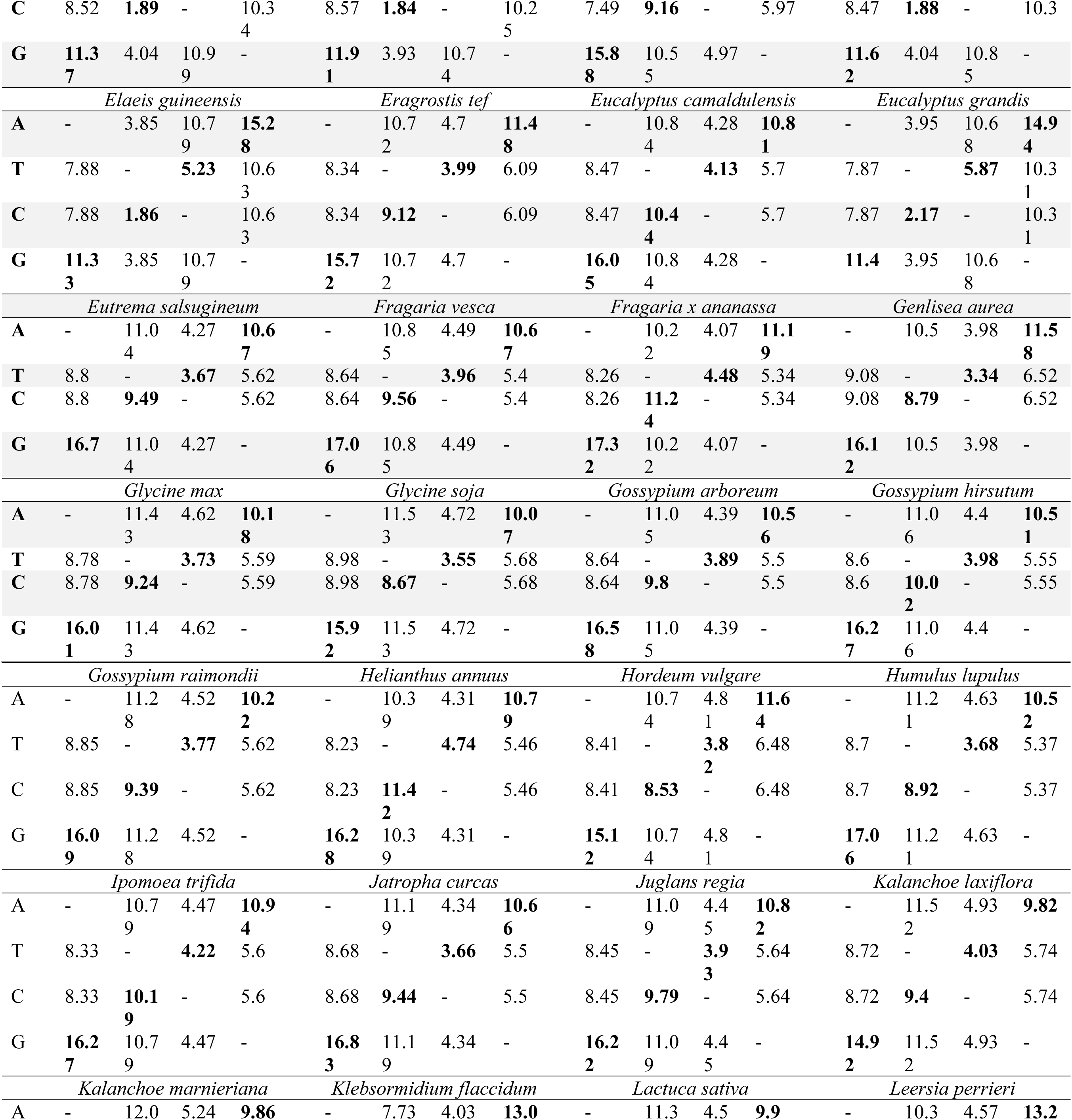
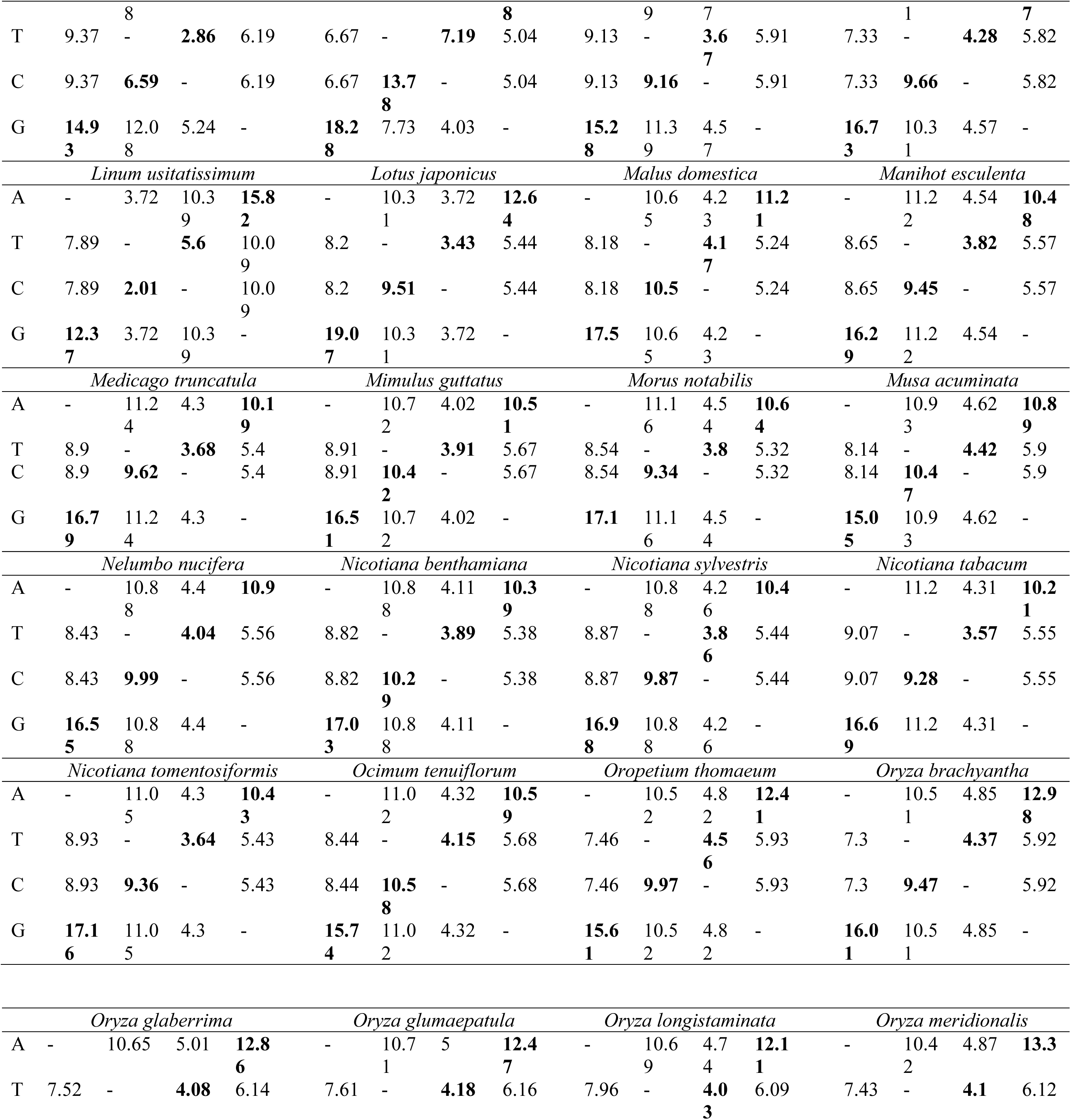
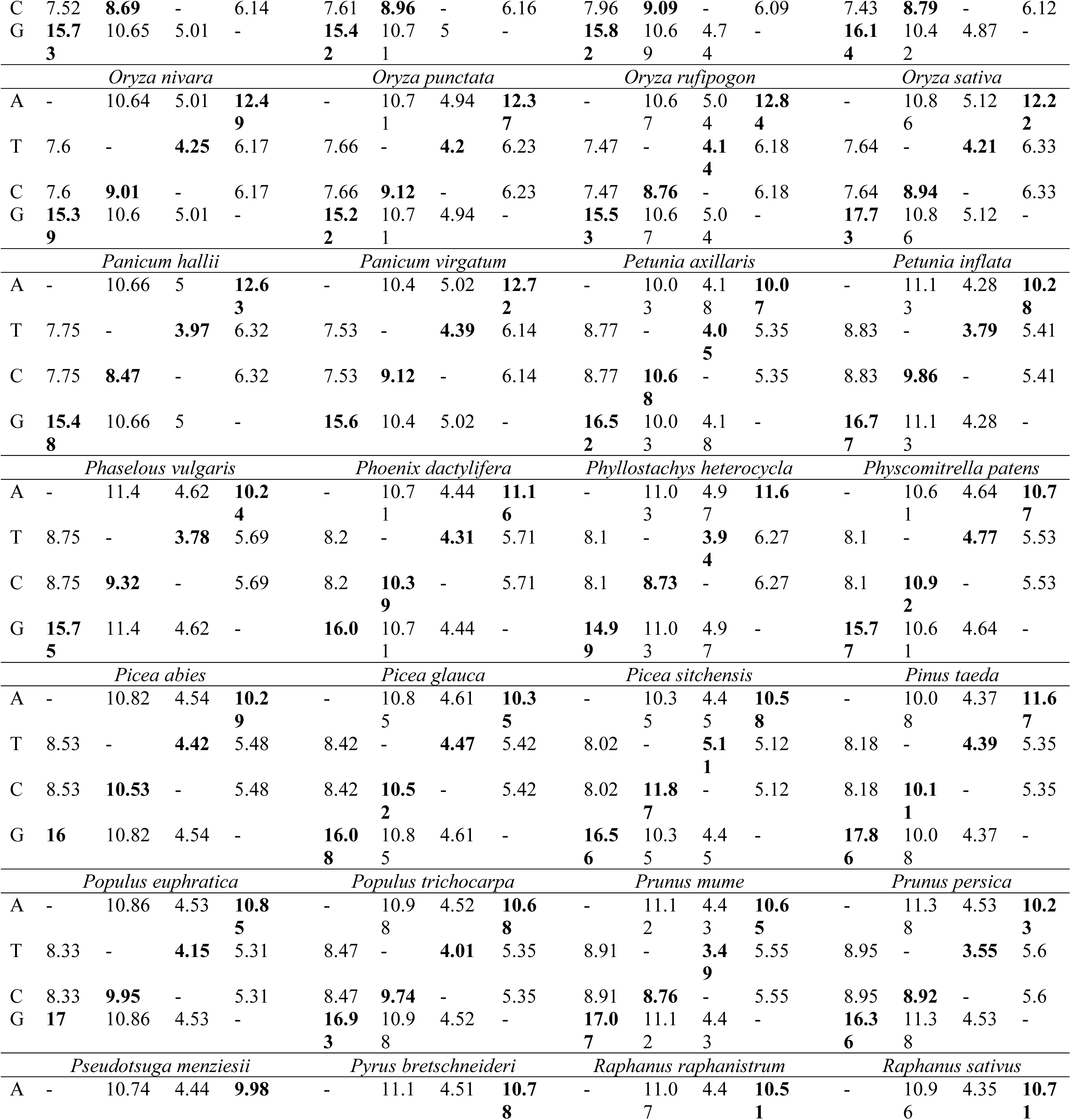
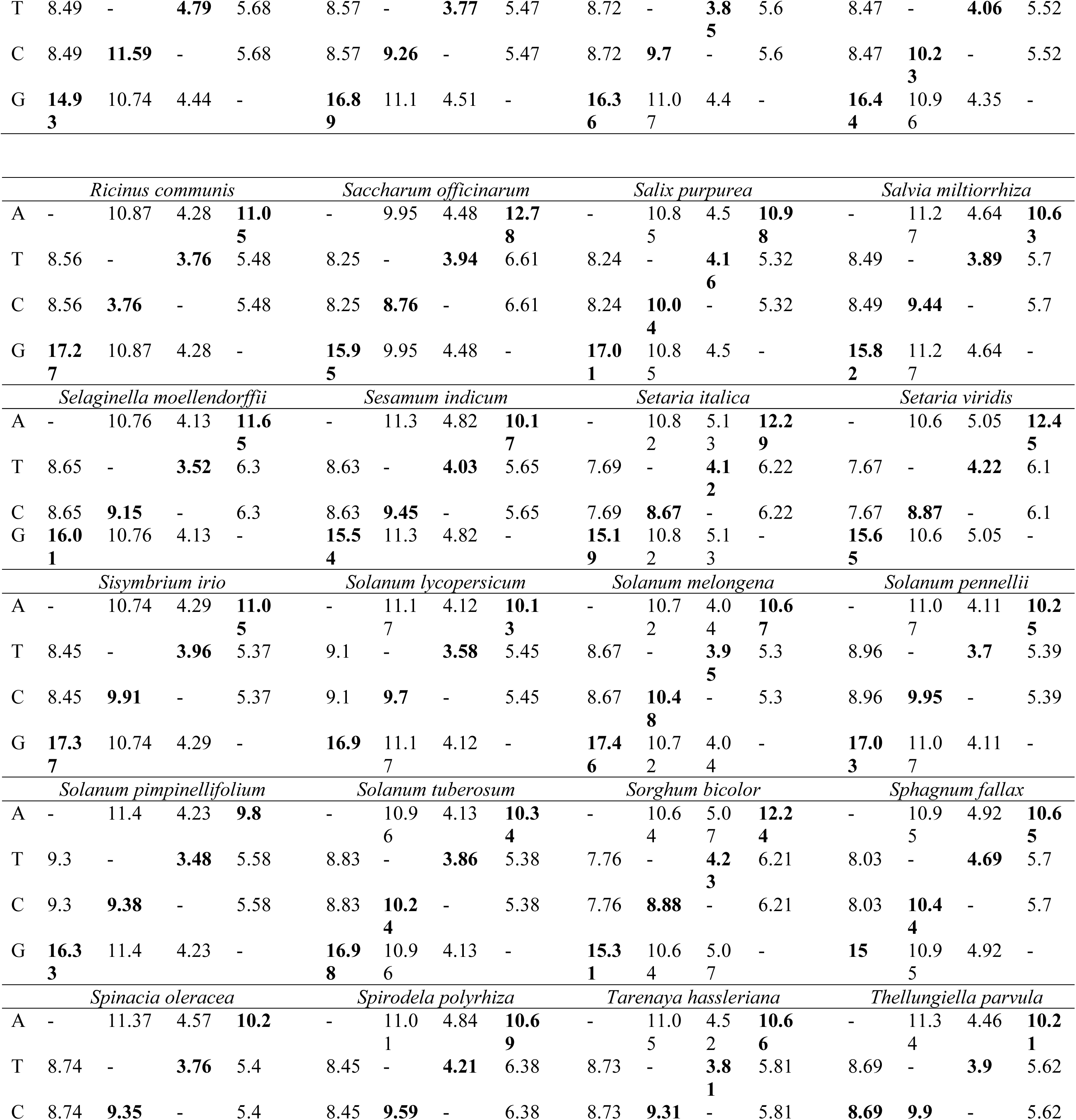
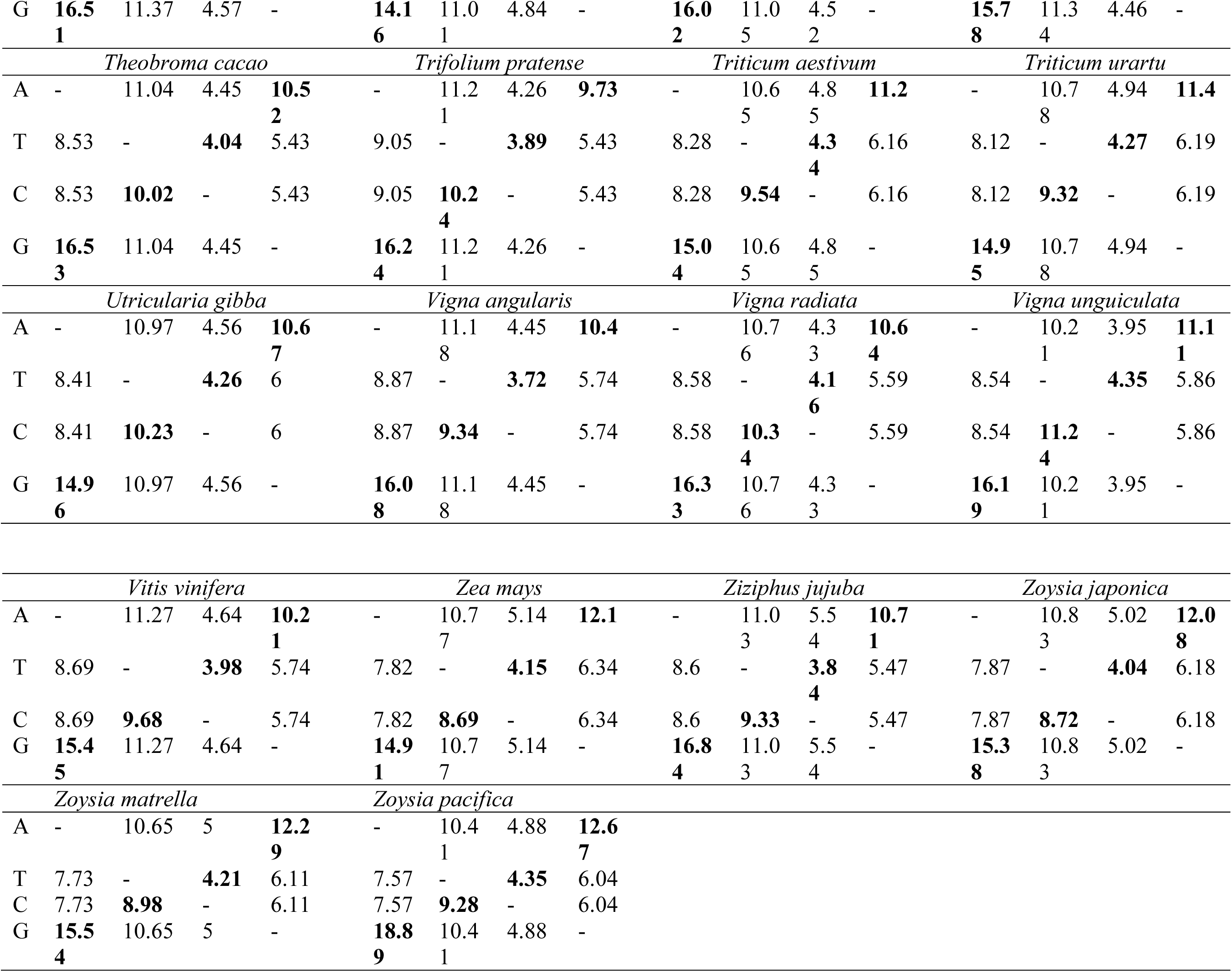
Substitution rate of NAC TFs of plants.

*Capsella grandiflora* is a close relative of *Arabidopsis thaliana* and is predicted to be the progenitor of *Capsella bursa-pastoris*. *Capsella grandiflora* is a self-pollinating plant and is used as a model organism in evolutionary studies and the change from self-incompatibility into self-compatibility. The genomic consequences of the evolution of selfing, however, is poorly understood. *Capsella rubella*, a close relative of *Capsella grandiflora*, that evolved self-compatibility 200,000 years ago ^62^ also exhibits a high rate of transversion from adenine to thiamine (11.19). Thus, the higher rate of transversion from adenine to thiamine in *Capsella grandiflora* and *Capsella rubella* may be a possible factor in the evolution of self-pollination. Higher rates of transversion were also found in *Solanum pimpinellifolium* (11.4) and *Castanea mollissima* ((11.31) Chinese chestnut). *Solanum pimpinellifolium* is self-pollinating and exhibits high levels of stress tolerance ^63^. *Castanea mollissima* has evolved over a period of time in coexistence with chestnut blight and is resistant to the pathogen. This indicates that higher rates of transversion from adenine to thiamine and vice versa are associated with self-pollination and stress tolerance in plants. The highest rate of substitution from guanine to cytosine and vice versa was found in *Arachis hypogaea* (11.07), and *Camelina sativa* (11.46), respectively (Table 4). The lowest rate of substitution from adenine to thiamine and vice versa was found in *Linum usitatissimum* (3.72) and *Klebsormidium flaccidum* (6.67), respectively. Notably, the highest rate of substitution from thiamine to cytosine was found in *Klebsormidium flaccidum* and the highest rate of substitution from adenine to guanine was found in *Linum usitatissimum*. This indicates that organisms which exhibit the highest rate of transition possess the lowest rate of transversion.

### NAC TFs evolved from orthologous ancestors

A phylogenetic tree of NAC TFs was constructed to understand their evolutionary relationships. A model selection was conducted before constructing the phylogenetic tree using the maximum likelihood statistical method. The phylogenetic tree revealed the presence of at least seven phylogenetic clustered orthologous (COGs) groups originating from a common, orthologous ancestor (Figure 7). Each phylogenetic cluster was further divided into two or more sub-groups. A phylogenetic tree of each individual species was subsequently constructed to examine duplication and loss events in NAC TFs. The phylogenetic tree of each species was independently reconciled with the collective species tree. This analysis indicated that NAC TFs in all of the species were duplicated and no NAC TFs were found to be lost. This suggest that NAC TFs evolved from common ancestors (orthology) and underwent numerous duplication events during speciation (paralogy), which gave rise to diverse gene functions in plant development and growth. We also checked for the presence of potential foreign or homologous sequences (xenologs) in NAC TFs. No primary xenologs, sibling donor xenologs, sibling recipient xenologs, incompatible xenologs, autoxenologs, or paraxenologs were identified in NAC TFs. Although the phylogenetic tree indicates the evolution NAC TFs from common ancestors, none of the NAC genes in the examined species were found to have been transferred from one species to another (Table 1). Previous studies of NAC TFs in six plant species also reported a high level of duplication and divergent evolution ^64^. The expansion of TF families was associated with an increase in the structural complexity of the organism ^65^. Previous studies reported the lineage-specific grouping of transcription factors ^54,64^. The phylogenetic tree of NAC TFs also revealed the presence of lineage-specific clustering as well. In a few cases, however, order-specific clustering of NAC TFs was also observed. For example, NAC TFs in dicot species of the Brassica lineage, including *A. thaliana, A. halleri, B. napus, B. rapa, R. sativus, R. raphanistrum, C. rubella, A. alpine*, and others, grouped together. Similarly, NAC TFs in monocot plant species, including *O. sativa*, *O. nivara, B. distachyon*, and others, also grouped together.

**Figure 7.**
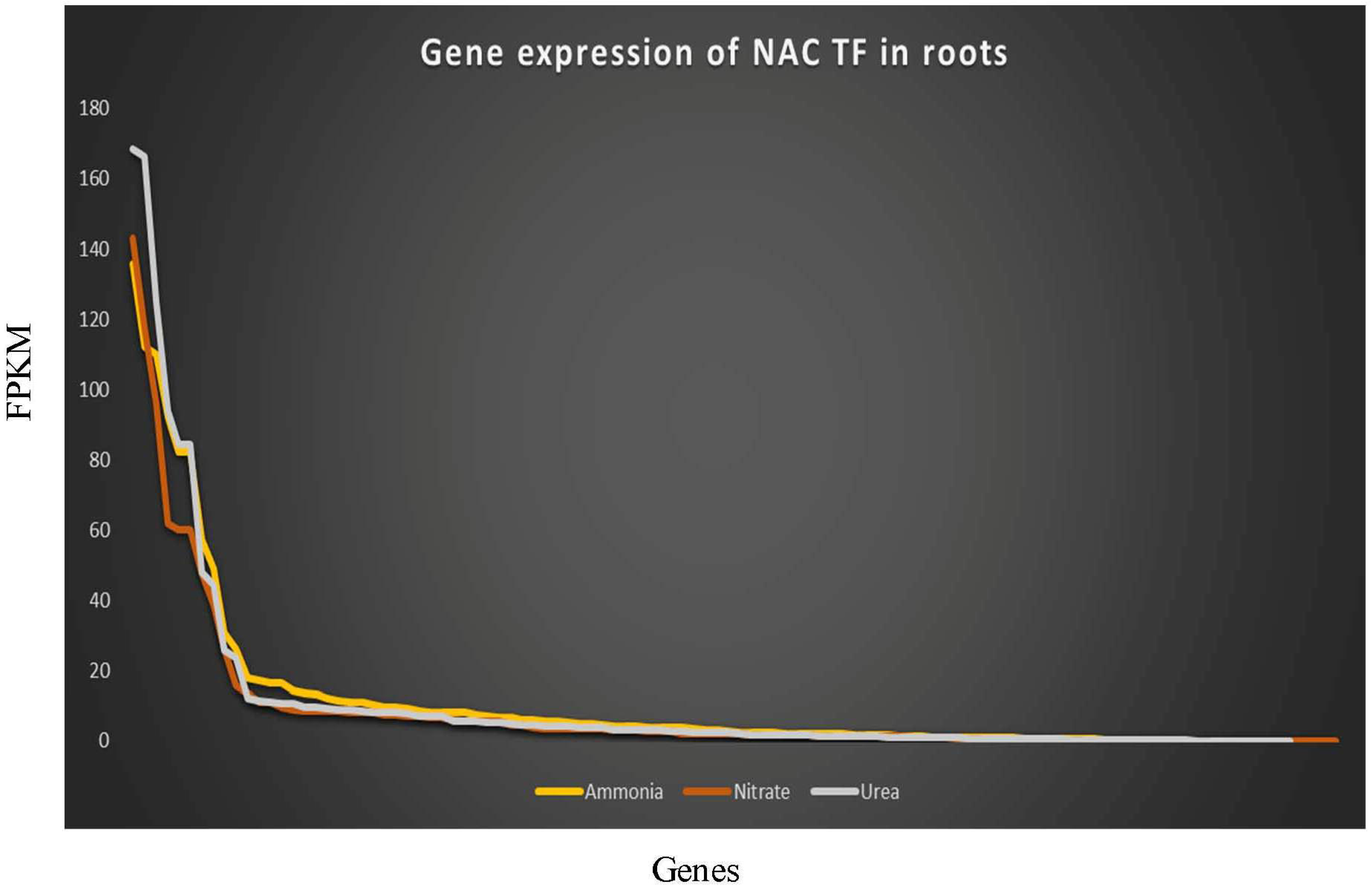
Differential expression of NAC TFs in roots of *A. thaliana* plants treated with ammonia, nitrate, and urea. The expression of *A. thaliana* NAC TFs was analysed to determine their response to different sources of nitrogen. Expression data were obtained from the PhytoMine database in Phytozome and presented as FPKM (Fragments per Kilobase of transcripts per million mapped reads).

**Figure 8.**
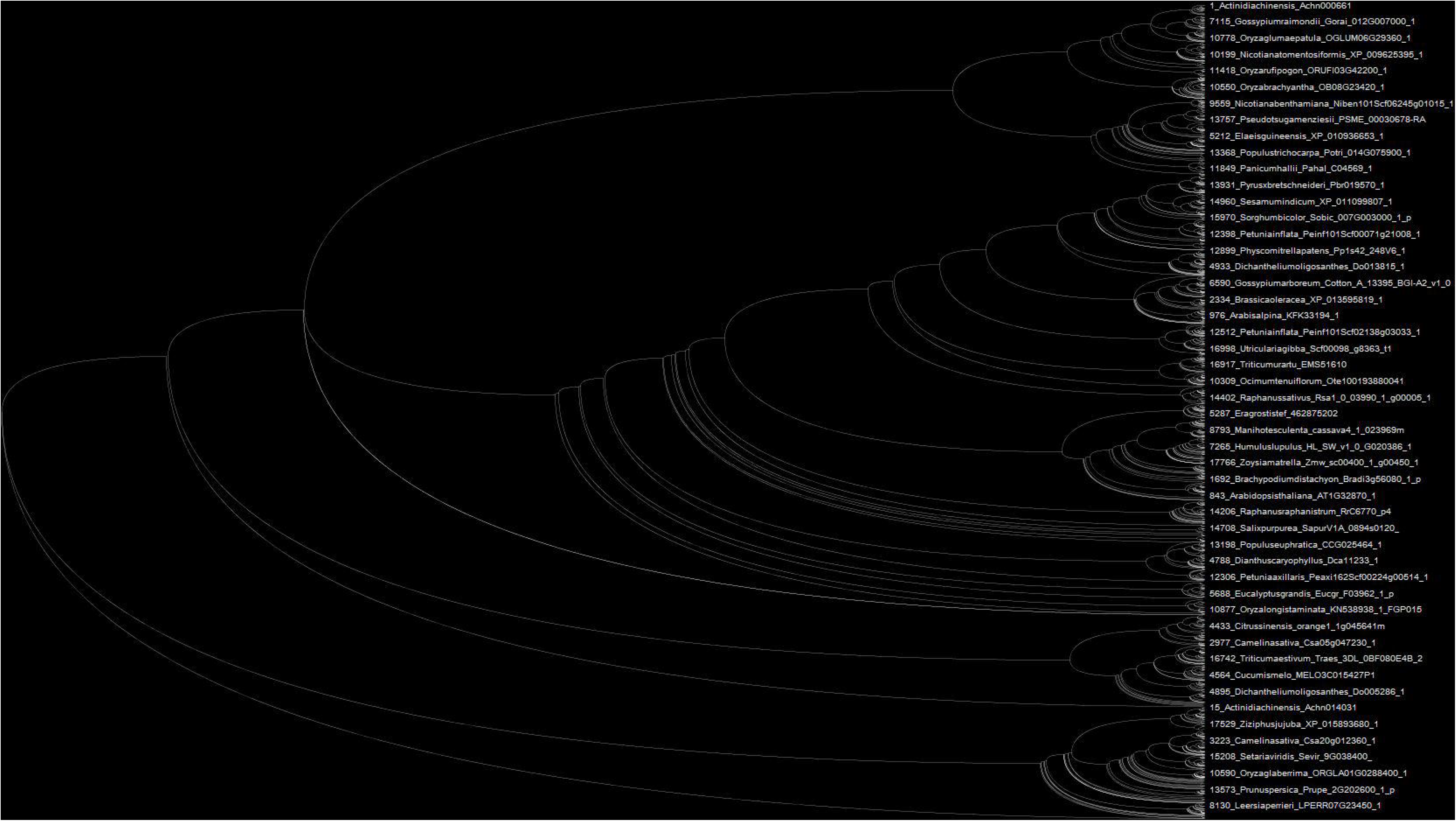
Phylogenetic tree of NAC TFs. A phylogenetic tree of NAC TF reveals the presence of seven clustered orthologous groups (COGs). Each group also possesses two or more sub-groups. The phylogenetic tree shows lineage (monocot/dicot) specific grouping of NAC TFs. The phylogenetic tree was constructed using the neighbor joining method with 1000 bootstrap replicates.

## Conclusion

NAC TFs are present in higher plants, as well as in a few species of algae. The number of NAC TFs per genome and their structural and functional properties increased with the complexity of the organism. The algae *Klebsormidium flaccidum*, a charophyte, was also found to possess NAC TFs; suggesting that the evolution of NAC TFs was associated with the adaptation of plant life from an aquatic to a terrestrial form. The paralogous evolution of NAC TFs underlies their diverse functional role in plant growth and development. Duplication events in NAC TFs were greater than deletion events and the absence of any loss of NAC TFs in different plant species indicates their evolution in recent times. As NAC TFs play a pivotal role within the nucleus regulating gene expression, the presence of bipartite and multipartite nuclear localization signals is of particular interest and provides the basis for further investigation of their functional roles.

## Materials and Methods

### Identification of NAC TFs

NAC genes in the studied plant species were obtained from searches in the National Centre for Biotechnology Information (NCBI), Phytozome, and Plant Genome databases ^66^. BLASTP and hidden Markov model were used to identify the NAC TFs in different species using AtNAC1 and AtBAC2 as the query sequences ^67^. Protein and CDS sequences of each species were collected and further analysed. Protein sequences of the NAC TFs were subjected to BLASTP analysis against a reference database to reconfirm them as a NAC TF of the identified species. All of the NAC TF protein sequences in the examined species were also subjected to ScanProsite and InterPro scans to confirm the presence of a NAC domain ^68,69^. Sequences that were found to contain a NAC domain were considered as NAC TFs. The presence of multiple NAC domains, along with the presence of chimeric NAC domains, were determined through ScanProsite and InterPro scans. The presence of multiple functional sites in NAC TFs were also analysed using ScanProsite software.

### Analysis of membrane attachment and nuclear localization signal sequences

The presence of transmembrane domains in NAC TFs of all of the examined species were identified using the TMHMM server v. 2.0 ^70^ and default parameters. Nuclear localization signal sequences in NAC TFs were identified using NLStradamus software, which uses a hidden Markov model for the prediction of nuclear localization signals ^71^. NAC TF protein sequences were uploaded in FASTA format to run the program. The parameters used to run the NLS analysis were; HMM state emission and transition frequencies, 2 state HMM static; prediction type Viterbi and posterior, prediction cut-off 0.4; prediction display, and image and graphic.

### Interactome analysis of NAC TFs

*A. thaliana* NAC TFs were used to examine the complex interactome network of NAC TFs. The individual interaction network of each NAC TF in *A. thaliana* was searched in a string database that contains 9.6 million proteins from 2031 organisms ^72,73^. The interactome network of each of NAC TF were noted and the results were later used to construct the interactome network of *A. thaliana* NAC TFs. The presented interactome network was based on an experimentally validated network, co-expressed network, and a mined network. These outputs were used to construct the interactome network. The NAC TFs used to construct the interactome network were subjected to GO (gene ontology) and cellular process analyses.

### Gene expression analysis

Differential gene expression of NAC TFs was analysed to elucidate their role in growth, development, and nitrogen assimilation. *A. thaliana* NAC TFs were used to examine differential gene expression. Transcriptome data from *A. thaliana* treated with ammonia, nitrate, and urea were utilized from the PhytoMine database in Phytozome. The expression pattern of NAC TFs for leaf and root tissues in the treated *A. thaliana* plants were analysed separately. The expression was measured in fragments per kilobase of exon per million fragments mapped (FPKM). Transcripts with a zero value were discarded from the study.

### Construction of a phylogenetic tree

Two approaches were used to construct phylogenetic trees. In the first approach, a phylogenetic tree was constructed using the NAC TFs of individual species. In the second approach, the NAC TFs of all of the examined species were used to construct a phylogenetic tree. The phylogenetic tree for individual species was constructed to determine the deletion and duplication events in NAC TFs within individual species. Prior to construction of the phylogenetic trees, a model selection was carried out in MEGA6 software. The following parameters were used in the model, analysis, model selection; tree to use, automatic (neighbor joining), statistical method, maximum likelihood; substitution type, nucleotides; gaps/missing data treatment, partial deletion; site coverage cut-off (%), 95; codons included, 1^st^+2^nd^+3^rd^+non-coding. Based on the lowest BIC values of model selection, phylogenetic trees of NAC TFs were carried out using the neighbor joining method, a GTR statistical model, and 1000 bootstrap replicates.

### Analysis of transition and transversion rates

Transition and transversion rates in NAC TFs within individual species were analysed using MEGA6 software. The converted MEGA file format of individual species was used to determine the rate of transition and transversion. The following statistical parameters were used to study the transition/transversion rate: estimate transition/transversion bias; maximum composite likelihood estimates of the pattern of nucleotide substitution; substitution type, nucleotides; model/method, Tamura-Nei; gaps/missing data treatment, pairwise deletion; codon position, 1^st^, 2^nd^, 3^rd^, and non-coding sites.

### Analysis of gene deletion and duplication

Prior to the analysis of deletion and duplication events in NAC TFs, a species tree was constructed in the NCBI taxonomy browser (https://www.ncbi.nlm.nih.gov/Taxonomy/CommonTree/wwwcmt.cgi). All of the studied species were used to construct the species tree. The resulting phylogenetic trees of individual species in a nwk file format were uploaded in Notung2.9 software as a gene tree and reconciled as a gene tree with the species tree to obtain duplicated and deleted genes. Deletion and duplication events were analysed in all of the studied species individually.

## Data availability

All the data used during this study was taken from publicly available genomic databases and details are mentioned in the materials and methods section.

## Author contributions

TKM: conceived the idea, performed the experiments and analysis, drafted and revised the manuscript, AK: performed the analysis, DY: revised the manuscript, AH: drafted and revised the manuscript, BT: revised the manuscript, ALK: analysed the data and revised the manuscript, EFA: revised the manuscript, AAH: revised the manuscript.

## Supplementary Data

**Supplementary Table 1** Supplementary table showing different chimeric domains of NAC TFs.

**Supplementary Table 2** NAC TFs showing the presence of novel functional domain along with NAC domains.

**Supplementary Table 3** NAC TFs showing their involvement in different pathways and biological process.

