## Supplementary Table 1 for "Genomics, Molecular and Evolutionary Perspective of NAC Transcription Factors"

Supplementary table showing different chimeric domains of NAC TFs.

| **Domains** | | | | | **Gene ID** | | | | | **Domain architecture** | |
| --- | --- | --- | --- | --- | --- | --- | --- | --- | --- | --- | --- |
| ***Actinidia chinensis*** | | | | | | | | | | | |
| Double domain | | | | | ACHN181121 | | | | | 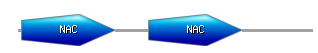 | |
| FE20G_Oxy+NAC | | | | | ACHN004841 | | | | | 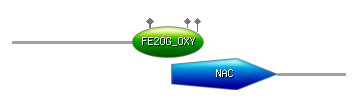 | |
| EF+NAC | | | | | ACHN274891 | | | | | 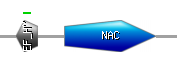 | |
| Kinase+NAC+CRM | | | | | ACHN089081 | | | | | 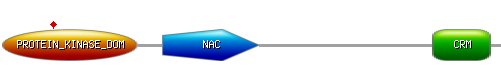 | |
| Kinase+NAC | | | | | ACHN335281  ACHN389881 | | | | | 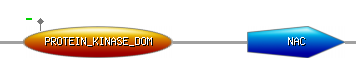 | |
| ***Amaranthas hypochondriacus*** | | | | | | | | | | | |
| Double domain | | | | | AHYPO_009690 | | | | |  | |
| ***Aegilops tauschii*** | | | | | | | | | | | |
| IQ+NAC | | | | | EMT05508 | | | | | 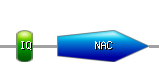 | |
| FBOX+NAC | | | | | EMT14186 | | | | | 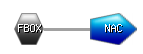 | |
| DNAJ+NAC+ZF_B | | | | | EMT28787 | | | | | 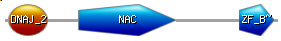 | |
| NAC+Kinase | | | | | EMT33859 | | | | |  | |
| ***Ananas comosus*** | | | | | | | | | | | |
| NAC+PPR Repeat | | | | | Aco018846 | | | | | 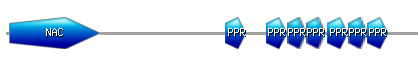 | |
| ***Arabidopsis lyrata*** | | | | | | | | | | | |
| 4 NAC domains | | | | | 338342 | | | | | 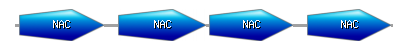 | |
| 2 NAC domains | | | | | 347836  475292  893515  893532 | | | | |  | |
| *Arabidopsis thaliana* | | | | | | | | | | | |
| Double domain | | | | | AT1G60340.1  AT1G60350.1  AT1G60380.1  AT1G60300.1  AT1G60280.1 | | | | |  | |
| *Brachypodium distachyon* | | | | | | | | | | | |
| NAC+ZF_B | | | | | Bradi4g03620.1 | | | | | 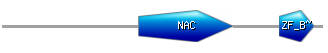 | |
| Double domain | | | | | Bradi2g46430.1  Bradi5g15230.1 | | | | |  | |
| *Brassica napus* | | | | | | | | | | | |
| Double domain | | GSBRNA2T00013424001  GSBRNA2T00020965001  GSBRNA2T00026805001  GSBRNA2T00026833001  GSBRNA2T00030633001  GSBRNA2T00040675001  GSBRNA2T00042092001  GSBRNA2T00050907001  GSBRNA2T00095486001  GSBRNA2T00095503001 | | | | | | | |  | |
| NAC+TIR | | GSBRNA2T00021663001 | | | | | | | |  | |
| NAC+ENT | | GSBRNA2T00050859001  GSBRNA2T00098817001  GSBRNA2T00155943001 | | | | | | | |  | |
| NAC+PPC | | GSBRNA2T00075489001  GSBRNA2T00139105001 | | | | | | | | 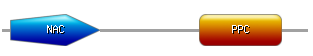 | |
| NAC+CYT_B561 | | GSBRNA2T00098300001 | | | | | | | | 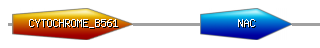 | |
| *Brassica rapa* | | | | | | | | | | | |
| Double domain | | | | | Bra002857  Bra035412  Bra029571  Bra031518 | | | | |  | |
| NAC+ENT | | | | | Bra003244 | | | | | 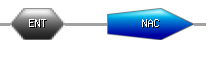 | |
| NAC+DFDF+DFDF+CYT B | | | | | Bra012470 | | | | | 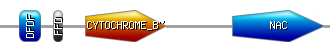 | |
| *Capsella grandiflora* | | | | | | | | | | | |
| Double domain | | | | | Cagra.1301s0009  Cagra.1301s0010 | | | | |  | |
| *Capsella rubella* | | | | | | | | | | | |
| Double domain | | | | | Carubv10008035m  Carubv10008036m  Carubv10021319m  Carubv10021818m  Carubv10022099m | | | | |  | |
| *Chenopodium quinoa* | | | | | | | | | | | |
| NAC+ABC_TM1F | | | | | AUR62022690 | | | | | 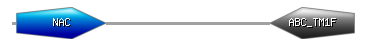 | |
| *Citrus sinensis* | | | | | | | | | | | |
| Double domain | | | | | orange1.1g044381m  orange1.1g045641m | | | | |  | |
| *Daucus carota* | | | | | | | | | | | |
| NAC+CRM | | | | | DCAR_021729 | | | | | 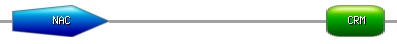 | |
| NAC+RWP+PB1 | | | | | DCAR_030911 | | | | | 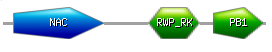 | |
| *Fragaria vesca* | | | | | | | | | | | |
| Double domain | | | | | mrna17720.1-v1.0-hybrid  mrna17721.1-v1.0-hybrid  mrna19382.1-v1.0-hybrid | | | | | |  |
| NAC+ZF_P | | | | | mrna04303.1-v1.0-hybrid | | | | | | 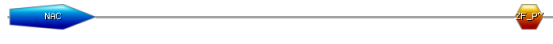 |
| NAC+PABC | | | | | mrna09127.1-v1.0-hybrid | | | | | | 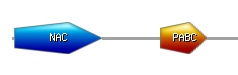 |
| NAC+EF | | | | | mrna23394.1-v1.0-hybrid | | | | | | 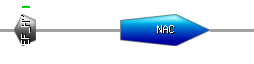 |
| NAC+Peptidase_A1 | | | | | mrna24179.1-v1.0-hybrid | | | | | | 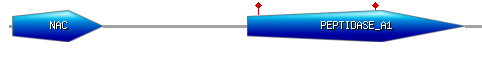 |
| NAC+Kinase | | | | | mrna27940.1-v1.0-hybrid | | | | | | 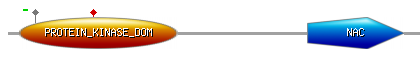 |
| NAC+RESPO | | | | | mrna31175.1-v1.0-hybrid | | | | | | 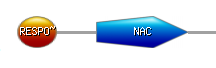 |
| *Jatropha curcas* | | | | | | | | | | | |
| NAC+Integrase | | | | | Jcr4S16185.10 | | | | | | 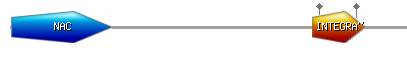 |
| *Lotus japonicus* | | | | | | | | | | | |
| Double domain | | | | | chr1.CM0207.190.r2.m  chr3.CM0208.570.r2.m | | | | | |  |
| *Linum usitatissimum* | | | | | | | | | | | |
| Double domain | | | | | Lus10002083 | | | | | |  |
| NAC+JMJN+JMJC | | | | | Lus10003548 | | | | | | 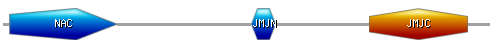 |
| *Malus domestica* | | | | | | | | | | | |
| Double domain | | | | | MDP0000289955  MDP0000689991 | | | | | |  |
| NCA+TIR+LRR+CS | | | | | MDP0000180046 | | | | | | 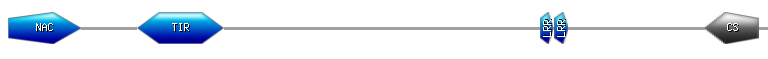 |
| NAC+SAM | | | | | MDP0000239596 | | | | | | 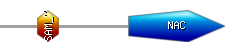 |
| NAC+Kinase | | | | | MDP0000276278 | | | | | | 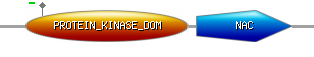 |
| NAC+BRX | | | | | MDP0000276982 | | | | | | 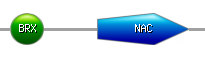 |
| NAC+FBox+Kinase | | | | | MDP0000283975 | | | | | | 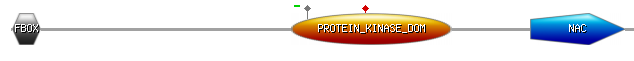 |
| NAC+WD Repeats | | | | | MDP0000288624 | | | | | | 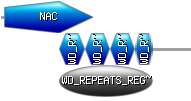 |
| NAC+Kinase | | | | | MDP0000309351 | | | | | | 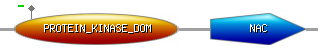 |
| NAC+G_TR_2 | | | | | MDP0000428265 | | | | | | 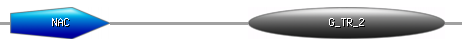 |
| NAC+CRM | | | | | MDP0000663564 | | | | | | 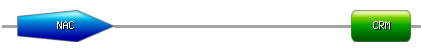 |
| *Medicago truncatula* | | | | | | | | | | | |
| Double domain | | | | | AC233657_10.1 | | | | | |  |
| *Musa acuminata* | | | | | | | | | | | |
| Double domain | | | | | GSMUA_Achr9P04960_001 | | | | | |  |
| NAC+Myosin+IQ+Dilute | | | | | GSMUA_Achr4P07150_001 | | | | | | 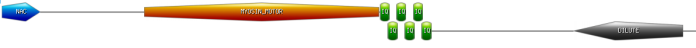 |
| *Oropetium thomaeum* | | | | | | | | | | | |
| Double domain | | | | | Oropetium_20150105_15183 | | | | | |  |
| *Oryza barthii* | | | | | | | | | | | |
| NAC+ZF_Ring_2 | | | | | ObartAA03S_FGP0027 | | | | | | 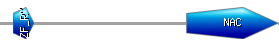 |
| NAC+Pentatricopeptide | | | | | ObartAA03S_FGP21921 | | | | | | 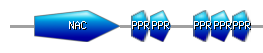 |
| NAC+DNAJ_2 | | | | | ObartAA03S_FGP31756 | | | | | | 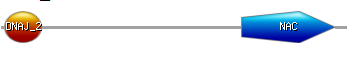 |
| NAC+WRKY | | | | | ObartAA03S_FGP8394 | | | | | | 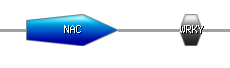 |
| *Oryza brachyantha* | | | | | | | | | | | |
| NAC+CHCH | | | | | OB04G21050.1 | | | | | | 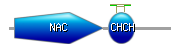 |
| Double domain | | | | | OB11G21460.1  OB03G25660.1 | | | | | |  |
| *Oryza glaberrima* | | | | | | | | | | | |
| Double domain | | | | | ORGLA04G0153400.1 | | | | | |  |
| *Oryza punctata* | | | | | | | | | | | |
| Double domain | | | | | OpuncBB_FGP18839 | | | | | |  |
| NAC+Pentatricopeptide | | | | | OpuncBB_FGP18634 | | | | | | 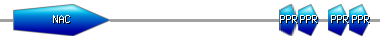 |
| *Oryza sativa* | | | | | | | | | | | |
| Double domain | | | | | LOC_Os04g42940.1 | | | | | |  |
| *Panicum hallii* | | | | | | | | | | | |
| Double domain | | | | | Pahal.A02025  Pahal.D02537 | | | | | |  |
| NAC+ZF_B, NAC+ZF_B | | | | | Pahal.C04574 | | | | | | 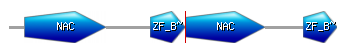 |
| *Panicum virgatum* | | | | | | | | | | | |
| Double domain | | | | | Pavir.Bb00848  Pavir.Db01755  Pavir.Db01806  Pavir.J34688 | | | | | |  |
| *Phoenix dactylifera* | | | | | | | | | | | |
| Double domain | | | | | PDK_30s698051g001  PDK_30s758201g003  PDK_30s830451g001 | | | | | |  |
| NAC+Pentatricopeptide | | | | | 30s1165641g010 | | | | | | 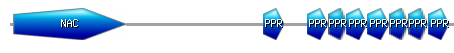 |
| *Phyllostachys heterocycla* | | | | | | | | | | | |
| Double domain | | | | | PH01000804G0640  PH01001619G0180 | | | | | |  |
| NAC+Pentatricopeptide | | | | | PH01000020G0110 | | | | | |  |
| NAC+ZF_C | | | | | PH01000143G0710 | | | | | | 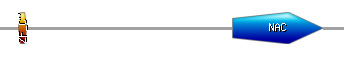 |
| *Picea abies* | | | | | | | | | | | |
| Double domain | | | | | MA_10431997g0010 | | | | | |  |
| *Populous trichocarpa* | | | | | | | | | | | |
| NAC+RDRP | | | | | Potri.006G051400.1 | | | | | | 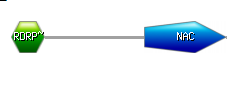 |
| *Prunus persica* | | | | | | | | | | | |
| Double domain | | | | | Prupe.1G284800.1.p | | | | | |  |
| NAC+FBox | | | | | Prupe.4G112200.1.p | | | | | |  |
| *Pyrus bretschneideri* | | | | | | | | | | | |
| Double domain | | | | | Pbr019214.6 | | | | | |  |
| NAC+BIOTINYL+LIPOYL | | | | | Pbr004711.1 | | | | | | 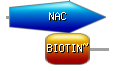 |
| NAC+TIR | | | | | Pbr009270.1 | | | | | | 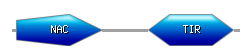 |
| NAC+CRM | | | | | Pbr012547.2 | | | | | | 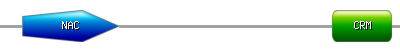 |
| NAC+Kinase+CRM | | | | | Pbr014522.2 | | | | | | 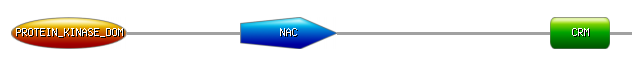 |
| NAC+ZF_RI | | | | | Pbr038615.1 | | | | | | 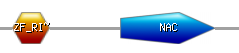 |
| *Raphanus sativus* | | | | | | | | | | | |
| Double domain | | | | | Rsa019644 | | | | | |  |
| *Setaria italica* | | | | | | | | | | | |
| Double domain | | | | | Si008305m  Si011665m  Si019248m  Si019775m | | | | | |  |
| *Setaria viridis* | | | | | | | | | | | |
| Double domain | | | | | Sevir.1G165700 | | | | | |  |
| *Solanum tuberosum* | | | | | | | | | | | |
| Double domain | | | | | PGSC0003DMP400054265 | | | | | |  |
| *Sorghum biolor* | | | | | | | | | | | |
| Double domain | | | | | Sobic.006G141900.1.p | | | | | |  |
| *Eutrema salsugineum* | | | | | | | | | | | |
| Double domain | | | | | Thhalv10015599m  Thhalv10024006m | | | | | |  |
| *Thellungiella parvula* | | | | | | | | | | | |
| Double domain | | | | | Tp2g04430 | | | | | |  |
| *Trifolium pratense* | | | | | | | | | | | |
| Double domain | | | | Tp57577_TGAC_v2_gene15980  Tp57577_TGAC_v2_gene30270 | | | | | | |  |
| NAC+PI3_4_Kinase | | | | Tp57577_TGAC_v2_gene37364 | | | | | | | 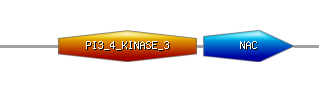 |
| *Triticum aestivum* | | | | | | | | | | | |
| Double domain | | | | Traes_2BL_75FBE2E45.1  Traes3BF073600060CFD_t1 | | | | | | |  |
| NAC+TSNARE | | | | Traes3BF073600100CFD_t1 | | | | | | |  |
| NAC+DNAJ_2+ZF_B | | | | Traes_1DS_4A2001BAC.1 | | | | | | | 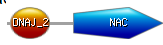 |
| *Triticum urartu* | | | | | | | | | | | |
| NAC+ZF_B | | | | EMS48536 | | | | | | |  |
| *Utricularia gibba* | | | | | | | | | | | |
| NAC+EF | | | | Scf00024.g3185.t1 | | | | | | | 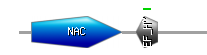 |
| *Vitis vinifera* | | | | | | | | | | | |
| Double domain | | | | GSVIVT01027475001 | | | | | | |  |
| *Zea mays* | | | | | | | | | | | |
| Double domain | | | | | | | | AC208663.3_FGP002 | | |  |
| NAc+Pentatricopeptide | | | | | | | | GRMZM2G312201_P01 | | |  |
| *Zostera marina* | | | | | | | | | | | |
| Double domain | | | | | | | | Zosma250g00090 | | |  |
| *Aethionema arabicum* | | | | | | | | | | | |
| Double domain | | | | | | | | AA10G00196  AA26G00262  AA93G00231 | | |  |
| *Arabidopsis halleri* | | | | | | | | | | | |
| Double domain | | | | | | | | Araha.16028s0003.1.p  Araha.61839s0001.1.p | | |  |
| *Arabis alpina* | | | | | | | | | | | |
| Double domain | | | | | | | | KFK42489.1 | | |  |
| *Boechera stricta* | | | | | | | | | | | |
| Double domain | | | | | | | | Bostr.10040s0190.1.p  Bostr.10040s0202.1.p | | |  |
| *Brachypodium stacei* | | | | | | | | | | | |
| Triple NAC domain | | | | | | | | Brast09G117600.1.p | | | 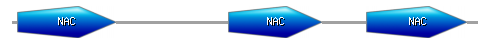 |
| Double domain | | | | | | | | Brast09G138900.1.p | | |  |
| *Brassica oleracea* | | | | | | | | | | | |
| Double domain | | | | | | | | XP_013594418.1  XP_013608221.1  XP_013610671.1  XP_013632799.1 | | |  |
| NAC+ENT | | | | | | | | XP_013589340.1  XP_013601975.1  XP_009104009.1 | | |  |
| *Camelina sativa* | | | | | | | | | | | |
| Double domain | | | | | | | | Csa01g011380.1  Csa03g002220.1  Csa03g037070.1  Csa04g065900.1  Csa05g047220.1  Csa05g047230.1  Csa06g007190.1  Csa08g006010.1  Csa08g060070.1  Csa09g013080.1  Csa14g002180.1  Csa14g002590.1  Csa15g003940.1  Csa17g001330.1  Csa17g001480.1  Csa17g097090.1  Csa20g023680.1 | | |  |
| Triple domain | | | | | | | | Csa09g081160.1  Csa14g044030.1 | | | 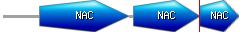 |
| Quadruple domain | | | | | | | | Csa16g052260.1 | | | 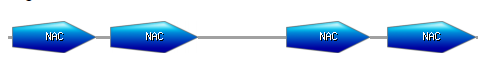 |
| *Castanea mollissima* | | | | | | | | | | | |
| Double domain | | | scaffold04550-snap-gene-0.24  scaffold04583-augustus-gene-0.13  scaffold07067-augustus-gene-0.6  scaffold08137-snap-gene-0.8 | | | | | | | |  |
| *Catharanthus roseus* | | | | | | | | | | | |
| NAC+Tricopeptide | | | cra_locus_1368_iso_3  cra_locus_1368_iso_4 | | | | | | | | 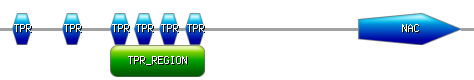 |
| *Dichanthelium oligosanthes* | | | | | | | | | | | |
| Double domain | | | Do002294.1  Do002300.1  Do009067.1  Do010830.1  Do011477.1  Do015402.1  Do015782.1  Do004998.1 | | | | | | | |  |
| NAC+PPR | | | Do020978.1 | | | | | | | |  |
| FBox+NAC+FBox | | | Do022798.1 | | | | | | | | 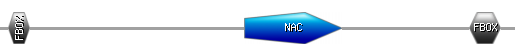 |
| [*Dorcoceras hygrometricum*](http://planttfdb.cbi.pku.edu.cn/index.php?sp=Dhy) | | | | | | | | | | | |
| NAC+JEF_N | | | KZV39672.1 | | | | | | | | 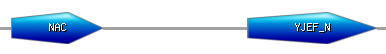 |
| NAC+Kinase | | | KZV46422.1 | | | | | | | |  |
| [*Elaeis guineensis*](http://planttfdb.cbi.pku.edu.cn/index.php?sp=Egu) | | | | | | | | | | | |
| Double domain | | | XP_010927165.1  XP_010904551.1 | | | | | | | |  |
| NAC+4FE4S | | | XP_010916182.1 | | | | | | | | 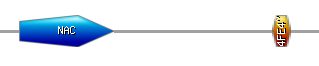 |
| *Eragrostis tef* | | | | | | | | | | | |
| Double domain | | | 462877018  462898851  462914386  462914484  462923579  462925604  462939746  462963614 | | | | | | | |  |
| NAC+Kinase | | | 462886127 | | | | | | | |  |
| NAC+HTH | | | 462890696 | | | | | | | | 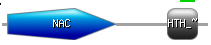 |
| 4 NAC domains | | | 462951506 | | | | | | | | 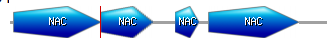 |
| *Eutrema salsugineum* | | | | | | | | | | | |
| Double domain | | | Thhalv10015599m  Thhalv10024006m | | | | | | | |  |
| *Fragaria ananassa* | | | | | | | | | | | |
| Double domain | FANhyb_icon00002021_a.1.g00001.1  FANhyb_rscf00000878.1.g00001.1 | | | | | | | | | |  |
| NAC+EF | FANhyb_rscf00001150.1.g00006.1 | | | | | | | | | | 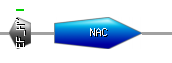 |
| *Genlisea aurea* | | | | | | | | | | | |
| NAC+Peroxidase 4 | | | | | | EPS67213.1 | | | | | 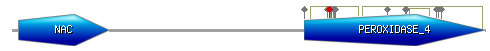 |
| *Glycine soja* | | | | | | | | | | | |
| NAC+LONGIN+V_SNA | | | | | | KHN21163.1 | | | | | 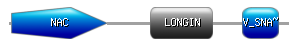 |
| [*Gossypium hirsutum*](http://planttfdb.cbi.pku.edu.cn/index.php?sp=Ghi) | | | | | | | | | | | |
| Double domain | | | | | | A10G1944 | | | | |  |
| NAC+RECA_2+RECA_3 | | | | | | Gh_A01G0250 | | | | | 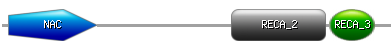 |
| NAC+Kinase | | | | | | Gh_D11G0701 | | | | |  |
| [*Ipomoea trifida*](http://planttfdb.cbi.pku.edu.cn/index.php?sp=Itr) | | | | | | | | | | | |
| Double domain | | | | | | Itr_sc004968.1_g00002.1 | | | | |  |
| NAC+Never growth factor | | | | | | Itr_sc000185.1_g00020.1 | | | | | 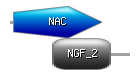 |
| NAC+integrase+RnaseH+Integrase | | | | | | Itr_sc004419.1_g00003.1 | | | | | 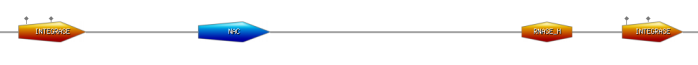 |
| *Jugulans regia* | | | | | | | | | | | |
| Double domain | | | | | | WALNUT_00018866-RA  WALNUT_00025170-RA  WALNUT_00031849-RA | | | | |  |
| *Leersia perrieri* | | | | | | | | | | | |
| Double domain | | | | | | LPERR04G14690.1  LPERR07G05500.1  LPERR07G05530.1  LPERR07G08560.1  LPERR07G12620.1 | | | | |  |
| NAC+CHCH | | | | | | LPERR04G10720.1 | | | | |  |
| NAC+PPR | | | | | | LPERR04G12800.1 | | | | |  |
| *Morus notabilis* | | | | | | | | | | | |
| NAC+PPR | | | | | | XP_010091280.1 | | | | | 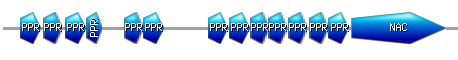 |
| NAC+CRM | | | | | | XP_010108843.1 | | | | |  |
| [*Nicotiana benthamiana*](http://planttfdb.cbi.pku.edu.cn/index.php?sp=Nbe) | | | | | | | | | | | |
| Double domain | | | | | | Niben101Scf04745g02009.1  Niben101Scf00193g01009.1 | | | | |  |
|  | | | | | | Niben101Scf07168g02012.1 | | | | | 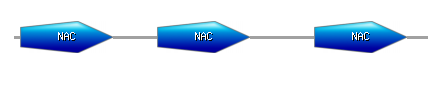 |
| NAC+PPR | | | | | | Niben101Scf09065g00002.1 | | | | |  |
| [*Ocimum tenuiflorum*](http://planttfdb.cbi.pku.edu.cn/index.php?sp=Ote) | | | | | | | | | | | |
| Double domain | | | | | | Ote100004440122  Ote100239950061 | | | | |  |
| NAC+Kinase | | | | | | Ote100092950061 | | | | |  |
| *Oropetium thomaeum* | | | | | | | | | | | |
| Double domain | | | | | | 20150105_15183A | | | | | 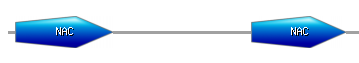 |
| *Oryza glumaepatula* | | | | | | | | | | | |
| Double domain | | | | | | OGLUM04G18300.1  OGLUM11G14240.1 | | | | |  |
| [*Oryza longistaminata*](http://planttfdb.cbi.pku.edu.cn/index.php?sp=Olo) | | | | | | | | | | | |
| Double domain | | | | | | KN539501.1_FGP004 | | | | |  |
| NAC+KH type | | | | | | KN538956.1_FGP004 | | | | | 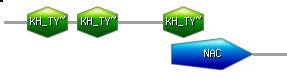 |
| NAC+PPR | | | | | | KN539106.1_FGP005 | | | | | 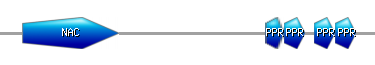 |
| NAC+LISH | | | | | | KN539151.1_FGP017 | | | | | 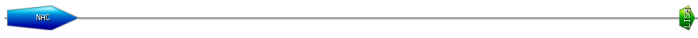 |
| NAC+RAB | | | | | | KN539582.1_FGP005 | | | | | 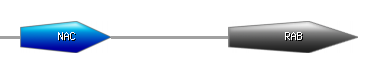 |
| NAC+DNAJ_2 | | | | | | KN540176.1_FGP005 | | | | |  |
| NAC+WRKY | | | | | | KN541034.1_FGP003 | | | | | 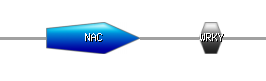 |
| *Oryza meridionalis* | | | | | | | | | | | |
| Double domain | | | | | | OMERI04G15260.1  OMERI12G00510.1 | | | | |  |
| NAC+CHCH | | | | | | OMERI04G12430.3 | | | | |  |
| NAC+RAB | | | | | | OMERI04G25580.2 | | | | |  |
| *Oryza nivara* | | | | | | | | | | | |
| Double domain | | | | | | ONIVA02G35680.1  ONIVA04G16580.1  ONIVA10G08840.1  ONIVA10G09230.1 | | | | |  |
| NAC+PPR | | | | | | ONIVA04G14230.1 | | | | |  |
| *Oryza punctata* | | | | | | | | | | | |
| Double domain | | | | | | OPUNC04G15950.1  OPUNC06G17430.1  OPUNC09G17350.1  OPUNC10G08540.1  OPUNC11G12330.1  OPUNC11G12330.1 | | | | |  |
| NAC+CHCH | | | | | | OPUNC04G11700.1 | | | | |  |
| *Oryza rufipogon* | | | | | | | | | | | |
| Double domain | | | | | | ORUFI02G12670.1  ORUFI04G19680.1  ORUFI09G20530.1  ORUFI10G09270.1 | | | | |  |
| Triple NAC domains | | | | | | ORUFI11G01500.1  ORUFI11G15730.1 | | | | | 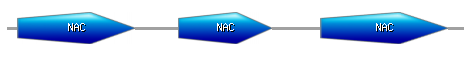 |
| NAC+PPR | | | | | | ORUFI04G17470.1 | | | | |  |
| *Oryza sativa* Indica | | | | | | | | | | | |
| Double domain | | | | | | BGIOSGA014751-PA | | | | |  |
| NAC+CHCH | | | | | | BGIOSGA015018-PA  BGIOSGA034998-PA | | | | |  |
| NAC+JACALIN_LECT | | | | | | BGIOSGA036980-PA | | | | |  |
| [*Panicum hallii*](http://planttfdb.cbi.pku.edu.cn/index.php?sp=Pha) | | | | | | | | | | | |
| Double domain | | | | | | | | | Pahal.A02025.1  Pahal.G00927.1  Pahal.G01013.1 | |  |
| NAC+ZF_BED | | | | | | | | | Pahal.C04447.1  Pahal.C04569.1  Pahal.J01717.1 | |  |
| NAC+ZF_BED+NAC+ZF_BED | | | | | | | | | Pahal.C04570.1  Pahal.C04572.1  Pahal.C04574.1 | | 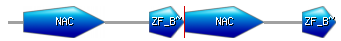 |
| *Panicum virgatum* | | | | | | | | | | | |
| Double domain | | | | | | | | | Pavir.1KG322500.1.p  Pavir.1KG445000.1.p  Pavir.2KG138100.1.p  Pavir.2KG140000.1.p  Pavir.4NG078000.1.p  Pavir.4NG078200.1.p  Pavir.5NG430200.1.p  Pavir.5NG465300.1.p  Pavir.7KG245100.1.p | |  |
| NAC+ZF_BED+ZF_BED | | | | | | | | | Pavir.3KG480900.1.p  Pavir.3NG254300.1.p | |  |
| NAC+ ZF_BED | | | | | | | | | Pavir.3KG499100.1.p  Pavir.3NG302500.1.p  Pavir.6NG320700.1.p  Pavir.6NG334600.1.p | |  |
| *Petunia axillaris* | | | | | | | | | | | |
| Double domain | | Peaxi162Scf00176g01638.1  Peaxi162Scf01007g00210.1  Peaxi162Scf00493g00439.1 | | | | | | | | |  |
| *Phoenix dactylifera* | | | | | | | | | | | |
| Double domain | | PDK_30s698051g001  PDK_30s758201g003  PDK_30s830451g001 | | | | | | | | |  |
| NAC+PPR | | PDK_30s1165641g010 | | | | | | | | |  |
| *Populus euphratica* | | | | | | | | | | | |
| Double domain | | | | | | | CCG003661.1  CCG010882.1 | | | |  |
| NAC+TIR | | | | | | | CCG001083.1 | | | |  |
| NAC+Zinc finger | | | | | | | CCG015305.2 | | | |  |
| NAC+RNA Recog Motif | | | | | | | CCG017923.1 | | | |  |
| *Prunus mume* | | | | | | | | | | | |
| Double NAC domain | | | | | | | XP_016648008.1 | | | |  |
| *Pseudotsuga menziesii* | | | | | | | | | | | |
| Double domain | | | | | | | PSME_00001409-RA  PSME_00011888-RA  PSME_00012319-RA  PSME_00017053-RA  PSME_00020635-RA | | | |  |
| NAC+JMJN | | | | | | | PSME_00016918-RA | | | | 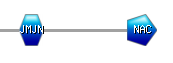 |
| NAC+APAG | | | | | | | PSME_00024440-RA | | | | 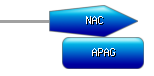 |
| NAC+HOMEO | | | | | | | PSME_00026525-RA | | | | 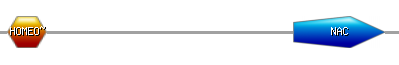 |
| [*Raphanus raphanistrum*](http://planttfdb.cbi.pku.edu.cn/index.php?sp=Rra) | | | | | | | | | | | |
| Double domain | | | | | | | RrC1847_p1  RrC9803_p2  RrC21868_p1  RrC5114_p2 | | | |  |
| NAC+RAB | | | | | | | RrC3821_p1 | | | |  |
| NAC+RRM+RRM | | | | | | | RrC20665_p1 | | | | 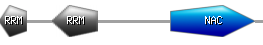 |
| NAC+ENT | | | | | | | RrC935_p1 | | | |  |
| *Raphanus sativus* | | | | | | | | | | | |
| Double domain | | | | | | | Rsa1.0_06656.1_g00001.1  Rsa1.0_00835.1_g00007.1  Rsa1.0_01306.1_g00007.1  Rsa1.0_01306.1_g00009.1  Rsa1.0_01979.1_g00001.1 | | | |  |
| NAC+ENT | | | | | | | Rsa1.0_00100.1_g00020.1 | | | |  |
| [*Salvia miltiorrhiza*](http://planttfdb.cbi.pku.edu.cn/index.php?sp=Smi) | | | | | | | | | | | |
| Double domain | | | | | | | SMil_00028195-RA_Salv | | | |  |
| NAC+Kinase | | | | | | | SMil_00004615-RA_Salv | | | |  |
| NAC+HOMEO | | | | | | | SMil_00008150-RA_Salv | | | |  |
| *Solanum melongena* | | | | | | | | | | | |
| Double domain | | | | | | | Sme2.5_00142.1_g00013.1 | | | |  |
| NAC+GH16+GH16+GH16 | | | | | | | Sme2.5_00096.1_g00018.1 | | | | 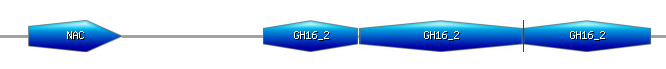 |
| NAC+INTEGRASE | | | | | | | Sme2.5_00756.1_g00007.1 | | | |  |
| NAC+IQ | | | | | | | Sme2.5_01725.1_g00009.1 | | | |  |
| *Solanum pennellii* | | | | | | | | | | | |
| NAC+ZF_C | | | | | | | Sopen02g012300.1  Sopen10g022760.1 | | | |  |
| [*Tarenaya hassleriana*](http://planttfdb.cbi.pku.edu.cn/index.php?sp=Tha) | | | | | | | | | | | |
| Double domain | | | | | | | XP_010536267.1 | | | |  |
| *Trifolium pratense* | | | | | | | | | | | |
| Double domain | | Tp57577_TGAC_v2_mRNA16525  Tp57577_TGAC_v2_mRNA31296 | | | | | | | | |  |
| NAC+Carrier+carrier | | Tp57577_TGAC_v2_mRNA14116 | | | | | | | | | 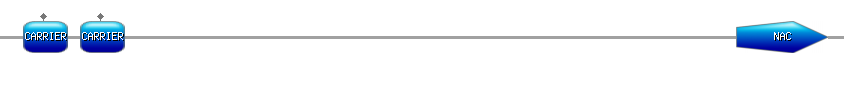 |
| NAC+PI3_4_Kinase3 | | Tp57577_TGAC_v2_mRNA38618 | | | | | | | | | 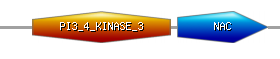 |
| *Vigna radiata* | | | | | | | | | | | |
| Double domain | | Vradi0051s00540.1  Vradi08g15940.1 | | | | | | | | |  |
| [*Zoysia japonica*](http://planttfdb.cbi.pku.edu.cn/index.php?sp=Zja) | | | | | | | | | | | |
| NAC+Response regulatory | | Zjn_sc00007.1.g11670.1.am.mk | | | | | | | | |  |
| NAC+CDPK | | sc00028.1.g03130.1.am.mk | | | | | | | | | 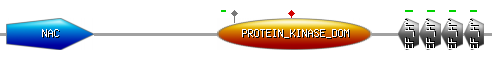 |
| NAC+BZIP | | Zjn_sc00067.1.g01060.1.am.mk | | | | | | | | | 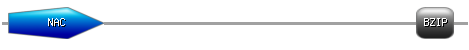 |
| NAC+ANK Repeat | | Zjn_sc00138.1.g00550.1.am.mk | | | | | | | | | 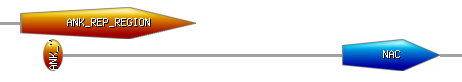 |
| [*Zoysia matrella*](http://planttfdb.cbi.pku.edu.cn/index.php?sp=Zmt) | | | | | | | | | | | |
| Double domain | | Zmw_sc00723.1.g00180.1 | | | | | | | | |  |
| NAC+ANK | | Zmw_sc04707.1.g00090.1 | | | | | | | | |  |
| NAC+ACT+ACT+ACT | | Zmw_sc00881.1.g00060.1 | | | | | | | | | 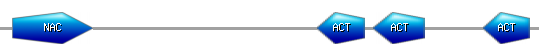 |
| NAC+BZIP | | Zmw_sc01551.1.g00060.1 | | | | | | | | |  |
| *Zoysia pacifica* | | | | | | | | | | | |
| Double domain | | Zpz_sc01162.1.g00050.1.am.mkhc | | | | | | | | |  |
| NAC+DCD | | Zpz_sc00130.1.g00590.1.am.mk | | | | | | | | | 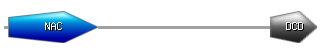 |
| NAC+ANK | | Zpz_sc04642.1.g00030.1.am.mk | | | | | | | | |  |
| *Glycine soja* | | | | | | | | | | | |
| NAC+LOGIN+V-SNARE | | | | | | | KHN21163.1 | | | | 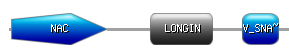 |
| *Sisymbrium irio* | | | | | | | | | | | |
| Double domain | | | | | | | 676713830  676785220 | | | |  |
| NAC+ENT | | | | | | | 676785308  676785330 | | | |  |
| *Thellungiella halophila* | | | | | | | | | | | |
| Double domain | | | | | | | Thhalv10015599m  Thhalv10024006m | | | |  |
