## Supplementary Table 2 for "Genomics, Molecular and Evolutionary Perspective of NAC Transcription Factors"

NAC TFs showing the presence of novel functional domain along with NAC domains.

| NAC TFs | Species | Active site |
|---|---|---|
| Pavir.Ha00781 | *Panicum virgatum* | *Aldehyde dehydrogenases cysteine active site* |
| Carubv10005139m | *Capsella rubella* | *Aldehyde dehydrogenases glutamic acid active site* |
| 491912 | *Arabidopsis lyrata* | *Aldehyde dehydrogenases glutamic acid active site* |
| GSBRNA2T00058100001 | *Brassica napus* | *Aldehyde dehydrogenases glutamic acid active site* |
| GSBRNA2T00078811001 | *Brassica napus* | *Aldehyde dehydrogenases glutamic acid active site* |
| XP_013638893.1 | *Brassica oleracea* | *Antenna complexes beta subunits signature* |
| Prupe.2G196600.1.p | *Prunus persica* | *ATP synthase alpha and beta subunits signature* |
| Pavir.J01198 | *Panicum virgatum* | *Cysteine proteases inhibitors signature* |
| GSMUA_Achr9P20400_001 | *Musa acuminata* | *Endopeptidase Clp serine active site* |
| ACHN289231 | *Actinidia chinensis* | *Heavy-metal-associated domain* |
| LOC_Os01g47670 | *Oryza sativa* | *Inorganic pyrophosphatase signature* |
| Thhalv10009374m | *Eutrema salsugineum* | *Inorganic pyrophosphatase signature* |
| Niben101Scf07168g02012.1 | *Nicotiana benthamiana* | *Lipocalin signature* |
| XP_011088938.1 | *Sesamum indicum* | *Pancreatic trypsin inhibitor (Kunitz) family signature* |
| KHN08560.1 | *Glycine soja* | *Putative AMP-binding domain signature* |
| MLOC_75795.1 | *Hordeum vulgare* | *Zinc carboxypeptidases, zinc-binding region 2 signature* |
| Vradi01g03390.1 | *Vigna radiata* | *2Fe-2S ferredoxin-type iron-sulfur binding region signature* |
| FANhyb_rscf00000665.1.g00005.1 | *Fragaria ananassa* | *2-oxo acid dehydrogenases acyltransferase component lipoyl binding site* |
| XP_010916182.1 | *Elaeis guineensi* | *4Fe-4S ferredoxin-type iron-sulfur binding domain profile* |
| XP_010916183.1 | *Elaeis guineensi* | *4Fe-4S ferredoxin-type iron-sulfur binding domain profile* |
| GSMUA_Achr11P03780_001 | *Musa acuminata* | *7,8-dihydro-6-hydroxymethylpterin-pyrophosphokinase signature* |
| GSMUA_Achr11P17510_001 | *Musa acuminata* | *7,8-dihydro-6-hydroxymethylpterin-pyrophosphokinase signature* |
| GSMUA_Achr6P36840_001 | *Musa acuminata* | *7,8-dihydro-6-hydroxymethylpterin-pyrophosphokinase signature* |
| GSMUA_Achr7P06640_001 | *Musa acuminata* | *7,8-dihydro-6-hydroxymethylpterin-pyrophosphokinase signature* |

| | | |
|---|---|---|
| GSMUA_Achr8P11590_001 | *Musa acuminata* | *7,8-dihydro-6-hydroxymethylpterin-pyrophosphokinase signature* |
| NNU_008894-RA | *Nelumbo nucifera* | *7,8-dihydro-6-hydroxymethylpterin-pyrophosphokinase signature* |
| NNU_012069-RA | *Nelumbo nucifera* | *7,8-dihydro-6-hydroxymethylpterin-pyrophosphokinase signature* |
| Phvul.011G005700 | *Phaseolus vulgaris* | *7,8-dihydro-6-hydroxymethylpterin-pyrophosphokinase signature* |
| PDK_30s1016001g009 | *Phoenix dactylifera* | *7,8-dihydro-6-hydroxymethylpterin-pyrophosphokinase signature* |
| Spipo1G0086600 | *Spirodela polyrhiza* | *7,8-dihydro-6-hydroxymethylpterin-pyrophosphokinase signature* |
| Zosma265g00020 | *Zostera marina* | *7,8-dihydro-6-hydroxymethylpterin-pyrophosphokinase signature* |
| ACHN002691 | *Actinidia chinensis* | *7,8-dihydro-6-hydroxymethylpterin-pyrophosphokinase signature* |
| ACHN093311 | *Actinidia chinensis* | *7,8-dihydro-6-hydroxymethylpterin-pyrophosphokinase signature* |
| ACHN157011 | *Actinidia chinensis* | *7,8-dihydro-6-hydroxymethylpterin-pyrophosphokinase signature* |
| ACHN317511 | *Actinidia chinensis* | *7,8-dihydro-6-hydroxymethylpterin-pyrophosphokinase signature* |
| ACO007609.1 | *Ananus comosus* | *7,8-dihydro-6-hydroxymethylpterin-pyrophosphokinase signature* |
| Aqcoe7G078200.1.p | *Aquilegiacoerulea* | *7,8-dihydro-6-hydroxymethylpterin-pyrophosphokinase signature* |
| Bv5_097990_fpfn.t1 | *Beta vulgaris* | *7,8-dihydro-6-hydroxymethylpterin-pyrophosphokinase signature* |
| Brast07G026300.1.p | *Brachypodium stacei* | *7,8-dihydro-6-hydroxymethylpterin-pyrophosphokinase signature* |
| XP_004514529.1 | *Cicer arietinum* | *7,8-dihydro-6-hydroxymethylpterin-pyrophosphokinase signature* |
| XP_010933718.1 | *Elaeis guineensi* | *7,8-dihydro-6-hydroxymethylpterin-pyrophosphokinase signature* |
| Glyma-09G231700.1.p | *Glycine max* | *7,8-dihydro-6-hydroxymethylpterin-pyrophosphokinase signature* |
| KHN11660.1 | *Glycine soja* | *7,8-dihydro-6-hydroxymethylpterin-pyrophosphokinase signature* |
| Medtr4g036030.1 | *Medicago truncatula* | *7,8-dihydro-6-hydroxymethylpterin-pyrophosphokinase signature* |
| XP_010108028.1 | *Morus notabilis* | *7,8-dihydro-6-hydroxymethylpterin-pyrophosphokinase signature* |
| PDK_30s1016001g009 | *Phoenix dactylifera* | *7,8-dihydro-6-hydroxymethylpterin-pyrophosphokinase signature* |
| Tp57577_TGAC_v2_mRNA3761 | *Trifolium pratense* | *7,8-dihydro-6-hydroxymethylpterin-pyrophosphokinase signature* |
| Vang05g07280 | *Vigna angularis* | *7,8-dihydro-6-hydroxymethylpterin-pyrophosphokinase signature* |
| C.cajan_20892 | *Cajanus cajan* | *7,8-dihydro-6-hydroxymethylpterin-pyrophosphokinase signature* |
| GSVIVT01001264001 | *Vitis vinifera* | *7,8-dihydro-6-hydroxymethylpterin-pyrophosphokinase signature* |
| Aradu.Y1DM8 | *Arachis duranensis* | *ABC transporters family signature* |
| Araip.HYM8C | *Arachis ipaensis* | *ABC transporters family signature* |

| | | |
|---|---|---|
| mrna03833-1-v1-0-hybrid | *Fragaria vesca* | *Adenosine and AMP deaminase signature* |
| PSME_00042313-RA | *Pseudotsuga menziesii* | *Adenosine and AMP deaminase signature* |
| Zosma69g00740 | *Zostera marina* | *Adipokinetic hormone family signature* |
| Pavir.8NG107600.1.p | *Panicum virgatum* | *Aldehyde dehydrogenases cysteine active site* |
| Pavir.8NG113600.1.p | *Panicum virgatum* | *Aldehyde dehydrogenases cysteine active site* |
| RrC2674_p3 | *Raphanus raphanistrum* | *Aldehyde dehydrogenases cysteine active site* |
| Rsa1.0_00285.1_g00018.1 | *Raphanus sativus* | *Aldehyde dehydrogenases cysteine active site* |
| Araha-3365s0007 | *Arabidopsis halleri* | *Aldehyde dehydrogenases glutamic acid active site* |
| AT4G28530 | *Arabidopsis thaliana* | *Aldehyde dehydrogenases glutamic acid active site* |
| Bra010362 | *Brassica rapa* | *Aldehyde dehydrogenases glutamic acid active site* |
| Bra011037 | *Brassica rapa* | *Aldehyde dehydrogenases glutamic acid active site* |
| Cagra-1225s0038 | *Capsella grandiflora* | *Aldehyde dehydrogenases glutamic acid active site* |
| evm.model.supercontig_80.94 | *Carica papaya* | *Aldehyde dehydrogenases glutamic acid active site* |
| Kalax.0013s0167 | *Kalanchoe laxiflora* | *Aldehyde dehydrogenases glutamic acid active site* |
| Kalax.0154s0005 | *Kalanchoe laxiflora* | *Aldehyde dehydrogenases glutamic acid active site* |
| Thhalv10025606m | *Thellungiella halophila* | *Aldehyde dehydrogenases glutamic acid active site* |
| Araha.3365s0007.1.p | *Arabidopsis halleri* | *Aldehyde dehydrogenases glutamic acid active site* |
| KFK29397.1 | *Arabis alpina* | *Aldehyde dehydrogenases glutamic acid active site* |
| XP_013603495.1 | *Brassica oleracea* | *Aldehyde dehydrogenases glutamic acid active site* |
| XP_009109087.1 | *Brassica rapa* | *Aldehyde dehydrogenases glutamic acid active site* |
| XP_009129548.1 | *Brassica rapa* | *Aldehyde dehydrogenases glutamic acid active site* |
| Csa10g014090.1 | *Camelina sativa* | *Aldehyde dehydrogenases glutamic acid active site* |
| Csa11g015230.1 | *Camelina sativa* | *Aldehyde dehydrogenases glutamic acid active site* |
| Csa12g021980.1 | *Camelina sativa* | *Aldehyde dehydrogenases glutamic acid active site* |
| Carubv10005139m | *Capsella rubella* | *Aldehyde dehydrogenases glutamic acid active site* |
| Thhalv10025606m | *Eutrema salsugineum* | *Aldehyde dehydrogenases glutamic acid active site* |
| Kalax-0013s0167.1.p | *Kalanchoe marnieriana* | *Aldehyde dehydrogenases glutamic acid active site* |
| Kalax-0154s0005.1.p | *Kalanchoe marnieriana* | *Aldehyde dehydrogenases glutamic acid active site* |

| | | |
|---|---|---|
| RrC20539_p1 | *Raphanus raphanistrum* | *Aldehyde dehydrogenases glutamic acid active site* |
| Rsa1.0_13977.1_g00001.1 | *Raphanus sativus* | *Aldehyde dehydrogenases glutamic acid active site* |
| GSBRNA2T00104607001 | *Brassica napus* | *Aldehyde dehydrogenases glutamic acid active site* |
| Bv3_050780_usim.t1 | *Beta vulgaris* | *Aldo/keto reductase family putative active site signature* |
| FANhyb_rscf00000207.1.g00009.1 | *Fragaria ananassa* | *Aldo/keto reductase family putative active site signature* |
| Zmw_sc01138.1.g00130.1 | *Zoysia matrella* | *Aldo/keto reductase family putative active site signature* |
| Zpz_sc00095.1.g00090.1.am.mk | *Zoysia pacifica* | *Aldo/keto reductase family putative active site signature* |
| LOC_Os12g22940 | *Oryza sativa* | *Alkaline phosphatase active site* |
| OMERI12G11070.1 | *Oryza meridionalis* | *Alkaline phosphatase active site* |
| ORUFI12G10910.1 | *Oryza rufipogon* | *Alkaline phosphatase active site* |
| BGIOSGA037293.1 | *Oryza sativa Indica* | *Alkaline phosphatase active site* |
| LOC_Os12g22940.1 | *Oryza sativa japonica* | *Alkaline phosphatase active site* |
| Carubv10012206m | *Capsella rubella* | *Aminoacyl-transfer RNA synthetases class-I signature* |
| Oropetium_20150105_22096 | *Oropetium thomaeum* | *Aminotransferases class-II pyridoxal-phosphate attachment site* |
| ObartAA03S_FGP10288 | *Oryza barthii* | *Aminotransferases class-II pyridoxal-phosphate attachment site* |
| ORGLA12G0169600.1 | *Oryza glaberrima* | *Aminotransferases class-II pyridoxal-phosphate attachment site* |
| OpuncBB_FGP10000 | *Oryza punctata* | *Aminotransferases class-II pyridoxal-phosphate attachment site* |
| LOC_Os12g43530 | *Oryza sativa* | *Aminotransferases class-II pyridoxal-phosphate attachment site* |
| Oropetium_20150105_22096A | *Oropetium thomaeum* | *Aminotransferases class-II pyridoxal-phosphate attachment site* |
| ORGLA12G0169600.1 | *Oryza glaberrima* | *Aminotransferases class-II pyridoxal-phosphate attachment site* |
| OGLUM12G21360.1 | *Oryza glumaepatula* | *Aminotransferases class-II pyridoxal-phosphate attachment site* |
| KN539001.1 | *Oryza longistaminata* | *Aminotransferases class-II pyridoxal-phosphate attachment site* |
| OMERI12G14510 | *Oryza meridionalis* | *Aminotransferases class-II pyridoxal-phosphate attachment site* |
| ONIVA12G18630.1 | *Oryza nivara* | *Aminotransferases class-II pyridoxal-phosphate attachment site* |
| OPUNC12G17870.1 | *Oryza punctata* | *Aminotransferases class-II pyridoxal-phosphate attachment site* |
| ORUFI12G21970.1 | *Oryza rufipogon* | *Aminotransferases class-II pyridoxal-phosphate attachment site* |
| BGIOSGA035784.1 | *Oryza sativa Indica* | *Aminotransferases class-II pyridoxal-phosphate attachment site* |
| LOC_Os12g43530.1 | *Oryza sativa japonica* | *Aminotransferases class-II pyridoxal-phosphate attachment site* |

| | | |
|---|---|---|
| XP_009146239.1 | *Brassica rapa* | *Antenna complexes beta subunits signature* |
| Zmw_sc05821.1.g00010.1 | *Zoysia matrella* | *ArgE / dapE / ACY1 / CPG2 / yscS family signature 1* |
| Zpz_sc00055.1.g00240.1.am.mkhc | *Zoysia pacifica* | *ArgE / dapE / ACY1 / CPG2 / yscS family signature 1* |
| Medtr4g134460.1 | *Medicago truncatula* | *Aspartate and glutamate racemases signature 1* |
| XP_010545324.1 | *Tarenaya hassleriana* | *Aspartate and glutamate racemases signature 1* |
| XP_010545325.1 | *Tarenaya hassleriana* | *Aspartate and glutamate racemases signature 1* |
| Brast02G382000.1.p | *Brachypodium stacei* | *Aspartokinase signature* |
| Brast02G383700.1.p | *Brachypodium stacei* | *Aspartokinase signature* |
| EMT33859 | *Aegilops tauschii* | ATP Binding site and proton acceptor |
| Gorai.008G130300.1 | *Gossypium raimondii* | *ATP synthase alpha and beta subunits signature* |
| PH01000820G0540 | *Phyllostachys heterocycla* | *ATP synthase alpha and beta subunits signature* |
| Pta007882 | *Pinus taeda* | *ATP synthase alpha and beta subunits signature* |
| ppa020746m | *Prunus persica* | *ATP synthase alpha and beta subunits signature* |
| Solyc08g008660.2.1 | *Solanum lycopersicum* | *ATP synthase alpha and beta subunits signature* |
| PGSC0003DMP400010296 | *Solanum tuberosum* | *ATP synthase alpha and beta subunits signature* |
| PGSC0003DMP400010297 | *Solanum tuberosum* | *ATP synthase alpha and beta subunits signature* |
| CA11g08290 | *Capsicum annuum* | *ATP synthase alpha and beta subunits signature* |
| Cotton_A_26426_BGI-A2_v1.0 | *Gossypium arboreum* | *ATP synthase alpha and beta subunits signature* |
| XP_010100656.1 | *Morus notabilis* | *ATP synthase alpha and beta subunits signature* |
| PH01000820G0540 | *Phyllostachys heterocycla* | *ATP synthase alpha and beta subunits signature* |
| XP_008233055.1 | *Prunus mume* | *ATP synthase alpha and beta subunits signature* |
| Sme2.5_02517.1_g00007.1 | *Solanum melongena* | *ATP synthase alpha and beta subunits signature* |
| Sopen08g004490.1 | *Solanum pennellii* | *ATP synthase alpha and beta subunits signature* |
| Zmw_sc04326.1.g00020.1 | *Zoysia matrella* | *ATP synthase alpha and beta subunits signature* |
| Ciclev10019282m | *Citrus clementina* | *ATP synthase alpha and beta subunits signature* |
| Gh_A12G1049 | *Gossypium hirsutum* | *ATP synthase alpha and beta subunits signature* |
| Sopim08g008660.0.1 | *Solanum pimpinellifolium* | *ATP synthase alpha and beta subunits signature* |
| Pbr032225.1 | *Pyrus bretschneideri* | *ATP-dependent DNA ligase AMP-binding site* |

| | | |
|---|---|---|
| Pbr032225.1 | *Pyrus bretschneideri* | *ATP-dependent DNA ligase AMP-binding site* |
| Spipo1G0084400 | *Spirodela polyrhiza* | *Bacterial regulatory proteins, araC family signature* |
| MLOC_19933.1 | *Hordeum vulgare* | *Beta-ketoacyl synthases active site* |
| MLOC_19933.1 | *Hordeum vulgare* | *Beta-ketoacyl synthases active site* |
| Thecc1EG018508t | *Theobroma cacao* | *C-5 cytosine-specific DNA methylases active site* |
| Aqcoe1G058100.1.p | *Aquilegiacoerulea* | *Cadherin domain signature* |
| KFK30695.1 | *Arabis alpina* | *Carbamoyl-phosphate synthase subdomain signature 2* |
| ObartAA03S_FGP25558 | *Oryza barthii* | *Cysteine proteases inhibitors signature* |
| ORGLA06G0236900.1 | *Oryza glaberrima* | *Cysteine proteases inhibitors signature* |
| OpuncBB_FGP25111 | *Oryza punctata* | *Cysteine proteases inhibitors signature* |
| LOC_Os06g51070 | *Oryza sativa* | *Cysteine proteases inhibitors signature* |
| 462845378 | *Eragrostis tef* | *Cysteine proteases inhibitors signature* |
| ORGLA06G0236900.1 | *Oryza glaberrima* | *Cysteine proteases inhibitors signature* |
| OGLUM06G29360.1 | *Oryza glumaepatula* | *Cysteine proteases inhibitors signature* |
| OMERI06G27920.1 | *Oryza meridionalis* | *Cysteine proteases inhibitors signature* |
| ONIVA06G30950.1 | *Oryza nivara* | *Cysteine proteases inhibitors signature* |
| OPUNC06G25510.1 | *Oryza punctata* | *Cysteine proteases inhibitors signature* |
| ORUFI06G29930.1 | *Oryza rufipogon* | *Cysteine proteases inhibitors signature* |
| BGIOSGA020507.1 | *Oryza sativa Indica* | *Cysteine proteases inhibitors signature* |
| LOC_Os06g51070.1 | *Oryza sativa japonica* | *Cysteine proteases inhibitors signature* |
| Pavir.4NG342400.1.p | *Panicum virgatum* | *Cysteine proteases inhibitors signature* |
| Pavir.7KG085500.1.p | *Panicum virgatum* | *Cysteine proteases inhibitors signature* |
| Zmw_sc02823.1.g00180.1 | *Zoysia matrella* | *Cysteine proteases inhibitors signature* |
| Zpz_sc01253.1.g00030.1.sm.mk | *Zoysia pacifica* | *Cysteine proteases inhibitors signature* |
| Zjn_sc00068.1.g03510.1.sm.mk | *Zoysia japonica* | *Cysteine proteases inhibitors signature* |
| OGLUM01G06950.1 | *Oryza glumaepatula* | *Cytochrome P450 cysteine heme-iron ligand signature* |
| KN539290.1 | *Oryza longistaminata* | *Cytochrome P450 cysteine heme-iron ligand signature* |
| KN539290.1 | *Oryza longistaminata* | *Cytochrome P450 cysteine heme-iron ligand signature* |

| | | |
|---|---|---|
| OMERI07G10400 | *Oryza meridionalis* | *Cytochrome P450 cysteine heme-iron ligand signature* |
| Zmw_sc00276.1.g00230.1 | *Zoysia matrella* | *Cytochrome P450 cysteine heme-iron ligand signature* |
| Zmw_sc02343.1.g00060.1 | *Zoysia matrella* | *Cytochrome P450 cysteine heme-iron ligand signature* |
| Zpz_sc03671.1.g00030.1.am.mk | *Zoysia pacifica* | *Cytochrome P450 cysteine heme-iron ligand signature* |
| Sobic.003G105800.1.p | *Sorghum bicolor* | *Endopeptidase Clp serine active site* |
| GRMZM2G123246_P01 | *Vzea mays* | *Endopeptidase Clp serine active site* |
| EMT14807 | *Aegilops tauschii* | *Eukaryotic and viral aspartyl proteases active site* |
| Tae025010 | *Triticum aestivum* | *Eukaryotic and viral aspartyl proteases active site* |
| Tae030617 | *Triticum aestivum* | *Eukaryotic and viral aspartyl proteases active site* |
| Tae045052 | *Triticum aestivum* | *Eukaryotic and viral aspartyl proteases active site* |
| EMS63374 | *Triticum urartu* | *Eukaryotic and viral aspartyl proteases active site* |
| Itr_sc004419.1_g00003.1 | *Ipomea trifida* | *Eukaryotic and viral aspartyl proteases active site* |
| Traes_7DL_BDD45DB24.2 | *Triticum aestivum* | *Eukaryotic and viral aspartyl proteases active site* |
| Zjn_sc00092.1.g00060.1.am.mk | *Zoysia japonica* | *Eukaryotic and viral aspartyl proteases active site* |
| Zmw_sc03455.1.g00010.1 | *Zoysia matrella* | *Eukaryotic and viral aspartyl proteases active site* |
| Zpz_sc03692.1.g00060.1.sm.mk | *Zoysia pacifica* | *Eukaryotic and viral aspartyl proteases active site* |
| Niben101Scf07508g02004.1 | *Nicotiana benthamiana* | *FGGY family of carbohydrate kinases signature 2* |
| MA_18939g0010 | *Picea abies* | *Fumarate lyases signature* |
| Rsa1.0_01603.1_g00007.1 | *Raphanus sativus* | *Fumarate lyases signature* |
| GSMUA_Achr10P10790_001 | *Musa acuminata* | *GHMP kinases putative ATP-binding domain* |
| Peinf101Scf00337g03028.1 | *Petunia inflata* | *Glucoamylase active site region signature* |
| Cucsa.083540.1 | *Cucumis sativus* | *Glyceraldehyde 3-phosphate dehydrogenase active* |
| ONIVA05G06750.1 | *Oryza nivara* | *Glycoprotease family signature* |
| Cc08_g16900 | *Coffea canephora* | *Glycosyl hydrolases family 5 signature* |
| 30147.m014211 | *Ricinus communis* | *Glycosyl hydrolases family 9 active sites signature 2* |
| Kalax-0686s0005.1.p | *Kalanchoe marnieriana* | *Hemopexin domain signature* |
| Kalax-0609s0006.1.p | *Kalanchoe marnieriana* | *Hemopexin domain signature* |
| Scf00060.g6196.t1 | *Utricularia gibba* | *Histone H4 signature* |

| | | |
|---|---|---|
| XP_016648489.1 | *Prunus mume* | *Histone H4 signature* |
| Prupe.4G138500.1.p | *Prunus persica* | *Histone H4 signature* |
| Zmw_sc06792.1.g00040.1 | *Zoysia matrella* | *Histone H4 signature* |
| ObartAA03S_FGP17273 | *Oryza barthii* | *HMG-I and HMG-Y DNA-binding domain (A+T-hook)* |
| LOC_Os03g59730 | *Oryza sativa* | *HMG-I and HMG-Y DNA-binding domain (A+T-hook)* |
| Bradi1g04229.1.p | *Brachypodium distachyon* | *HMG-I and HMG-Y DNA-binding domain (A+T-hook)* |
| Brast02G355100.1.p | *Brachypodium stacei* | *HMG-I and HMG-Y DNA-binding domain (A+T-hook)* |
| OGLUM03G38320.1 | *Oryza glumaepatula* | *HMG-I and HMG-Y DNA-binding domain (A+T-hook)* |
| OMERI03G35410.1 | *Oryza meridionalis* | *HMG-I and HMG-Y DNA-binding domain (A+T-hook)* |
| ORUFI03G40160.1 | *Oryza rufipogon* | *HMG-I and HMG-Y DNA-binding domain (A+T-hook)* |
| BGIOSGA009581.1 | *Oryza sativa Indica* | *HMG-I and HMG-Y DNA-binding domain (A+T-hook)* |
| LOC_Os03g59730.1 | *Oryza sativa japonica* | *HMG-I and HMG-Y DNA-binding domain (A+T-hook)* |
| Pbr020079.1 | *Pyrus bretschneideri* | *Immunoglobulins and major histocompatibility complex proteins signature* |
| EcC046909.20 | *Eucalyptus camaldulensis* | *Immunoglobulins and major histocompatibility complex proteins signature* |
| EcS504758.10 | *Eucalyptus camaldulensis* | *Immunoglobulins and major histocompatibility complex proteins signature* |
| Pbr020079.1 | *Pyrus bretschneideri* | *Immunoglobulins and major histocompatibility complex proteins signature* |
| ORGLA01G0211000.1 | *Oryza glaberrima* | *Inorganic pyrophosphatase signature* |
| Pbr026815.1 | *Pyrus bretschneideri* | *Inorganic pyrophosphatase signature* |
| Thhalv10009374m | *Thellungiella halophila* | *Inorganic pyrophosphatase signature* |
| XP_010108843.1 | *Morus notabilis* | *Inorganic pyrophosphatase signature* |
| ORGLA01G0211000.1 | *Oryza glaberrima* | *Inorganic pyrophosphatase signature* |
| OGLUM01G29620.1 | *Oryza glumaepatula* | *Inorganic pyrophosphatase signature* |
| OMERI01G23440.1 | *Oryza meridionalis* | *Inorganic pyrophosphatase signature* |
| ONIVA01G29020.1 | *Oryza nivara* | *Inorganic pyrophosphatase signature* |
| ORUFI01G28690.1 | *Oryza rufipogon* | *Inorganic pyrophosphatase signature* |

| | | |
|---|---|---|
| LOC_Os01g47670.1 | *Oryza sativa japonica* | *Inorganic pyrophosphatase signature* |
| Pbr026815.1 | *Pyrus bretschneideri* | *Inorganic pyrophosphatase signature* |
| GSVIVT01026495001 | *Vitis vinifera* | *Iron-containing alcohol dehydrogenases signature 1* |
| OB07G24770.1 | *Oryza brachyantha* | *Legume lectins beta-chain signature* |
| ORGLA03G0364100.1 | *Oryza glaberrima* | *Legume lectins beta-chain signature* |
| OGLUM03G39450.1 | *Oryza glumaepatula* | *Legume lectins beta-chain signature* |
| OGLUM03G39510.1 | *Oryza glumaepatula* | *Legume lectins beta-chain signature* |
| KN538833.1 | *Oryza longistaminata* | *Legume lectins beta-chain signature* |
| OMERI03G36390.1 | *Oryza meridionalis* | *Legume lectins beta-chain signature* |
| OMERI03G36430.1 | *Oryza meridionalis* | *Legume lectins beta-chain signature* |
| ONIVA03G41430.1 | *Oryza nivara* | *Legume lectins beta-chain signature* |
| ONIVA03G41480.1 | *Oryza nivara* | *Legume lectins beta-chain signature* |
| OPUNC03G36710.1 | *Oryza punctata* | *Legume lectins beta-chain signature* |
| OPUNC03G36760.1 | *Oryza punctata* | *Legume lectins beta-chain signature* |
| BGIOSGA013858.1 | *Oryza sativa Indica* | *Legume lectins beta-chain signature* |
| BGIOSGA013866.1 | *Oryza sativa Indica* | *Legume lectins beta-chain signature* |
| LOC_Os03g61249.1 | *Oryza sativa japonica* | *Legume lectins beta-chain signature* |
| LOC_Os03g61319.1 | *Oryza sativa japonica* | *Legume lectins beta-chain signature* |
| Nta011914 | *Nicotiana tabacum* | *Lipocalin signature* |
| Niben101Ctg05879g00002.1 | *Nicotiana benthamiana* | *Lipocalin signature* |
| Niben101Scf01337g00001.1 | *Nicotiana benthamiana* | *Lipocalin signature* |
| Niben101Scf08665g00007.1 | *Nicotiana benthamiana* | *Lipocalin signature* |
| Niben101Scf13438g00001.1 | *Nicotiana benthamiana* | *Lipocalin signature* |
| Niben101Scf15279g00002.1 | *Nicotiana benthamiana* | *Lipocalin signature* |
| XP_009762991.1 | *Nicotiana sylvestris* | *Lipocalin signature* |
| XP_009763090.1 | *Nicotiana sylvestris* | *Lipocalin signature* |
| XP_009782335.1 | *Nicotiana sylvestris* | *Lipocalin signature* |
| XP_009792222.1 | *Nicotiana sylvestris* | *Lipocalin signature* |

| | | |
|---|---|---|
| XP_009794230.1 | *Nicotiana sylvestris* | *Lipocalin signature* |
| XP_016449500.1 | *Nicotiana tabacum* | *Lipocalin signature* |
| XP_016469037.1 | *Nicotiana tabacum* | *Lipocalin signature* |
| XP_016474545.1 | *Nicotiana tabacum* | *Lipocalin signature* |
| XP_016477162.1 | *Nicotiana tabacum* | *Lipocalin signature* |
| XP_016490755.1 | *Nicotiana tabacum* | *Lipocalin signature* |
| XP_016504577.1 | *Nicotiana tabacum* | *Lipocalin signature* |
| XP_016510726.1 | *Nicotiana tabacum* | *Lipocalin signature* |
| XP_009598748.1 | *Nicotiana tomentosiformis* | *Lipocalin signature* |
| XP_009601217.1 | *Nicotiana tomentosiformis* | *Lipocalin signature* |
| XP_009602879.1 | *Nicotiana tomentosiformis* | *Lipocalin signature* |
| XP_009628837.1 | *Nicotiana tomentosiformis* | *Lipocalin signature* |
| KFK42489.1 | *Arabis alpina* | *Mannitol dehydrogenases signature* |
| OB09G21580.1 | *Oryza brachyanta* | *N-6 Adenine-specific DNA methylases signature* |
| Sobic.003G110900.1.p | *Sorghum bicolor* | *N-6 Adenine-specific DNA methylases signature* |
| Zosma53g00110 | *Zostera marina* | *N-6 Adenine-specific DNA methylases signature* |
| Aqcoe2G375500.1.p | *Aquilegiacoerulea* | *N-6 Adenine-specific DNA methylases signature* |
| scaffold07459-snap-gene-0.9 | *Castanea mollissima* | *N-6 Adenine-specific DNA methylases signature* |
| OB04G27030.1 | *Oryza brachyantha* | *N-6 Adenine-specific DNA methylases signature* |
| OB09G21580.1 | *Oryza brachyantha* | *N-6 Adenine-specific DNA methylases signature* |
| Csa16g023250.1 | *Camelina sativa* | *Neutral zinc metallopeptidases, zinc-binding region signature* |
| Ote100091280041 | *Ocimum tenuiflorum* | *Neutral zinc metallopeptidases, zinc-binding region signature* |
| Neem_8133_f_4 | *Azadirachta indica* | *Nt-dnaJ domain signature* |
| FANhyb_rscf00000018.1.g00013.1 | *Fragaria ananassa* | *Nt-dnaJ domain signature* |
| EPS67213.1 | *Genlisea aurea* | *Peroxidases active site signature* |
| Ciclev10008619m | *Citrus clementina* | *pfkB family of carbohydrate kinases signature 1* |
| Pavir.J21162 | *Panicum virgatum* | *pfkB family of carbohydrate kinases signature 2* |
| Thhalv10006408m | *Thellungiella halophila* | *pfkB family of carbohydrate kinases signature 2* |

| | | |
|---|---|---|
| Thhalv10006408m | *Eutrema salsugineum* | *pfkB family of carbohydrate kinases signature 2* |
| Zpz_sc00266.1.g00320.1.am.mk | *Zoysia pacifica* | *Phospholipase A2 histidine active site* |
| GSMUA_Achr11P08970_001 | *Musa acuminata* | *Phosphopantetheine attachment site* |
| ObartAA03S_FGP27339 | *Oryza barthii* | *Phosphopantetheine attachment site* |
| ORGLA07G0243600.1 | *Oryza glaberrima* | *Phosphopantetheine attachment site* |
| OpuncBB_FGP26901 | *Oryza punctata* | *Phosphopantetheine attachment site* |
| LOC_Os07g37920 | *Oryza sativa* | *Phosphopantetheine attachment site* |
| Sobic.002G342100.1.p | *Sorghum bicolor* | *Phosphopantetheine attachment site* |
| Sobic.002G342100.1.p | *Sorghum bicolor* | *Phosphopantetheine attachment site* |
| GRMZM2G179885_P01 | *Zea mays* | *Phosphopantetheine attachment site* |
| scaffold07067-augustus-gene-0.6 | *Castanea mollissima* | *Phosphopantetheine attachment site* |
| Glyma-13G174700.1.p | *Glycine max* | *Phosphopantetheine attachment site* |
| KHN09961.1 | *Glycine soja* | *Phosphopantetheine attachment site* |
| LPERR07G16320.1 | *Leersia perrieri* | *Phosphopantetheine attachment site* |
| ORGLA07G0243600.1 | *Oryza glaberrima* | *Phosphopantetheine attachment site* |
| OGLUM07G18780.1 | *Oryza glumaepatula* | *Phosphopantetheine attachment site* |
| KN538693.1 | *Oryza longistaminata* | *Phosphopantetheine attachment site* |
| OMERI07G15720.1 | *Oryza meridionalis* | *Phosphopantetheine attachment site* |
| ONIVA07G17340.1 | *Oryza nivara* | *Phosphopantetheine attachment site* |
| OPUNC07G17890.1 | *Oryza punctata* | *Phosphopantetheine attachment site* |
| ORUFI07G19770.1 | *Oryza rufipogon* | *Phosphopantetheine attachment site* |
| BGIOSGA024071.1 | *Oryza sativa Indica* | *Phosphopantetheine attachment site* |
| LOC_Os07g37920.1 | *Oryza sativa japonica* | *Phosphopantetheine attachment site* |
| KFK41460.1 | *Arabis alpina* | *Polygalacturonase active site* |
| Rsa1.0_00061.1_g00015.1 | *Raphanus sativus* | *Polygalacturonase active site* |
| Sobic.005G041000.1.p | *Sorghum bicolor* | *Polyprenyl synthases signature 1* |
| .XP_010536267.1 | *Tarenaya hassleriana* | *PPM-type phosphatase domain signature* |
| Solyc04g079940.2.1 | *Solanum lycopersicum* | *Prokaryotic membrane lipoprotein lipid attachment site profile* |

| | | |
|---|---|---|
| Scf00187.g11893.t1 | *Utricularia gibba* | *Prokaryotic membrane lipoprotein lipid attachment site profile* |
| Niben101Scf01153g00009.1 | *Nicotiana benthamiana* | *Prokaryotic membrane lipoprotein lipid attachment site profile* |
| Niben101Scf01153g00009.1 | *Nicotiana benthamiana* | *Prokaryotic membrane lipoprotein lipid attachment site profile* |
| XP_009789698.1 | *Nicotiana sylvestris* | *Prokaryotic membrane lipoprotein lipid attachment site profile* |
| XP_009789699.1 | *Nicotiana sylvestris* | *Prokaryotic membrane lipoprotein lipid attachment site profile* |
| XP_016453035.1 | *Nicotiana tabacum* | *Prokaryotic membrane lipoprotein lipid attachment site profile* |
| XP_016471015.1 | *Nicotiana tabacum* | *Prokaryotic membrane lipoprotein lipid attachment site profile* |
| XP_016487616.1 | *Nicotiana tabacum* | *Prokaryotic membrane lipoprotein lipid attachment site profile* |
| XP_016487617.1 | *Nicotiana tabacum* | *Prokaryotic membrane lipoprotein lipid attachment site profile* |
| XP_009603640.1 | *Nicotiana tomentosiformis* | *Prokaryotic membrane lipoprotein lipid attachment site profile* |
| Sopim04g079940.0.1 | *Solanum pimpinellifolium* | *Prokaryotic membrane lipoprotein lipid attachment site profile* |
| Glyma-13G243200.1.p | *Glycine max* | *Putative AMP-binding domain signature* |
| Si039061m | *Setaria italica* | *Regulator of chromosome condensation (RCC1) signature 2* |
| Seita.9G169600.1.p | *Setaria italica* | *Regulator of chromosome condensation (RCC1) signature 2* |
| GRMZM2G156977_P01 | *Zea mays* | *Ribosomal protein L24e signature* |
| Cotton_A_00183_BGI-A2_v1.0 | *Gossypium arboreum* | *Ribosomal protein L31 signature* |
| Gh_A01G0250 | *Gossypium hirsutum* | *Ribosomal protein L31 signature* |
| KN539025.1 | *Oryza longistaminata* | *Ribosomal protein L36 signature* |
| RrC831_p3 | *Raphanus raphanistrum* | *Ribosomal protein S10 signature* |
| Cotton_A_09649_BGI-A2_v1.0 | *Gossypium arboreum* | *Ribosomal protein S19e signature* |
| Zjn_sc00034.1.g02530.1.am.mk | *Zoysia japonica* | *Ribosomal protein S8 signature* |
| Spipo0G0093000 | *Spirodela polyrhiza* | *Ribosome-binding factor A signature* |
| PH01000491G0390 | *Phyllostachys heterocycla* | *Rubredoxin signature* |
| PH01000491G0390 | *Phyllostachys heterocycla* | *Rubredoxin signature* |
| Bradi3g12470.1 | *Brachypodium distachyon* | *Serine proteases, subtilase family, aspartic acid active site* |
| Pahal.F00337 | *Panicum hallii* | *Serine proteases, subtilase family, aspartic acid active site* |
| Pavir.Fa00048 | *Panicum virgatum* | *Serine proteases, subtilase family, aspartic acid active site* |
| Si013361m | *Setaria italica* | *Serine proteases, subtilase family, aspartic acid active site* |

| | | |
|---|---|---|
| Si013378m | *Setaria italica* | *Serine proteases, subtilase family, aspartic acid active site* |
| Si013524m | *Setaria italica* | *Serine proteases, subtilase family, aspartic acid active site* |
| Sevir.6G256500 | *Setaria viridis* | *Serine proteases, subtilase family, aspartic acid active site* |
| GRMZM2G125777_P01 | *Zea mays* | *Serine proteases, subtilase family, aspartic acid active site* |
| Bradi3g12470.1.p | *Brachypodium distachyon* | *Serine proteases, subtilase family, aspartic acid active site* |
| Brast03G309200.1.p | *Brachypodium stacei* | *Serine proteases, subtilase family, aspartic acid active site* |
| Do003945.1 | *Dichanthelium oligosanthes* | *Serine proteases, subtilase family, aspartic acid active site* |
| Itr_sc000194.1_g00003.1 | *Ipomea trifida* | *Serine proteases, subtilase family, aspartic acid active site* |
| Pahal.F00337.1 | *Panicum hallii* | *Serine proteases, subtilase family, aspartic acid active site* |
| Pavir.6KG413300.1.p | *Panicum virgatum* | *Serine proteases, subtilase family, aspartic acid active site* |
| Pavir.6NG362800.1.p | *Panicum virgatum* | *Serine proteases, subtilase family, aspartic acid active site* |
| Seita.6G251900.1.p | *Setaria italica* | *Serine proteases, subtilase family, aspartic acid active site* |
| Sme2.5_05012.1_g00010.1 | *Solanum melongena* | *Serine proteases, subtilase family, aspartic acid active site* |
| XP_010543143.1 | *Tarenaya hassleriana* | *Serine proteases, subtilase family, aspartic acid active site* |
| XP_010543151.1 | *Tarenaya hassleriana* | *Serine proteases, subtilase family, aspartic acid active site* |
| XP_010543159.1 | *Tarenaya hassleriana* | *Serine proteases, subtilase family, aspartic acid active site* |
| Traes_5BL_3BA7EA9A3 | *Triticum aestivum* | *Serine proteases, subtilase family, aspartic acid active site* |
| cassava4-1_025821m | *Manihot esculenta* | *Serine proteases, trypsin family, serine active site* |
| KZV46422.1 | *Dorcoceras hygrometricum* | *Serine/Threonine protein kinases active-site signature* |
| Gh_D11G0701 | *Gossypium hirsutum* | *Serine/Threonine protein kinases active-site signature* |
| XP_010551540.1 | *Tarenaya hassleriana* | *Serine/threonine specific protein phosphatases signature* |
| OMERI04G25580.2 | *Oryza meridionalis* | *Sigma-54 interaction domain ATP-binding region A signature* |
| RrC3821_p1 | *Raphanus raphanistrum* | *Sigma-54 interaction domain ATP-binding region A signature* |
| AUR62000898 | *Chenopodium quinoa* | *Signal peptidases I serine active site* |
| Cucsa.284890.1 | *Cucumis sativus* | *Signal peptidases I serine active site* |
| scaffold02306-augustus-gene-0.21 | *Castanea mollissima* | *Signal peptidases I serine active site* |
| OB07G32110.1 | *Oryza brachyanta* | *Signal peptidases I signature 3* |
| GSBRNA2T00106219001 | *Brassica napus* | *Signal peptidases I signature 3* |

| | | |
|---|---|---|
| OB07G32110.1 | *Oryza brachyantha* | *Signal peptidases I signature 3* |
| XP_013583708.1 | *Brassica oleracea* | *Signal peptidases I signature 3* |
| Tp57577_TGAC_v2_mRNA33521 | *Trifolium pratense* | *Soybean trypsin inhibitor (Kunitz) protease inhibitors family signature* |
| Aqcoe3G142100.1.p | *Aquilegiacoerulea* | *SRP54-type proteins GTP-binding domain signature* |
| EcC047434.20 | *Eucalyptus camaldulensis* | *Sugar transport proteins signature 2* |
| Eucgr-I01940.1.p | *Eucalyptus grandis* | *Sugar transport proteins signature 2* |
| KHN21163.1 | *Glycine soja* | *Synaptobrevin signature* |
| TRAES3BF073600100CFD_t1 | *Triticum aestivum* | *Syntaxin / epimorphin family signature* |
| PH01002656G0020 | *Phyllostachys heterocycla* | *TonB-dependent receptor proteins signature 1* |
| AA11G00017 | *Aethionema arabicum* | *TonB-dependent receptor proteins signature 1* |
| AA32G01100 | *Aethionema arabicum* | *TonB-dependent receptor proteins signature 1* |
| PH01002656G0020 | *Phyllostachys heterocycla* | *TonB-dependent receptor proteins signature 1* |
| XP_010520901.1 | *Tarenaya hassleriana* | *TonB-dependent receptor proteins signature 1* |
| XP_010539252.1 | *Tarenaya hassleriana* | *TonB-dependent receptor proteins signature 1* |
| C.cajan_41044 | *Cajanus cajan* | *TonB-dependent receptor proteins signature 1* |
| Zosma1g02650 | *Zostera marina* | *Translationally controlled tumor protein (TCTP) domain signature 2* |
| Spipo0G0093000 | *Spirodela polyrhiza* | *Trp-Asp (WD) repeats signature* |
| Tp4g25130 | *Thellungiella parvula* | *Trp-Asp (WD) repeats signature* |
| 483400 | *Arabidopsis lyrata* | *Trp-Asp (WD) repeats signature* |
| Bostr.25993s0479.1.p | *Boechera stricta* | *Trp-Asp (WD) repeats signature* |
| GSBRNA2T00045062001 | *Brassica napus* | *Trp-Asp (WD) repeats signature* |
| GSBRNA2T00084755001 | *Brassica napus* | *Trp-Asp (WD) repeats signature* |
| XP_013634618.1 | *Brassica oleracea* | *Trp-Asp (WD) repeats signature* |
| XP_013634619.1 | *Brassica oleracea* | *Trp-Asp (WD) repeats signature* |
| .XP_013634620.1 | *Brassica oleracea* | *Trp-Asp (WD) repeats signature* |
| XP_009142151.1 | *Brassica rapa* | *Trp-Asp (WD) repeats signature* |
| XP_008221120.1 | *Prunus mume* | *Trp-Asp (WD) repeats signature* |
| KFK37142.1 | *Arabis alpina* | *Trp-Asp (WD) repeats signature* |

| | | |
|---|---|---|
| Kalax-0139s0008.1.p | *Kalanchoe marnieriana* | *Tubulin subunits alpha, beta, and gamma signature* |
| Kalax-0355s0005.1.p | *Kalanchoe marnieriana* | *Tubulin subunits alpha, beta, and gamma signature* |
| XP_016502907.1 | *Nicotiana tabacum* | *Tubulin-beta mRNA autoregulation signal* |
| Zmw_sc03169.1.g00010.1 | *Zoysia matrella* | *Tubulin-beta mRNA autoregulation signal* |
| GSMUA_Achr7P09510_001 | *Musa acuminata* | *Zinc carboxypeptidases, zinc-binding region 2 signature* |
| Pavir.Ab01115 | *Panicum virgatum* | *Zinc carboxypeptidases, zinc-binding region 2 signature* |
| Pavir.J12496 | *Panicum virgatum* | *Zinc carboxypeptidases, zinc-binding region 2 signature* |
| PDK_30s745071g005 | *Phoenix dactylifera* | *Zinc carboxypeptidases, zinc-binding region 2 signature* |
| MLOC_75795.1 | *Hordeum vulgare* | *Zinc carboxypeptidases, zinc-binding region 2 signature* |
| Pavir-1KG158700.1.p | *Panicum virgatum* | *Zinc carboxypeptidases, zinc-binding region 2 signature* |
| Pavir-1KG168100.1.p | *Panicum virgatum* | *Zinc carboxypeptidases, zinc-binding region 2 signature* |
| PDK_30s745071g005 | *Phoenix dactylifera* | *Zinc carboxypeptidases, zinc-binding region 2 signature* |
| Traes_3AS_74E386CE7.1 | *Triticum aestivum* | *Zinc carboxypeptidases, zinc-binding region 2 signature* |
| TRAES3BF096900070CFD_t1 | *Triticum aestivum* | *Zinc carboxypeptidases, zinc-binding region 2 signature* |
| Do016529.1 | *Dichanthelium oligosanthes* | *Zinc finger BED-type profile* |
| Araip.9MG9F | *Arachis ipaensis* | *Zinc finger C2H2 type domain signature* |
| PDK_30s663981g010 | *Phoenix dactylifera* | *Zinc-containing alcohol dehydrogenases signature* |
| PDK_30s663981g010 | *Phoenix dactylifera* | *Zinc-containing alcohol dehydrogenases signature* |
| Araha.61839s0001.1.p | *Arabidopsis halleri* | |
| OB11G21460.1 | *Oryza brachyantha* | |
