## Supplementary Table 3 for "Genomics, Molecular and Evolutionary Perspective of NAC Transcription Factors"

NAC TFs showing their involvement in different pathways and biological process.

| Pathways of NAC TFs | Gene Ontology identifier | False discovery rate |
|---|---|---|
| Abscisic acid-activated signaling pathway | GO.0009738 | 0.042 |
| Aerenchyma formation | GO.0010618 | 0.00901 |
| Aging | GO.0007568 | 8.89E-06 |
| Anatomical structure development | GO.0048856 | 7.77E-21 |
| Anatomical structure formation involved in morphogenesis | GO.0048646 | 0.00169 |
| Anatomical structure morphogenesis | GO.0009653 | 1.23E-16 |
| Androecium development | GO.0048466 | 0.0436 |
| Anthocyanin accumulation in tissues in response to UV light | GO.0043481 | 0.0239 |
| Anthocyanin-containing compound biosynthetic process | GO.0009718 | 0.00442 |
| Aromatic compound biosynthetic process | GO.0019438 | 1.44E-35 |
| Asymmetric cell division | GO.0008356 | 0.0265 |
| Auxin influx | GO.0060919 | 1.46E-06 |
| Auxin polar transport | GO.0009926 | 0.0357 |
| Auxin transport | GO.0060918 | 0.0128 |
| Auxin-activated signaling pathway | GO.0009734 | 0.00338 |
| Biological regulation | GO.0065007 | 7.85E-33 |
| Biological_process | GO.0008150 | 0.000629 |
| Biosynthetic process | GO.0009058 | 1.44E-16 |
| Cell communication | GO.0007154 | 1.07E-11 |
| Cell development | GO.0048468 | 2.09E-09 |
| Cell differentiation | GO.0030154 | 1.97E-12 |
| Cell fate commitment | GO.0045165 | 0.015 |
| Cell growth | GO.0016049 | 0.000551 |
| Cell morphogenesis | GO.0000902 | 0.000244 |
| Cell morphogenesis involved in differentiation | GO.0000904 | 0.000137 |
| Cell tip growth | GO.0009932 | 0.0014 |
| Cell wall organization | GO.0071555 | 0.00345 |
| Cell wall organization or biogenesis | GO.0071554 | 8.29E-06 |
| Cellular aromatic compound metabolic process | GO.0006725 | 1.46E-22 |
| Cellular biosynthetic process | GO.0044249 | 2.58E-18 |
| Cellular component organization | GO.0016043 | 0.021 |
| Cellular component organization or biogenesis | GO.0071840 | 0.000535 |
| Cellular developmental process | GO.0048869 | 1.19E-10 |
| Cellular heat acclimation | GO.0070370 | 0.00717 |
| Cellular macromolecule biosynthetic process | GO.0034645 | 9.31E-34 |
| Cellular macromolecule metabolic process | GO.0044260 | 4.50E-16 |
| Cellular metabolic process | GO.0044237 | 5.22E-06 |
| Cellular nitrogen compound biosynthetic process | GO.0044271 | 7.48E-28 |
| Cellular nitrogen compound metabolic process | GO.0034641 | 5.23E-18 |
| Cellular process | GO.0009987 | 1.86E-07 |
| Cellular response to acid chemical | GO.0071229 | 2.33E-07 |
| Cellular response to auxin stimulus | GO.0071365 | 0.00188 |
| Cellular response to chemical stimulus | GO.0070887 | 9.87E-07 |
| Cellular response to gibberellin stimulus | GO.0071370 | 3.53E-06 |
| Cellular response to hormone stimulus | GO.0032870 | 2.11E-06 |
| Cellular response to jasmonic acid stimulus | GO.0071395 | 0.000857 |

| | | |
|---|---|---|
| Cellular response to lipid | GO.0071396 | 2.97E-05 |
| Cellular response to organic substance | GO.0071310 | 2.15E-07 |
| Cellular response to oxygen-containing compound | GO.1901701 | 2.66E-06 |
| Cellular response to stimulus | GO.0051716 | 3.41E-14 |
| Cellular response to stress | GO.0033554 | 3.01E-07 |
| Cellular response to sulfur starvation | GO.0010438 | 0.00231 |
| Defense response | GO.0006952 | 1.78E-13 |
| Defense response to bacterium | GO.0042742 | 0.00012 |
| Defense response to bacterium, incompatible interaction | GO.0009816 | 0.00141 |
| Defense response to fungus, incompatible interaction | GO.0009817 | 0.0192 |
| Defense response to other organism | GO.0098542 | 0.000304 |
| Defense response, incompatible interaction | GO.0009814 | 3.74E-06 |
| Detection of bacterium | GO.0016045 | 0.0239 |
| Developmental cell growth | GO.0048588 | 0.00412 |
| Developmental growth | GO.0048589 | 0.00171 |
| Developmental growth involved in morphogenesis | GO.0060560 | 0.00161 |
| Developmental maturation | GO.0021700 | 6.83E-06 |
| Developmental process | GO.0032502 | 5.95E-23 |
| Developmental process involved in reproduction | GO.0003006 | 5.00E-05 |
| Embryo development ending in seed dormancy | GO.0009793 | 0.00221 |
| Embryonic meristem development | GO.0048508 | 2.69E-08 |
| Embryonic meristem initiation | GO.0090421 | 3.54E-10 |
| ERAD pathway | GO.0036503 | 0.0138 |
| ER-associated ubiquitin-dependent protein catabolic process | GO.0030433 | 0.0138 |
| Floral organ development | GO.0048437 | 0.00856 |
| Floral whorl development | GO.0048438 | 0.00115 |
| Flower development | GO.0009908 | 0.0245 |
| Formation of organ boundary | GO.0010160 | 0.000179 |
| Fruit development | GO.0010154 | 0.000277 |
| Gene expression | GO.0010467 | 1.59E-28 |
| Gibberellic acid mediated signaling pathway | GO.0009740 | 2.10E-06 |
| Gravitropism | GO.0009630 | 0.0182 |
| Heat acclimation | GO.0010286 | 0.015 |
| Heterocycle metabolic process | GO.0046483 | 1.46E-22 |
| Hormone-mediated signaling pathway | GO.0009755 | 1.34E-06 |
| Immune system process | GO.0002376 | 7.48E-08 |
| Indole glucosinolate biosynthetic process | GO.0009759 | 0.0239 |
| Innate immune response | GO.0045087 | 5.60E-08 |
| Integument development | GO.0080060 | 0.00717 |
| Intercellular transport | GO.0010496 | 0.0427 |
| Jasmonic acid mediated signaling pathway | GO.0009867 | 0.000531 |
| Lateral root development | GO.0048527 | 6.03E-06 |
| Lateral root morphogenesis | GO.0010102 | 0.00115 |
| Leaf development | GO.0048366 | 4.16E-05 |
| Leaf senescence | GO.0010150 | 4.15E-06 |
| Macromolecule metabolic process | GO.0043170 | 3.11E-13 |
| Meristem development | GO.0048507 | 5.08E-06 |
| Meristem initiation | GO.0010014 | 7.99E-10 |
| Meristem maintenance | GO.0010073 | 0.0474 |
| Meristem structural organization | GO.0009933 | 9.68E-08 |
| Metabolic process | GO.0008152 | 0.00882 |

| | | |
|---|---|---|
| Multicellular organismal development | GO.0007275 | 3.66E-20 |
| Multicellular organismal process | GO.0032501 | 2.78E-21 |
| Multi-organism process | GO.0051704 | 2.29E-07 |
| Negative regulation of biological process | GO.0048519 | 2.74E-08 |
| Negative regulation of cell communication | GO.0010648 | 2.40E-05 |
| Negative regulation of cell differentiation | GO.0045596 | 0.00254 |
| Negative regulation of cellular biosynthetic process | GO.0031327 | 8.59E-05 |
| Negative regulation of cellular macromolecule biosynthetic process | GO.2000113 | 0.00017 |
| Negative regulation of cellular metabolic process | GO.0031324 | 0.000298 |
| Negative regulation of cellular process | GO.0048523 | 9.51E-10 |
| Negative regulation of developmental process | GO.0051093 | 0.000857 |
| Negative regulation of gene expression | GO.0010629 | 0.00156 |
| Negative regulation of gibberellic acid mediated signaling pathway | GO.0009938 | 0.0107 |
| Negative regulation of macromolecule metabolic process | GO.0010605 | 0.00221 |
| Negative regulation of metabolic process | GO.0009892 | 0.00221 |
| Negative regulation of multicellular organismal process | GO.0051241 | 0.00882 |
| Negative regulation of nitrogen compound metabolic process | GO.0051172 | 2.00E-05 |
| Negative regulation of nucleobase-containing compound metabolic process | GO.0045934 | 0.000157 |
| Negative regulation of response to stimulus | GO.0048585 | 0.00345 |
| Negative regulation of signal transduction | GO.0009968 | 0.00373 |
| Negative regulation of signaling | GO.0023057 | 2.40E-05 |
| Negative regulation of transcription, DNA-templated | GO.0045892 | 0.000231 |
| Negative regulation of trichome patterning | GO.1900033 | 0.000278 |
| Nitrogen compound metabolic process | GO.0006807 | 3.32E-15 |
| Nucleic acid metabolic process | GO.0090304 | 4.93E-31 |
| Nucleobase-containing compound metabolic process | GO.0006139 | 1.92E-25 |
| Organ development | GO.0048513 | 1.76E-23 |
| Organ formation | GO.0048645 | 0.000144 |
| Organ morphogenesis | GO.0009887 | 0.00116 |
| Organic cyclic compound biosynthetic process | GO.1901362 | 1.31E-33 |
| Organic cyclic compound metabolic process | GO.1901360 | 1.51E-21 |
| Organic substance biosynthetic process | GO.1901576 | 8.42E-18 |
| Organic substance metabolic process | GO.0071704 | 0.00021 |
| Pattern specification process | GO.0007389 | 4.24E-08 |
| Phyllome development | GO.0048827 | 1.12E-08 |
| Plant epidermis development | GO.0090558 | 1.76E-07 |
| Plant epidermis morphogenesis | GO.0090626 | 0.0177 |
| Plant-type cell wall biogenesis | GO.0009832 | 1.71E-06 |
| Plant-type hypersensitive response | GO.0009626 | 0.00901 |
| Plant-type secondary cell wall biogenesis | GO.0009834 | 2.75E-09 |
| Positive gravitropism | GO.0009958 | 0.0312 |
| Positive regulation of biological process | GO.0048518 | 2.40E-17 |
| Positive regulation of cell death | GO.0010942 | 2.86E-05 |
| Positive regulation of cellular biosynthetic process | GO.0031328 | 2.32E-17 |
| Positive regulation of cellular component biogenesis | GO.0044089 | 0.000106 |
| Positive regulation of cellular metabolic process | GO.0031325 | 2.38E-17 |
| Positive regulation of cellular process | GO.0048522 | 8.92E-23 |
| Positive regulation of defense response | GO.0031349 | 0.0257 |

| | | |
|---|---|---|
| Positive regulation of defense response to bacterium | GO.1900426 | 0.0151 |
| Positive regulation of leaf senescence | GO.1900057 | 0.00439 |
| Positive regulation of macromolecule metabolic process | GO.0010604 | 4.58E-16 |
| Positive regulation of metabolic process | GO.0009893 | 6.63E-13 |
| Positive regulation of nitrogen compound metabolic process | GO.0051173 | 3.21E-18 |
| Positive regulation of nucleobase-containing compound metabolic process | GO.0045935 | 2.35E-18 |
| Positive regulation of programmed cell death | GO.0043068 | 0.0265 |
| Positive regulation of response to stimulus | GO.0048584 | 0.00169 |
| Positive regulation of secondary cell wall biogenesis | GO.1901348 | 7.35E-09 |
| Positive regulation of sequence-specific DNA binding transcription factor activity | GO.0051091 | 8.89E-06 |
| Positive regulation of signal transduction | GO.0009967 | 0.042 |
| Positive regulation of transcription, DNA-templated | GO.0045893 | 1.23E-18 |
| Post-embryonic development | GO.0009791 | 0.000358 |
| Post-embryonic morphogenesis | GO.0009886 | 0.00169 |
| Post-embryonic organ development | GO.0048569 | 6.79E-07 |
| Primary metabolic process | GO.0044238 | 2.86E-05 |
| Primary shoot apical meristem specification | GO.0010072 | 1.02E-08 |
| Programmed cell death | GO.0012501 | 3.38E-06 |
| Regionalization | GO.0003002 | 1.11E-07 |
| Regulation of abscisic acid-activated signaling pathway | GO.0009787 | 0.00901 |
| Regulation of actin filament depolymerization | GO.0030834 | 0.0427 |
| Regulation of biological process | GO.0050789 | 2.14E-40 |
| Regulation of cell communication | GO.0010646 | 9.30E-09 |
| Regulation of cell death | GO.0010941 | 0.0069 |
| Regulation of cell differentiation | GO.0045595 | 1.12E-06 |
| Regulation of cell fate commitment | GO.0010453 | 0.000108 |
| Regulation of cell wall organization or biogenesis | GO.1903338 | 1.87E-10 |
| Regulation of cellular biosynthetic process | GO.0031326 | 1.89E-44 |
| Regulation of cellular component biogenesis | GO.0044087 | 8.91E-06 |
| Regulation of cellular macromolecule biosynthetic process | GO.2000112 | 8.08E-45 |
| Regulation of cellular metabolic process | GO.0031323 | 1.84E-44 |
| Regulation of cellular process | GO.0050794 | 1.00E-44 |
| Regulation of defense response | GO.0031347 | 0.000101 |
| Regulation of defense response to insect | GO.2000068 | 0.0239 |
| Regulation of developmental process | GO.0050793 | 3.37E-06 |
| Regulation of embryonic development | GO.0045995 | 0.0265 |
| Regulation of epidermal cell differentiation | GO.0045604 | 0.0432 |
| Regulation of flavonoid biosynthetic process | GO.0009962 | 0.00442 |
| Regulation of gene expression | GO.0010468 | 1.45E-44 |
| Regulation of gibberellic acid mediated signaling pathway | GO.0009937 | 0.00579 |
| Regulation of glucosinolate biosynthetic process | GO.0010439 | 0.00102 |
| Regulation of leaf development | GO.2000024 | 0.000146 |
| Regulation of leaf senescence | GO.1900055 | 0.00442 |
| Regulation of macromolecule biosynthetic process | GO.0010556 | 4.27E-44 |
| Regulation of macromolecule metabolic process | GO.0060255 | 5.52E-43 |
| Regulation of metabolic process | GO.0019222 | 5.74E-41 |
| Regulation of multicellular organismal development | GO.2000026 | 6.67E-06 |
| Regulation of multicellular organismal process | GO.0051239 | 6.89E-06 |
| Regulation of multi-organism process | GO.0043900 | 0.0177 |

| | | |
|---|---|---|
| Regulation of nitrogen compound metabolic process | GO.0051171 | 7.17E-46 |
| Regulation of nucleic acid-templated transcription | GO.1903506 | 3.26E-46 |
| Regulation of nucleobase-containing compound metabolic process | GO.0019219 | 3.26E-46 |
| Regulation of organ morphogenesis | GO.2000027 | 0.00717 |
| Regulation of primary metabolic process | GO.0080090 | 9.66E-43 |
| Regulation of reactive oxygen species metabolic process | GO.2000377 | 0.0312 |
| Regulation of response to biotic stimulus | GO.0002831 | 0.00626 |
| Regulation of response to external stimulus | GO.0032101 | 0.0109 |
| Regulation of response to stimulus | GO.0048583 | 1.43E-09 |
| Regulation of response to stress | GO.0080134 | 9.34E-05 |
| Regulation of salicylic acid biosynthetic process | GO.0080142 | 0.0432 |
| Regulation of salicylic acid metabolic process | GO.0010337 | 0.0265 |
| Regulation of secondary cell wall biogenesis | GO.2000652 | 6.82E-10 |
| Regulation of secondary metabolite biosynthetic process | GO.1900376 | 0.000119 |
| Regulation of sequence-specific DNA binding transcription factor activity | GO.0051090 | 4.22E-06 |
| Regulation of shoot system development | GO.0048831 | 0.00282 |
| Regulation of signal transduction | GO.0009966 | 6.42E-07 |
| Regulation of signaling | GO.0023051 | 6.28E-09 |
| Regulation of sulfur metabolic process | GO.0042762 | 0.000179 |
| Regulation of transcription, DNA-templated | GO.0006355 | 9.79E-47 |
| Regulation of trichome morphogenesis | GO.2000039 | 0.0239 |
| Reproductive process | GO.0022414 | 4.46E-05 |
| Reproductive shoot system development | GO.0090567 | 0.0139 |
| Reproductive structure development | GO.0048608 | 0.000238 |
| Reproductive system development | GO.0061458 | 0.000238 |
| Response to abiotic stimulus | GO.0009628 | 1.56E-10 |
| Response to abscisic acid | GO.0009737 | 1.03E-05 |
| Response to acid chemical | GO.0001101 | 1.10E-22 |
| Response to auxin | GO.0009733 | 1.38E-07 |
| Response to bacterium | GO.0009617 | 0.000157 |
| Response to chemical | GO.0042221 | 3.74E-17 |
| Response to chitin | GO.0010200 | 9.07E-05 |
| Response to endogenous stimulus | GO.0009719 | 1.23E-16 |
| Response to endoplasmic reticulum stress | GO.0034976 | 0.000173 |
| Response to ethylene | GO.0009723 | 0.00538 |
| Response to external stimulus | GO.0009605 | 7.93E-07 |
| Response to fungus | GO.0009620 | 0.0254 |
| Response to gibberellin | GO.0009739 | 1.56E-15 |
| Response to hormone | GO.0009725 | 7.57E-14 |
| Response to hydrogen peroxide | GO.0042542 | 0.022 |
| Response to inorganic substance | GO.0010035 | 0.00168 |
| Response to insect | GO.0009625 | 0.000146 |
| Response to jasmonic acid | GO.0009753 | 1.16E-17 |
| Response to lipid | GO.0033993 | 2.73E-13 |
| Response to nitrogen compound | GO.1901698 | 0.0184 |
| Response to organic cyclic compound | GO.0014070 | 4.11E-12 |
| Response to organic substance | GO.0010033 | 1.22E-20 |
| Response to osmotic stress | GO.0006970 | 1.12E-06 |
| Response to other organism | GO.0051707 | 1.46E-06 |

| | | |
|---|---|---|
| Response to oxygen-containing compound | GO.1901700 | 1.92E-22 |
| Response to radiation | GO.0009314 | 0.0455 |
| Response to salicylic acid | GO.0009751 | 3.96E-16 |
| Response to salt stress | GO.0009651 | 1.39E-06 |
| Response to stimulus | GO.0050896 | 1.23E-16 |
| Response to stress | GO.0006950 | 1.16E-17 |
| Response to unfolded protein | GO.0006986 | 0.0151 |
| Response to UV | GO.0009411 | 0.0334 |
| Response to water deprivation | GO.0009414 | 0.000207 |
| Response to wounding | GO.0009611 | 0.000108 |
| RNA metabolic process | GO.0016070 | 4.00E-34 |
| Root cap development | GO.0048829 | 1.88E-09 |
| Root development | GO.0048364 | 1.88E-15 |
| Root epidermal cell differentiation | GO.0010053 | 1.15E-05 |
| Root hair cell development | GO.0080147 | 0.000551 |
| Root hair cell differentiation | GO.0048765 | 1.20E-05 |
| Root hair cell tip growth | GO.0048768 | 0.00741 |
| Root hair elongation | GO.0048767 | 0.0048 |
| Root morphogenesis | GO.0010015 | 1.05E-09 |
| Root system development | GO.0022622 | 2.08E-15 |
| Seed development | GO.0048316 | 0.000723 |
| Shoot system development | GO.0048367 | 1.02E-08 |
| Shoot system morphogenesis | GO.0010016 | 0.000919 |
| Sieve element differentiation | GO.0090603 | 0.00901 |
| Sieve element enucleation | GO.0090602 | 0.00901 |
| Signal transduction | GO.0007165 | 1.61E-10 |
| Single organism reproductive process | GO.0044702 | 7.43E-06 |
| Single organism signaling | GO.0044700 | 3.53E-11 |
| Single-organism cellular process | GO.0044763 | 0.0279 |
| Single-organism developmental process | GO.0044767 | 4.17E-23 |
| Single-organism process | GO.0044699 | 0.000756 |
| Specification of symmetry | GO.0009799 | 0.0432 |
| Stamen development | GO.0048443 | 0.0436 |
| Stamen filament development | GO.0080086 | 0.0107 |
| Stem cell differentiation | GO.0048863 | 0.0432 |
| System development | GO.0048731 | 4.40E-16 |
| Tissue development | GO.0009888 | 2.58E-18 |
| Transcription, DNA-templated | GO.0006351 | 8.08E-45 |
| Transport of virus in host, cell to cell | GO.0046740 | 0.00901 |
| Trichoblast differentiation | GO.0010054 | 1.85E-05 |
| Trichome differentiation | GO.0010026 | 5.88E-05 |
| Trichome morphogenesis | GO.0010090 | 0.0109 |
| Trichome patterning | GO.0048629 | 0.000278 |
| Unidimensional cell growth | GO.0009826 | 0.00531 |
| Xylem development | GO.0010089 | 4.16E-06 |
| Xylem vessel member cell differentiation | GO.0048759 | 1.43E-09 |
